# pyMeSHSim: an integrative python package for biomedical named entity recognition, normalization and comparison

**DOI:** 10.1101/459172

**Authors:** Zhi-Hui Luo, Meng-Wei Shi, Zhuang Yang, Hong-Yu Zhang, Zhen-Xia Chen

## Abstract

**Motivation:** Increasing disease causal genes have been identified through different methods, while there are still no uniform biomedical named entity (bio-NE) annotations of the disease phenotypes. Furthermore, semantic similarity comparison between two bio-NE annotations, like disease descriptions, has become important for data integration or system genetics analysis.

**Methods:** The package pyMeSHSim realizes bio-NEs recognition using MetaMap, which produces Unified Medical Language System (UMLS) concepts in natural language process. To map the UMLS concepts to MeSH, pyMeSHSim embedded a house made dataset containing the Medical Subject Headings (MeSH) main headings (MHs), supplementary concept records (SCRs) and relations between them. Based on the dataset, pyMeSHSim implemented four information content (IC) based algorithms and one graph-based algorithm to measure the semantic similarity between two MeSH terms.

**Results:** To evaluate its performance, we used pyMeSHSim to parse OMIM and GWAS phenotypes. The inclusion of SCRs and the curation strategy of non-MeSH-synonymous UMLS concepts used by pyMeSHSim improved the performance of pyMeSHSim in the recognition of OMIM phenotypes. In the curation of GWAS phenotypes, pyMeSHSim and previous manual work recognized the same MeSH terms from 276/461 GWAS phenotypes, and the correlation between their semantic similarity calculated by pyMeSHSim and another semantic analysis tool *meshes* was as high as 0.53-0.97.

**Conclusion:** With the embedded dataset including both MeSH MHs and SCRs, the integrative MeSH tool pyMeSHSim realized the disease recognition, normalization and comparison in biomedical text-mining.

**Availability:** Package’s source code and test datasets are available under the GPLv3 license at https://github.com/luozhhub/pyMeSHSim

## INTRODUCTION

Biomedical Named entity (bio-NE) recognition, normalization and comparison are fundamental and important tasks for extracting and utilizing valuable biomedical knowledge from textual data (1-4). They are realized by identifying key entities in unstructured text, mapping identified entities to a controlled vocabulary, and measuring the semantic similarity between the controlled vocabulary terms (5).

Medical Subject Headings (MeSH) is a controlled vocabulary that can be used in bio-NE recognition, normalization and comparison (6). It consists of three main record types including descriptor records, qualifier records and supplementary concept records (SCRs). MeSH is curated by the National Library of Medicine (NLM) and serves as the index system in PubMed/MEDLINE and other NLM databases. NLM has used Medical Text Indexer (MTI) to provide indexing recommendations based on MeSH in the bio-NE recognition for literatures since 2002 (7). As precise literature annotations, MeSH has become more and more popular in the normalization of bio-NEs, including disease names, in medical and genetic public databases (8, 9). Also, the structure of MeSH as a directed acyclic graph like Gene Ontology (10) and Disease Ontology (11) enables the comparison of semantic similarity between two MeSH terms in the graph.

Several MeSH tools have been developed to realize bio-NE recognition, normalization or comparison. As a MeSH tool for bio-NE recognition and normalization, NLM MeSH has provided a browser online (https://meshb.nlm.nih.gov/search) to parse MeSH terms from the input phrases. However, it is neither tolerant to even subtle difference of input phrases from MeSH terms, nor applicable to batch processing. As MeSH tools for bio-NE comparison, *meshes* (12) and *meshSim* (13) have been developed recently to measure MeSH semantic similarity using the R dataset MeSH.Hsa.eg.db (3) as data framework. However, the lack of SCRs embedded in its MeSH dataset limits the use of both bio-NE comparison tools for rare diseases such as “alzheimer’s disease 7” and “Bardet-Biedl syndrome 11”. Furthermore, there is still a lack of an integrated one-stop MeSH toolkit to realize bio-NE recognition, normalization and comparison.

To solve above problems, an integrative python package pyMeSHSim was developed to realize bio-NE recognition, normalization and comparison. It can directly curate MeSH terms from free biomedical descriptions and measure the semantic similarity between the MeSH term pairs. Additionally, to enable batch processing and the application of pyMeSHSim to both common diseases and rare diseases, a lightweight and comprehensive MeSH dataset, was generated and embedded as the data framework of pyMeSHSim.

## METHODS

### Dataset construction

A comprehensive MeSH dataset is fundamental to MeSH tools, while the MeSH dataset used by most popular MeSH tools contains only MeSH Main Headings (MHs), a component of MeSH descriptor records, but not SCRs. To construct a comprehensive MeSH dataset, we exacted MeSH information, including MHs, SCRs, and their relations, from Unified Medical Language System (UMLS, 2018AA version), which is a large biomedical thesaurus integrated nearly 200 vocabularies including MeSH (14).

The multiple-to-one relationship between MeSH-synonymous UMLS concepts and MeSH MHs was curated from the UMLS table MRSAT. For example, the MeSH MH “Alzheimer Disease” (D000544) includes seven MeSH concepts, each of which corresponds to several MeSH entry terms and a UMLS concept (Supplementary table 1). In our dataset, we included the MeSH MHs and related UMLS concepts, while excluded the MeSH concept and MeSH entry term information. Moreover, we curated the most useful “parent” and “child” relationship between MeSH MHs from the UMLS table MRREL.

**Table 1.**
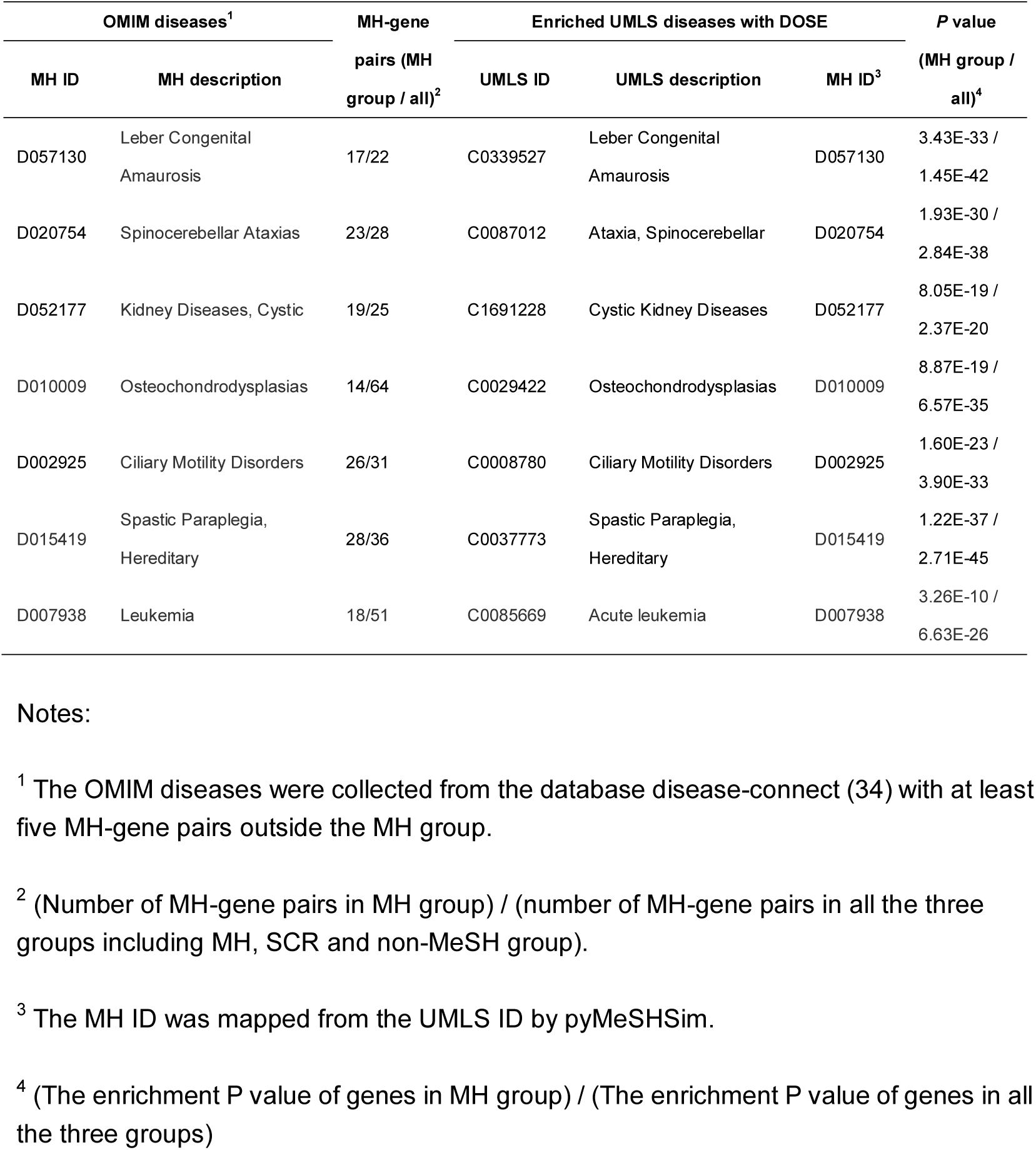
Disease enrichment analysis of the genes assigned to the MHs before and after addition of MH-gene pairs from SCR and non-MeSH groups.

The one-to-one relationship between MeSH-synonymous UMLS concepts and SCRs was curated from the UMLS table MRSAT. We included the SCRs and its corresponding UMLS concepts, as well as the “narrower” and “broader” relationship between SCRs and MeSH MHs curated from the UMLS table MRREL, in our dataset.

The qualifier records and other MeSH descriptor records except MeSH MHs were not included in our dataset. We used “MeSH term” in the following to refer MeSH MH or SCR.

### Bio-NE recognition and normalization

The bio-NE recognition were realized by MetaMap (15), a widely used biomedical natural language processing software that recognizes UMLS concepts from free texts. Although machine learning methods might have better performance than MetaMap in recommending MeSH MHs to MEDLINE citations, their use were constrained by the requirement of large amount of training data to establish the model, and the potential imbalance of the training data (16). Meanwhile, the bio-NEs, such as disease phenotypes from GWASdb (17), OMIM (18), and GAD (19), and drug indications from public databases DrugBank (20) and TTD (21), might have to be curated from free texts without large amount of training data, which is the expertise of MetaMap. The UMLS concepts curated by MetaMap were then converted to MeSH terms based on our dataset. MeSH-synonymous UMLS concepts were mapped to MHs or SCRs, and non-MeSH-synonymous UMLS concepts were curated as free texts and mapped to MHs or SCRs.

### Bio-NE comparison

We compared the bio-NEs based on the similarity between their corresponding MeSH terms. The semantic similarity is usually calculated by graph-based or information content (IC)-based measures. The graph-based method measures the path distance between two MeSH terms on the MeSH hierarchical structure, while the IC-based method depends on the specificity and informativeness of MeSH terms (22).

We retrieved the number of publications indexed by MeSH terms using the NCBI E-Utility (23), and calculated the IC vaules as below.

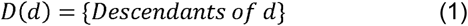

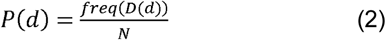

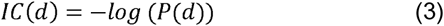

Where *D(d)* is the descendent terms of MeSH term *d*; *freq(x)* is the number of publications indexed by term *x*; *N* is the total number of publications indexed by MeSH; and *IC(d)* is the IC value of term *d*.

We implemented the following four IC-based algorithms:

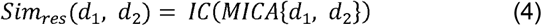

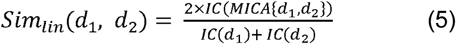

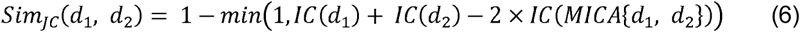

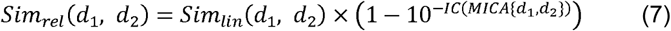

Where *d*_1_ and *d*_2_ are MeSH terms; *Sim*_*lin*_, *Sim*_*res*_, *Sim*_*rel*_, *Sim*_*JC*_ corresponds to Lin’s (24), Resnik’s (25), Schlicker’s (26), and Jiang and Conrath’s (27) algorithms, respectively; MICA (the most informative common ancestor) is the ancestor of the selected two MeSH terms with the maximal IC value among all ancestors. We assigned MICA between MeSH terms from different categories, denoted by the first character of the tree number of MeSH terms, to 0. For example, MICA between the MeSH terms “Tauopathies” (tree number: “C10.574.945”) and “Schizophrenia” (tree number: “F03.700.750”) is 0 because they belong to different categories (“C” for diseases vs “F” for psychiatry and psychology).

We also implemented the graph based Wang’s (28) algorithm as below.

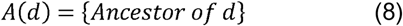

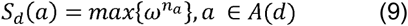

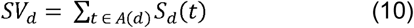

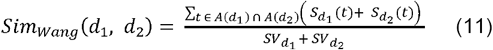

Where *d* is a MeSH term; *A(d)* is the ancestors, deduced from tree numbers, of *d*; *ω*, which is set to 0.7 in our work, is the weight used to measure the relation between two terms; *n*_*a*_ is the number of edges between *d* to *a*; *S*_*d*_(*a*) is the semantic contribution of *a* to *d*; *SV*_*d*_ is the total semantic contributions of all ancestors to *d*; *Sim*_*wang*_(*d*_1_, *d*_2_) is Wang’s algorithm score between MeSH terms *d*_1_ and *d*_2_.

Noteworthily, both IC-based and graph-based methods depend on the tree numbers, which may be not unique, of the MeSH terms, and thus lead to multiple similarity values. We retained only the maximal similarity value between two MeSH terms.

## IMPLEMENTATION

The pyMeSHSim consists of three subpackages (1) the *metamapWrap* subpackage that recognizes bio-NEs from the text, and converts bio-NEs into MeSH terms in bio-NE recognition (2) the *data* subpackage translates UMLS concepts to MeSH terms, by the embedded MeSH dataset in bio-NE normalization, and (3) the *Sim* subpackage that measures the distance between MeSH terms in the bio-NE comparison (Figure 1).

**Figure 1.**
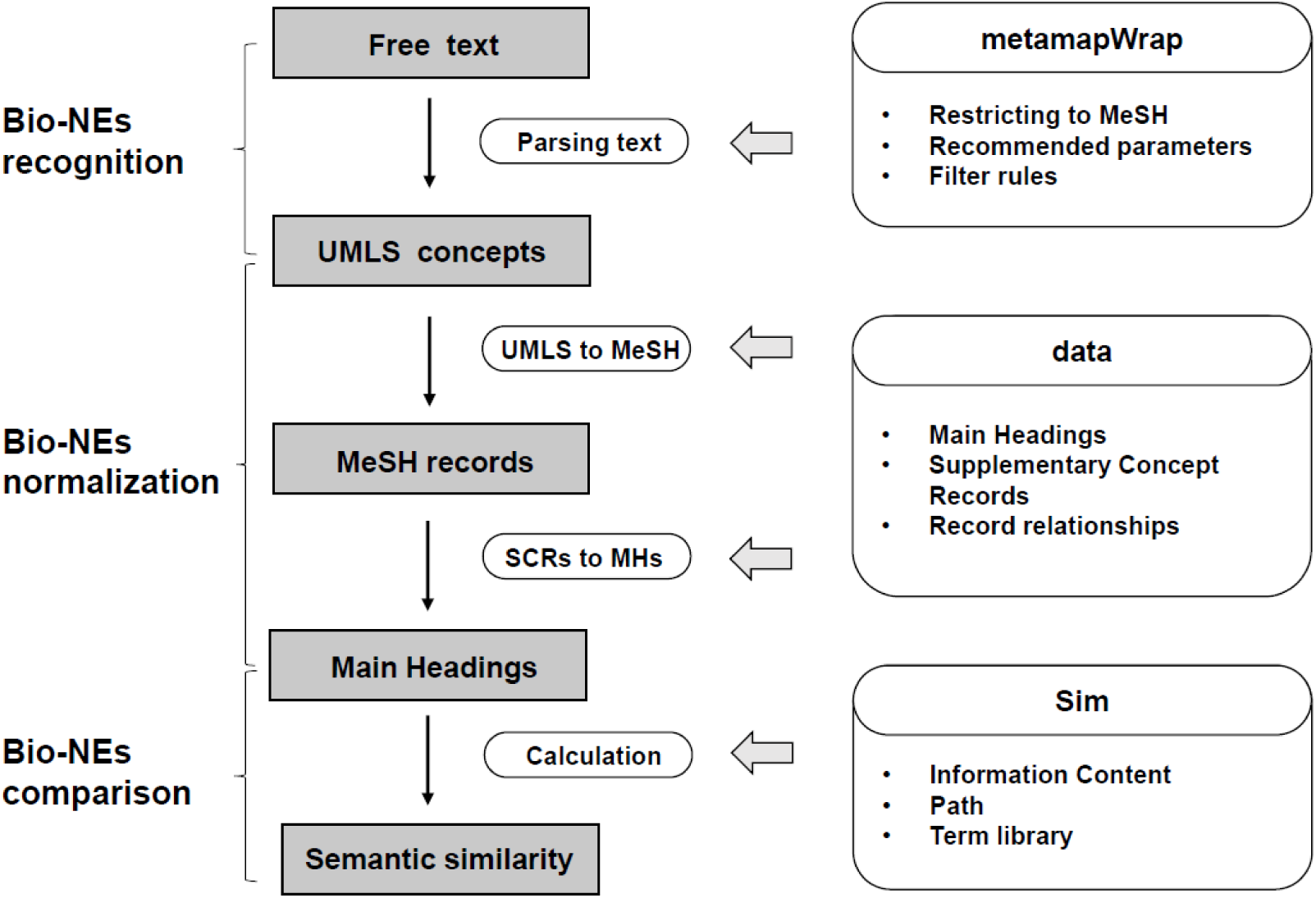
The components and workflow of pyMeSHSim. pyMeSHSim consists of three subpackages, including *metamapWrap, data* and *Sim*. In bio-NE recognition, *metamapWarp* curates the UMLS concepts from free text. In bio-NE normalization, *data* translates UMLS concepts to MeSH terms, and maps SCRs to MHs using selected records and relationships between records in MeSH. In bio-NEs comparison, *Sim* uses IC-based and graph-based methods to measure semantic similarity between two bio-NEs.

### The metamapWrap subpackage

The bio-NE recognition and normalization of pyMeSHSim were realized by the *metamapWrap* subpackage, which was a wrapper for MetaMap (15). The subpackage *metamapWrap* curated MeSH-synonymous UMLS concepts from free texts, including non-MeSH-synonymous UMLS concepts, and then converted UMLS concepts to corresponding MeSH terms via the data subpackage. We set parameters “-N –J list_semantic_type –R MSH –I –z –conj –Q 4 –silent --sldi”, where list_semantic_type was the list of disease-related semantic types “inpo,dsyn,phpranab,orgf,clna,hlca,genf,orga,neop,emod,inbe,lbtr,anst,npop,celc,cell,bpoc,act y,mobd,celf,evnt,sosy,patf,tisu,moft,fndg,bdsu,ortf,menp,acab,comd,sbst,cgab”, as default of pyMeSHSim (see manuals at http://pymeshsim.systemsgenetics.cn/). Users can tune a series of parameters to get more accurate results.

### The data subpackage

The MeSH dataset was embedded in the *data* subpackage in bcolz format with a corresponding data interface (Supplementary table 2). It included five tables: (1) table “MainHeadingDetailData” stored all the MH information, including MeSH unique id, tree code, prefer name, category, term semantic type, IC frequency, and UMLS id. The semantic type was curated from the UMLS table MRSTY, and each UMLS concept was characterized by at least one of the 133 semantic types (29). (2) table “supplementMainHeading” stored all the UMLS concepts related to MHs; (3) table “RNDetailData” stored the basic information of SCRs; (4) table “RNandRBRel” stored the narrower-and-broader relationship between SCRs and MHs; and (5) table “ParentChildRel” stored the fundamental tree structure. The five tables enabled the conversion of UMLS concepts to MeSH terms, and the measurement of the semantic similarity between MeSH terms.

**Table 2.** Correlation of calculated semantic similarities between pyMeSHSim and *meshes*.

| **Method** | **Lin's** | **Res'** | **Jiang's** | **Rel's** | **Wang's** |
| --- | --- | --- | --- | --- | --- |
| **Correlation coefficient** | 0.97 | 0.53 | 0.96 | 0.97 | 0.84 |
| ***P* value** | <2.2e-16 | 1.8e-03 | <2.2e-16 | <2.2e-16 | 1.7e-11 |

### The Sim subpackage

The bio-NE comparison of pyMeSHSim was realized by the *Sim* subpackage via measuring the distance between MeSH terms. Each narrower record from the SCRs was converted to one or more broader terms from MHs before the measurement. As the tool *meshes*, pyMeSHSim offered five representative methods of semantic similarity measures, including four information content (IC) based (Lin’s, Resnik’s, Schlicker’s, and Jiang and Conrath’s) and one graph-based (Wang’s) algorithms.

## PERFORMANCE

### Evaluation with OMIM phenotypes

To test whether the involvement of SCRs and our curation strategy of non-MeSH-synonymous UMLS concepts would contribute to improved performance of pyMeSHSim in bio-NE recognition, we compared the genes annotated with MeSH MHs and SCRs from OMIM (18) phenotype-gene pairs. The OMIM phenotype-gene pairs were collected from the database disease-connect (30), which used MetaMap to process the disease phenotypes into MeSH-synonymous and non-MeSH-synonymous UMLS concepts. Using pyMeSHSim, MeSH-synonymous UMLS concepts were mapped to MHs and SCRs, which were further mapped to their “broader” MHs, and non-MeSH-synonymous UMLS concepts were curated as free texts and mapped to MHs. We classified OMIM phenotypes into MH, SCR, and non-MeSH groups based on the source of their corresponding UMLS concepts, and compared genes corresponded to the same MHs from all the three groups (Figure 2). The genes without Entrez IDs, which were required for the following disease enrichment analysis, and the MHs with less than 10 genes in two or three groups were excluded from subsequently analysis. After the filtering, 36 MHs and 1498 MH-gene pairs (Supplementary table 3) were remained, including 761 MH-gene pairs from MH group, 552 from SCR group and 215 from non-MeSH group. About 87.5% MH-gene pairs in SCR group also presented in MH group, demonstrating high overlap of genetic features between subtype diseases and its corresponding MH diseases, and validating the significance of SCRs in disease curation (Figure 3). Additionally, 59.5% and 10.6% MH-gene pairs in non-MeSH group presented in MH and SCR group, respectively, indicating the effectiveness of our curation strategy of non-MeSH-synonymous UMLS concepts.

**Figure 2.**
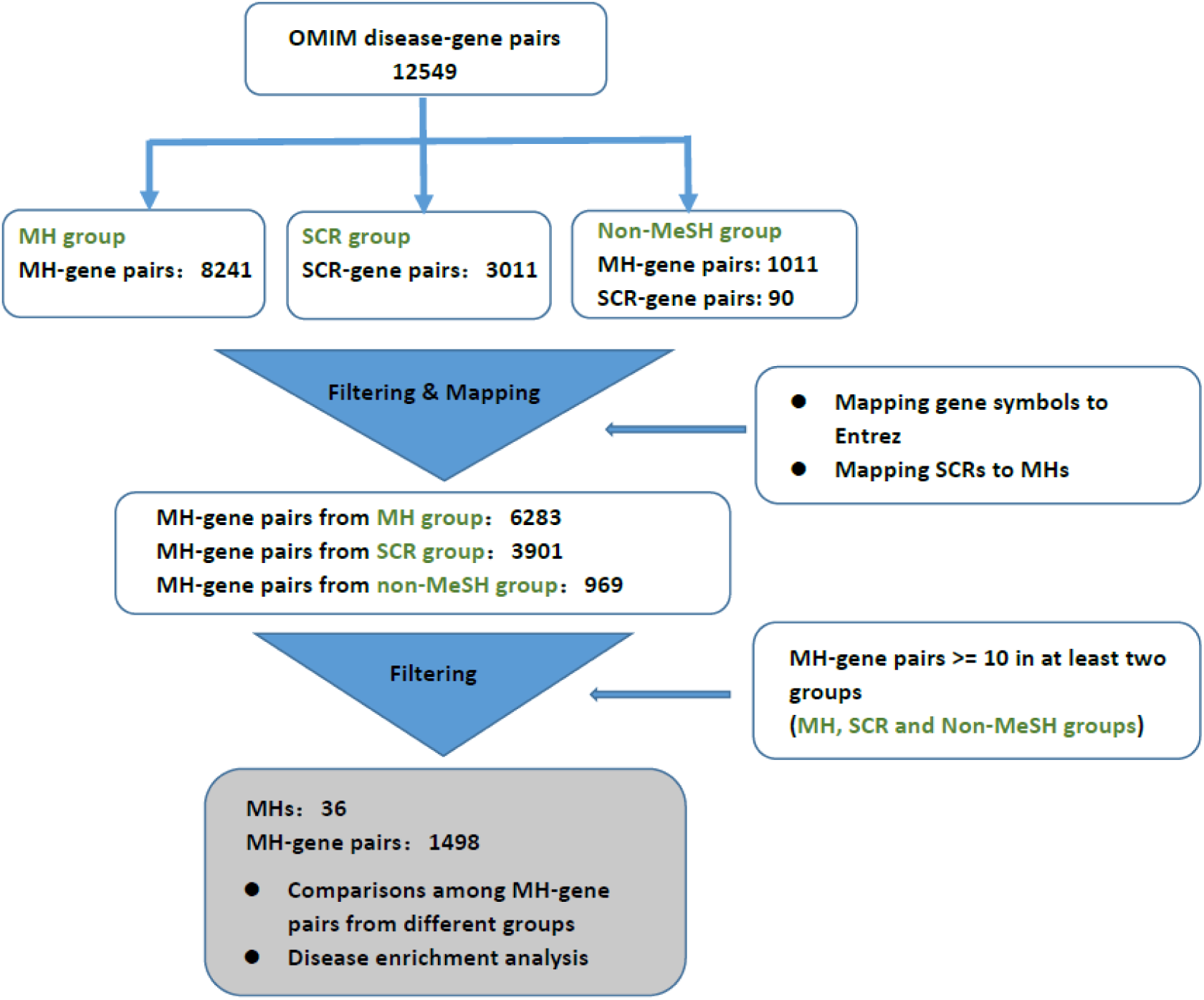
OMIM UMLS diseases processing pipeline. MeSH-synonymous UMLS concepts were mapped to MHs or SCRs by pyMeSHSim directly. Meanwhile, non-MeSH-synonymous UMLS concepts were processed as free texts into MeSH-synonymous UMLS concepts, and then mapped to MeSH terms. All gene symbols were mapped to Entrez IDs. SCRs were mapped to its broader MHs. MHs with at least 10 genes in at least two groups were remained for further analysis.

**Figure 3.**
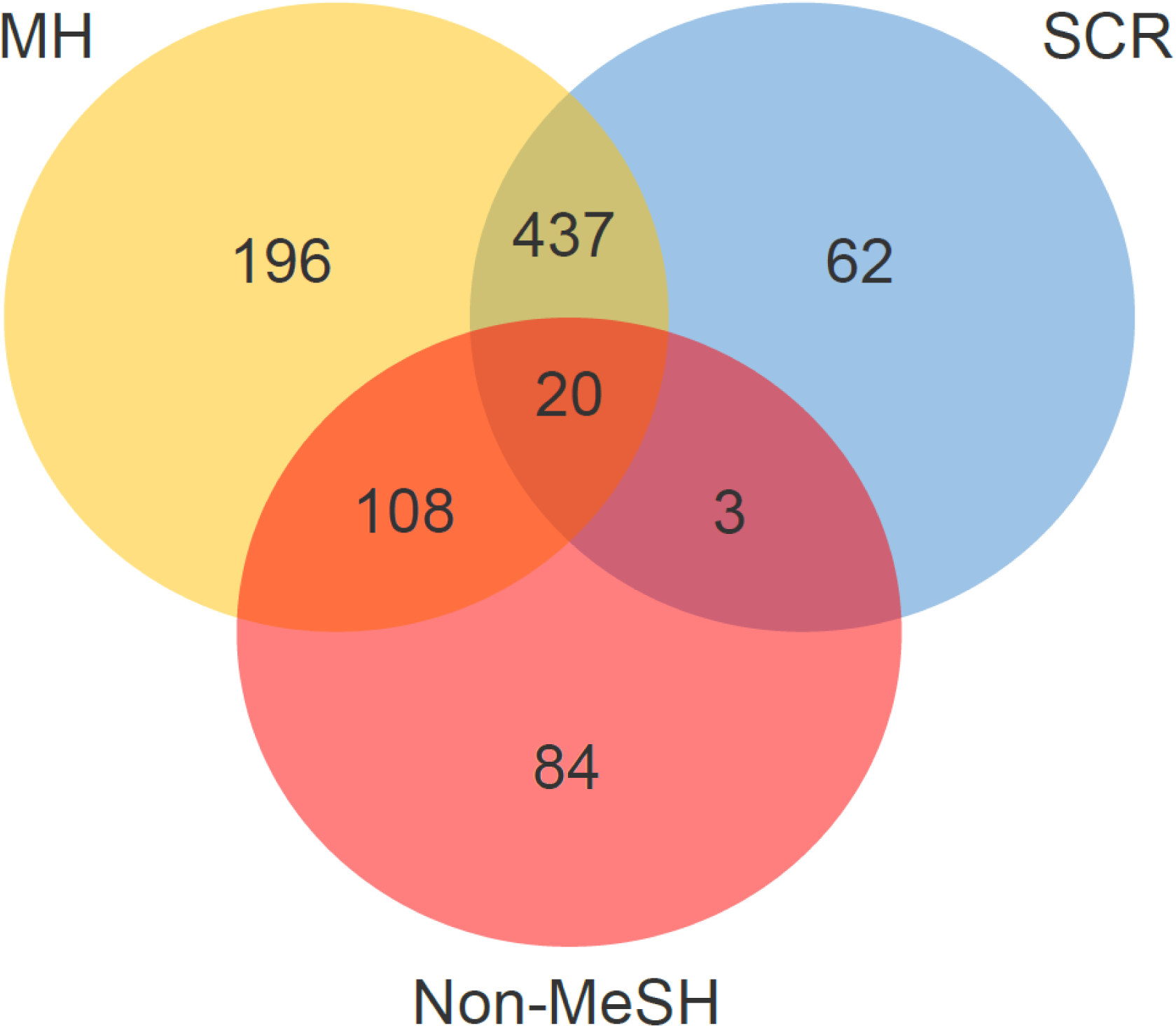
Venn diagram of MH-gene pairs in MH, SCR and Non-MeSH groups.

To further validate the reasonability of SCR involvement and our curation strategy of non-MeSH-synonymous UMLS concepts, we explored whether the additional MH-gene pairs deduced from SCRs and non-MeSH-synonymous UMLS concepts were enriched in the MH diseases. We remained the seven MHs with at least five MH-gene pairs outside the MH group, and tested their enriched diseases using the UMLS-based disease enrichment analysis tool DOSE (31). For each of the seven MHs, the addition of genes from SCR and non-MeSH groups led to more significant enrichment in the disease mapped to the MH by pyMeSHSim (Table 1). Especially, the addition of 50 genes from SCR and non-MeSH groups to the 14 genes from MH group, of the MeSH MH “Osteochondrodysplasias (D010009)” led to the higher pvalue (6.57E-35 vs 8.87E-19) of enrichment in the disease “Osteochondrodysplasias” (Table 1). The more significant enrichment indicated the contribution of SCRs in our dataset and our curation strategy of non-MeSH-synonymous UMLS concepts to the improved performance of pyMeSHSim in bio-NE recognition and normalization.

### Evaluation with GWAS phenotypes

We compared the performance of pyMeSHSim in the recognition and normalization of bio-NEs from GWAS phenotypes with the manual results of Nelson’s group using MeSH browser (4), and compared their semantic similarity calculated by pyMeSHSim and *meshes* (12).

We firstly used pyMeSHSim to parse the 461 GWAS phenotypes collected by Nelson’s group to MeSH terms. PyMeSHSim successfully curated MeSH terms from 443 (96%) GWAS phenotypes. The curated MeSH terms from 310 (67%) GWAS phenotypes had Lin score > 0.75 with the manual work, demonstrating high performance of pyMeSHSim in bio-NE recognition and normalization (Supplementary table 4).

We then calculated the overall semantic similarity between the curated MeSH terms using two methods by pyMeSHSim and the latest semantic analysis tool *meshes* (Supplementary table 4). The similarity calculated using both packages was 1 between the same terms, and 0 between terms in different categories. For the comparison between the different term pairs of 53 GWAS phenotypes in the same category with Nelson’s group, pyMeSHSim succeeded in calculating the similarities between the term pairs of all the phenotypes, while *meshes*, which only compared MH-MH pairs, failed for SCR-MH pairs of 15 phenotypes (Supplementary table 4). Among the 15 SCRs parsed by pyMeSHSim, 13 were mapped to the same MHs as parsed by Nelson’s group. The similarity correlation of the remaining 38 term pairs between pyMeSHSim and *meshes* was 0.53 (Res’)-0.97 (Lin’s and Rel’s) (Table 2), demonstrating similar, if not better, performance of pyMeSHSim compared with *meshes* in bio-NE comparison.

Comparing the different terms of the 114 GWAS phenotypes curated by pyMeSHSim and the Nelson’s group, we found that the latter preferred mapping the phenotypes to disease category (C). For example, “Color, Eye” was returned by MeSH browser with GWAS phenotype “Eye color” as the input. However, “Eye color” was curated as “color vision defects” by Nelson’s group, while as “Color, Eye” by pyMeSHSim. Similarly, GWAS phenotypes “Hair color” and “Serum urate” were curated as “hair diseases” and “urinary calculi” by Nelson’s group, while as “Color, Hair” and “Acid, Uric” by pyMeSHSim (Supplementary table 4). Therefore, at least a part of the differences between the manual work and pyMeSHSim were from the human bias in the manual work.

In summary, we used OMIM phenotypes and GWAS phenotypes to evaluate pyMeSHSim performance in bio-NE recognition, normalization and comparison. Both evaluations illustrate its high effectiveness and precision in bio-NEs recognition and normalization. GWAS analysis also demonstrated its power in bio-NE comparison. Moreover, OMIM analysis validated its effectiveness in the integration of genetic information.

## DISCUSSION

We developed pyMeSHSim, an integrative, lightweight and data-rich python package for biomedical text mining. To the best of our knowledge, this is the first universal MeSH toolkit for integrated bio-NE recognition, normalization and comparison analysis. PyMeSHSim is expected to be widely used as a powerful tool in bioinformatics, computational biology and biomedical research.

Considering that MeSH is one of the most widely used biomedical vocabulary, pyMeSHSim will further contribute to data integration. Also, the inclusion of SCRs in the implemented dataset enabled pyMeSHSim to handle rare diseases in public databases like OMIM and Orphanet (www.orpha.net). However, whether general concepts like MHs or more specific concepts like SCRs are preferable will depend on the end use. Users should be cautious to select the right terms accordingly in using pyMeSHSim.

## AUTHOR CONTRIBUTIONS

Z.C. and H.Z. supervised the work. H.Z. and Z.L. conceived the idea. Z.L. and M.S. developed the software. Z.L. and Z.Y. performed the research. Z.L. and Z.C. wrote the manuscript.

## FUNDING

This work was supported by Huazhong Agricultural University Scientific & Technological Self-innovation Foundation [2016RC011]; and the Fundamental Research Funds for the Central Universities [2662018PY021, 2662017PY115].

