## Supplementary table 1 for "pyMeSHSim: an integrative python package for biomedical named entity recognition, normalization and comparison"

**Supplementary table 1.** MeSH terms and UMLS concepts correspond to MeSH MH D000544.

| **UMLS**  **concept ID** | **MeSH** | | | |
| --- | --- | --- | --- | --- |
|  | **Concept ID** | **Entry term ID** | **Entry term** | **Main heading** |
| C0002395 | M0000842 | T001655 | Alzheimer Disease | D000544  (Alzheimer Disease) |
|  |  | T001654 | Alzheimer's Disease |  |
|  |  | T010820 | Dementia, Senile |  |
|  |  | T365964 | Dementia, Alzheimer Type |  |
|  |  | T843727 | Alzheimer-Type Dementia (ATD) |  |
|  |  | T365981 | Alzheimer Type Senile Dementia |  |
|  |  | T366224 | Primary Senile Degenerative Dementia |  |
|  |  | T369933 | Dementia, Primary Senile Degenerative |  |
|  |  | T840887 | Alzheimer Sclerosis |  |
|  |  | T840888 | Alzheimer Syndrome |  |
|  |  | T000915952 | Alzheimer Dementia |  |
|  |  | T365965 | Senile Dementia, Alzheimer Type |  |
| C0546126 | M0333929 | T365966 | Acute Confusional Senile Dementia |  |
|  |  | T366225 | Senile Dementia, Acute Confusional |  |
| C0011265 | M0005798 | T010818 | Dementia, Presenile |  |
| C0494463 | M0333941 | T365982 | Alzheimer Disease, Late Onset |  |
|  |  | T366226 | Late Onset Alzheimer Disease |  |
| C0750900 | M0333961 | T365983 | Alzheimer's Disease, Focal Onset |  |
|  |  | T366227 | Focal Onset Alzheimer's Disease |  |
| C0276496 | M0584546 | T843728 | Familial Alzheimer Disease (FAD) |  |
| C0750901 | M0333931 | T365968 | Alzheimer Disease, Early Onset |  |
|  |  | T369934 | Early Onset Alzheimer Disease |  |
|  |  | T365967 | Presenile Alzheimer Dementia |  |
