## Supplementary table 2 for "pyMeSHSim: an integrative python package for biomedical named entity recognition, normalization and comparison"

**Supplementary table 2.** Number of MHs and SCRs in each MeSH category.

| Category Name | Abbreviations | No. MHs | No. SCRs |
| --- | --- | --- | --- |
| Chemicals and Drugs | D | 9,933 | 239,512 |
| Analytical, Diagnostic and Therapeutic Techniques and Equipment | E | 2,923 | 1,446 |
| Diseases | C | 4,798 | 6,473 |
| Anatomy | A | 1,825 | 1,150 |
| Organisms | B | 3,814 | 486 |
| Psychiatry and Psychology | F | 1,082 | 559 |
| Phenomena and Processes | G | 2,258 | 645 |
| Health Care | N | 1,794 | 3 |
| Anthropology, Education, Sociology and Social Phenomena | I | 640 | 3 |
| Geographicals | Z | 401 | 0 |
| Disciplines and Occupations | H | 418 | 0 |
| Humanities | K | 199 | 0 |
| Information Science | L | 404 | 0 |
| Named Groups | M | 289 | 1 |
| Technology, Industry, Agriculture | J | 581 | 4,731 |
