## Supplementary table 3 for "pyMeSHSim: an integrative python package for biomedical named entity recognition, normalization and comparison"

**Supplementary table 3**. The OMIM MH-gene pairs from MH, SCR, and Non-MesH.groups.

| **MeSH_ID** | **Gene_symbol** | **Entrez_ID** | **Source** |
| --- | --- | --- | --- |
| D003317 | CSAD | 51380 | MH |
| D003317 | KRT12 | 3859 | MH |
| D003317 | TACSTD2 | 4070 | MH |
| D003317 | UBIAD1 | 29914 | MH |
| D003317 | EFEMP1 | 2202 | MH |
| D003317 | CHST6 | 4166 | MH |
| D003317 | ZEB1 | 6935 | MH |
| D003317 | SLC4A11 | 83959 | MH |
| D003317 | AGBL1 | 123624 | MH |
| D003317 | VSX1 | 30813 | MH |
| D003317 | KRT3 | 3850 | MH |
| D003317 | TGFBI | 7045 | MH |
| D003317 | GSN | 2934 | MH |
| D003317 | COL8A2 | 1296 | MH |
| D003317 | CYP4V2 | 285440 | MH |
| D003317 | DCN | 1634 | MH |
| D003317 | CSAD | 51380 | SCR |
| D003317 | DCN | 1634 | SCR |
| D003317 | VSX1 | 30813 | SCR |
| D003317 | TACSTD2 | 4070 | SCR |
| D003317 | UBIAD1 | 29914 | SCR |
| D003317 | ZEB1 | 6935 | SCR |
| D003317 | TGFBI | 7045 | SCR |
| D003317 | GSN | 2934 | SCR |
| D003317 | SLC4A11 | 83959 | SCR |
| D003317 | CYP4V2 | 285440 | SCR |
| D003317 | CDK13 | 8621 | Non-MeSH |
| D002386 | GALK1 | 2584 | MH |
| D002386 | OPA3 | 80207 | MH |
| D002386 | EPHA2 | 1969 | MH |
| D002386 | SRD5A3 | 79644 | MH |
| D002386 | RAB3GAP2 | 25782 | MH |
| D002386 | LIM2 | 3982 | MH |
| D002386 | CRYBA4 | 1413 | MH |
| D002386 | GJA3 | 2700 | MH |
| D002386 | C19orf12 | 83636 | MH |
| D002386 | SNX3 | 8724 | MH |
| D002386 | SLC33A1 | 9197 | MH |
| D002386 | AGK | 55750 | MH |
| D002386 | PIK3R4 | 30849 | MH |
| D002386 | GFER | 2671 | MH |
| D002386 | ABHD12 | 26090 | MH |
| D002386 | CTDP1 | 9150 | MH |
| D002386 | NAA10 | 8260 | MH |
| D002386 | BCOR | 54880 | MH |
| D002386 | BFSP1 | 631 | MH |
| D002386 | VIM | 7431 | MH |
| D002386 | MIP | 4284 | MH |
| D002386 | FTL | 2512 | MH |
| D002386 | CHMP4B | 128866 | MH |
| D002386 | CRYGB | 1419 | MH |
| D002386 | CRYAA | 1409 | MH |
| D002386 | BEST1 | 7439 | MH |
| D002386 | PITX3 | 5309 | MH |
| D002386 | BRD4 | 23476 | MH |
| D002386 | ERCC2 | 2068 | MH |
| D002386 | SIX6 | 4990 | MH |
| D002386 | COL11A1 | 1301 | MH |
| D002386 | CRYAB | 1410 | MH |
| D002386 | GJA8 | 2703 | MH |
| D002386 | CRYBA1 | 1411 | MH |
| D002386 | ERCC1 | 2067 | MH |
| D002386 | RAB3GAP1 | 22930 | MH |
| D002386 | HSF4 | 3299 | MH |
| D002386 | TDRD7 | 23424 | MH |
| D002386 | CRYBB3 | 1417 | MH |
| D002386 | NHS | 4810 | MH |
| D002386 | SLC16A12 | 387700 | MH |
| D002386 | RAB18 | 22931 | MH |
| D002386 | ITM2B | 9445 | MH |
| D002386 | FYCO1 | 79443 | MH |
| D002386 | AKR1A1 | 10327 | MH |
| D002386 | CRYBB1 | 1414 | MH |
| D002386 | CCDC180 | 100499483 | MH |
| D002386 | CRYBB2 | 1415 | MH |
| D002386 | PAX6 | 5080 | MH |
| D002386 | CRYGS | 1427 | MH |
| D002386 | BFSP2 | 8419 | MH |
| D002386 | EPG5 | 57724 | MH |
| D002386 | RAB3GAP2 | 25782 | SCR |
| D002386 | SRD5A3 | 79644 | SCR |
| D002386 | C19orf12 | 83636 | SCR |
| D002386 | SNX3 | 8724 | SCR |
| D002386 | AGK | 55750 | SCR |
| D002386 | PIK3R4 | 30849 | SCR |
| D002386 | GFER | 2671 | SCR |
| D002386 | NAA10 | 8260 | SCR |
| D002386 | BCOR | 54880 | SCR |
| D002386 | FTL | 2512 | SCR |
| D002386 | VIM | 7431 | SCR |
| D002386 | CRYGB | 1419 | SCR |
| D002386 | ERCC2 | 2068 | SCR |
| D002386 | COL11A1 | 1301 | SCR |
| D002386 | CRYGC | 1420 | SCR |
| D002386 | CRYAB | 1410 | SCR |
| D002386 | GJA8 | 2703 | SCR |
| D002386 | RAB3GAP1 | 22930 | SCR |
| D002386 | ERCC1 | 2067 | SCR |
| D002386 | NHS | 4810 | SCR |
| D002386 | RAB18 | 22931 | SCR |
| D002386 | SLC16A12 | 387700 | SCR |
| D002386 | ITM2B | 9445 | SCR |
| D002386 | AKR1A1 | 10327 | SCR |
| D002386 | EPG5 | 57724 | SCR |
| D002386 | SLC33A1 | 9197 | Non-MeSH |
| D002386 | CRYGD | 1421 | Non-MeSH |
| D002386 | CTDP1 | 9150 | Non-MeSH |
| D057130 | IMPDH1 | 3614 | MH |
| D057130 | LRAT | 9227 | MH |
| D057130 | GUCY2D | 3000 | MH |
| D057130 | AIPL1 | 23746 | MH |
| D057130 | HSD11B1L | 374875 | MH |
| D057130 | LCA5 | 167691 | MH |
| D057130 | IQCB1 | 9657 | MH |
| D057130 | GDF6 | 392255 | MH |
| D057130 | RDH12 | 145226 | MH |
| D057130 | RPE65 | 6121 | MH |
| D057130 | CEP290 | 80184 | MH |
| D057130 | RPGRIP1 | 57096 | MH |
| D057130 | NMNAT1 | 64802 | MH |
| D057130 | NPHP1 | 4867 | MH |
| D057130 | RD3 | 343035 | MH |
| D057130 | SPATA7 | 55812 | MH |
| D057130 | NXPH1 | 30010 | MH |
| D057130 | IMPDH1 | 3614 | SCR |
| D057130 | LRAT | 9227 | SCR |
| D057130 | AIPL1 | 23746 | SCR |
| D057130 | GUCY2D | 3000 | SCR |
| D057130 | IQCB1 | 9657 | SCR |
| D057130 | LCA5 | 167691 | SCR |
| D057130 | RPE65 | 6121 | SCR |
| D057130 | RPGRIP1 | 57096 | SCR |
| D057130 | NMNAT1 | 64802 | SCR |
| D057130 | CEP290 | 80184 | SCR |
| D057130 | RDH12 | 145226 | SCR |
| D057130 | RD3 | 343035 | SCR |
| D057130 | NPHP1 | 4867 | SCR |
| D057130 | NXPH1 | 30010 | SCR |
| D057130 | CRB1 | 23418 | Non-MeSH |
| D057130 | SDCCAG8 | 10806 | Non-MeSH |
| D057130 | RPGR | 6103 | Non-MeSH |
| D057130 | CRX | 1406 | Non-MeSH |
| D057130 | TULP1 | 7287 | Non-MeSH |
| D000309 | POMC | 5443 | MH |
| D000309 | FGD3 | 89846 | MH |
| D000309 | NR5A1 | 2516 | MH |
| D000309 | CDKN1C | 1028 | MH |
| D000309 | SF1 | 7536 | MH |
| D000309 | MC2R | 4158 | MH |
| D000309 | MRAP | 56246 | MH |
| D000309 | AAAS | 8086 | MH |
| D000309 | CYP11A1 | 1583 | MH |
| D000309 | CYP11B2 | 1585 | MH |
| D000309 | ABCD1 | 215 | MH |
| D000309 | AMN | 81693 | MH |
| D000309 | H19 | 283120 | MH |
| D000309 | MCM4 | 4173 | MH |
| D000309 | TBX19 | 9095 | MH |
| D000309 | CIB2 | 10518 | MH |
| D000309 | FGD2 | 221472 | MH |
| D000309 | AAAS | 8086 | SCR |
| D000309 | MC2R | 4158 | SCR |
| D000309 | H19 | 283120 | SCR |
| D000309 | MCM4 | 4173 | SCR |
| D000309 | TBX19 | 9095 | SCR |
| D000309 | CIB2 | 10518 | SCR |
| D000309 | FGD3 | 89846 | SCR |
| D000309 | CDKN1C | 1028 | SCR |
| D000309 | CYP11A1 | 1583 | Non-MeSH |
| D000309 | CYP11B2 | 1585 | Non-MeSH |
| D000309 | POMC | 5443 | Non-MeSH |
| D000309 | NR5A1 | 2516 | Non-MeSH |
| D000309 | SF1 | 7536 | Non-MeSH |
| D010013 | PPIB | 5479 | MH |
| D010013 | INTS1 | 26173 | MH |
| D010013 | CRTAP | 10491 | MH |
| D010013 | SERPINF1 | 5176 | MH |
| D010013 | ANO5 | 203859 | MH |
| D010013 | PLOD2 | 5352 | MH |
| D010013 | IFITM5 | 387733 | MH |
| D010013 | COL1A2 | 1278 | MH |
| D010013 | LHX2 | 9355 | MH |
| D010013 | WNT1 | 7471 | MH |
| D010013 | FKBP10 | 60681 | MH |
| D010013 | PPIB | 5479 | SCR |
| D010013 | CRTAP | 10491 | SCR |
| D010013 | SERPINF1 | 5176 | SCR |
| D010013 | ANO5 | 203859 | SCR |
| D010013 | PLOD2 | 5352 | SCR |
| D010013 | IFITM5 | 387733 | SCR |
| D010013 | COL1A2 | 1278 | SCR |
| D010013 | LHX2 | 9355 | SCR |
| D010013 | FKBP10 | 60681 | SCR |
| D010013 | FKBP10 | 60681 | Non-MeSH |
| D010013 | SP7 | 121340 | Non-MeSH |
| D006228 | MGP | 4256 | MH |
| D006228 | FAT4 | 79633 | MH |
| D006228 | TBC1D24 | 57465 | MH |
| D006228 | EZH2 | 2146 | MH |
| D006228 | MIPOL1 | 145282 | MH |
| D006228 | HYLS1 | 219844 | MH |
| D006228 | TRPS1 | 7227 | MH |
| D006228 | CLMP | 79827 | MH |
| D006228 | SALL4 | 57167 | MH |
| D006228 | ROBO3 | 64221 | MH |
| D006228 | CDH3 | 1001 | MH |
| D006228 | ERGIC2 | 51290 | MH |
| D006228 | FGD1 | 2245 | MH |
| D006228 | KIF22 | 3835 | MH |
| D006228 | CDH19 | 28513 | MH |
| D006228 | SMAD4 | 4089 | MH |
| D006228 | KLKB1 | 3818 | MH |
| D006228 | WNT7A | 7476 | MH |
| D006228 | RIMS4 | 140730 | MH |
| D006228 | TP63 | 8626 | MH |
| D006228 | COL11A1 | 1301 | MH |
| D006228 | GYPE | 2996 | MH |
| D006228 | FGFR1 | 2260 | MH |
| D006228 | DCHS1 | 8642 | MH |
| D006228 | LMNA | 4000 | MH |
| D006228 | HOXA13 | 3209 | MH |
| D006228 | FGF16 | 8823 | MH |
| D006228 | NFIX | 4784 | MH |
| D006228 | PRG4 | 10216 | MH |
| D006228 | PITX1 | 5307 | MH |
| D006228 | SETBP1 | 26040 | MH |
| D006228 | NOG | 9241 | MH |
| D006228 | GJB2 | 2706 | MH |
| D006228 | NUBP2 | 10101 | MH |
| D006228 | EZH1 | 2145 | MH |
| D006228 | FGFR2 | 2263 | MH |
| D006228 | FGFR3 | 2261 | MH |
| D006228 | RBBP8 | 5932 | MH |
| D006228 | FLNA | 2316 | MH |
| D006228 | CRISP3 | 10321 | MH |
| D006228 | LMBR1 | 64327 | MH |
| D006228 | FAT4 | 79633 | SCR |
| D006228 | MGP | 4256 | SCR |
| D006228 | TBC1D24 | 57465 | SCR |
| D006228 | EZH2 | 2146 | SCR |
| D006228 | MIPOL1 | 145282 | SCR |
| D006228 | HYLS1 | 219844 | SCR |
| D006228 | TRPS1 | 7227 | SCR |
| D006228 | CLMP | 79827 | SCR |
| D006228 | SALL4 | 57167 | SCR |
| D006228 | ROBO3 | 64221 | SCR |
| D006228 | ERGIC2 | 51290 | SCR |
| D006228 | FGD1 | 2245 | SCR |
| D006228 | KIF22 | 3835 | SCR |
| D006228 | CDH19 | 28513 | SCR |
| D006228 | SMAD4 | 4089 | SCR |
| D006228 | KLKB1 | 3818 | SCR |
| D006228 | WNT7A | 7476 | SCR |
| D006228 | COL11A1 | 1301 | SCR |
| D006228 | GYPE | 2996 | SCR |
| D006228 | FGFR1 | 2260 | SCR |
| D006228 | DCHS1 | 8642 | SCR |
| D006228 | FGF16 | 8823 | SCR |
| D006228 | HOXA13 | 3209 | SCR |
| D006228 | NFIX | 4784 | SCR |
| D006228 | CLCF1 | 23529 | SCR |
| D006228 | LMNA | 4000 | SCR |
| D006228 | PRG4 | 10216 | SCR |
| D006228 | PITX1 | 5307 | SCR |
| D006228 | SETBP1 | 26040 | SCR |
| D006228 | NOG | 9241 | SCR |
| D006228 | CLC | 1178 | SCR |
| D006228 | GJB2 | 2706 | SCR |
| D006228 | NUBP2 | 10101 | SCR |
| D006228 | EZH1 | 2145 | SCR |
| D006228 | FGFR2 | 2263 | SCR |
| D006228 | FGFR3 | 2261 | SCR |
| D006228 | FLNA | 2316 | SCR |
| D006228 | CRISP3 | 10321 | SCR |
| D006228 | LMBR1 | 64327 | SCR |
| D006228 | KIF7 | 374654 | Non-MeSH |
| D008850 | SNX3 | 8724 | MH |
| D008850 | VSX2 | 338917 | MH |
| D008850 | MFRP | 83552 | MH |
| D008850 | VSNL1 | 7447 | MH |
| D008850 | OTX2 | 5015 | MH |
| D008850 | NAA10 | 8260 | MH |
| D008850 | BCOR | 54880 | MH |
| D008850 | HCCS | 3052 | MH |
| D008850 | HDAC6 | 10013 | MH |
| D008850 | RARB | 5915 | MH |
| D008850 | SOX2 | 6657 | MH |
| D008850 | SIX6 | 4990 | MH |
| D008850 | GDF3 | 9573 | MH |
| D008850 | RTN3 | 10313 | MH |
| D008850 | GDF6 | 392255 | MH |
| D008850 | TENM3 | 55714 | MH |
| D008850 | RAX | 30062 | MH |
| D008850 | STRA6 | 64220 | MH |
| D008850 | SHH | 6469 | MH |
| D008850 | BMP4 | 652 | MH |
| D008850 | OTX2 | 5015 | SCR |
| D008850 | NAA10 | 8260 | SCR |
| D008850 | BCOR | 54880 | SCR |
| D008850 | HCCS | 3052 | SCR |
| D008850 | SNX3 | 8724 | SCR |
| D008850 | GDF6 | 392255 | SCR |
| D008850 | TENM3 | 55714 | SCR |
| D008850 | STRA6 | 64220 | SCR |
| D008850 | VSX2 | 338917 | SCR |
| D008850 | BMP4 | 652 | SCR |
| D008850 | MFRP | 83552 | SCR |
| D008850 | GDF3 | 9573 | Non-MeSH |
| D008850 | GDF6 | 392255 | Non-MeSH |
| D008831 | IER3IP1 | 51124 | MH |
| D008831 | CEP152 | 22995 | MH |
| D008831 | RAD50 | 10111 | MH |
| D008831 | NHEJ1 | 79840 | MH |
| D008831 | CASK | 8573 | MH |
| D008831 | SNX3 | 8724 | MH |
| D008831 | FAM20C | 56975 | MH |
| D008831 | VPS13B | 157680 | MH |
| D008831 | MED17 | 9440 | MH |
| D008831 | SLC25A19 | 60386 | MH |
| D008831 | ZEB2 | 9839 | MH |
| D008831 | EFTUD2 | 9343 | MH |
| D008831 | RIMS4 | 140730 | MH |
| D008831 | RNU4ATAC | 100151683 | MH |
| D008831 | PCNT | 5116 | MH |
| D008831 | RAB3GAP1 | 22930 | MH |
| D008831 | WDR62 | 284403 | MH |
| D008831 | MCPH1 | 79648 | MH |
| D008831 | ARFGEF2 | 10564 | MH |
| D008831 | ATR | 545 | MH |
| D008831 | SFRP1 | 6422 | MH |
| D008831 | KMT2A | 4297 | MH |
| D008831 | MYCN | 4613 | MH |
| D008831 | RAB23 | 51715 | MH |
| D008831 | RAB18 | 22931 | MH |
| D008831 | STIL | 6491 | MH |
| D008831 | CENPJ | 55835 | MH |
| D008831 | PHGDH | 26227 | MH |
| D008831 | RBBP8 | 5932 | MH |
| D008831 | PSAT1 | 29968 | MH |
| D008831 | ASPM | 259266 | MH |
| D008831 | CEP152 | 22995 | SCR |
| D008831 | RAD50 | 10111 | SCR |
| D008831 | NHEJ1 | 79840 | SCR |
| D008831 | FAM20C | 56975 | SCR |
| D008831 | SNX3 | 8724 | SCR |
| D008831 | CASK | 8573 | SCR |
| D008831 | VPS13B | 157680 | SCR |
| D008831 | SLC25A19 | 60386 | SCR |
| D008831 | ZEB2 | 9839 | SCR |
| D008831 | EFTUD2 | 9343 | SCR |
| D008831 | RIMS4 | 140730 | SCR |
| D008831 | RNU4ATAC | 100151683 | SCR |
| D008831 | PCNT | 5116 | SCR |
| D008831 | RAB3GAP1 | 22930 | SCR |
| D008831 | MCPH1 | 79648 | SCR |
| D008831 | ATR | 545 | SCR |
| D008831 | SFRP1 | 6422 | SCR |
| D008831 | KMT2A | 4297 | SCR |
| D008831 | MYCN | 4613 | SCR |
| D008831 | RAB18 | 22931 | SCR |
| D008831 | STIL | 6491 | SCR |
| D008831 | CENPJ | 55835 | SCR |
| D008831 | PHGDH | 26227 | SCR |
| D008831 | RBBP8 | 5932 | SCR |
| D008831 | PSAT1 | 29968 | SCR |
| D008831 | ASPM | 259266 | SCR |
| D008831 | CEP152 | 22995 | Non-MeSH |
| D015418 | OPA1 | 4976 | MH |
| D015418 | IQCB1 | 9657 | MH |
| D015418 | CEP290 | 80184 | MH |
| D015418 | ATR | 545 | MH |
| D015418 | ANTXR1 | 84168 | MH |
| D015418 | NPHP1 | 4867 | MH |
| D015418 | TMEM126A | 84233 | MH |
| D015418 | NXPH1 | 30010 | MH |
| D015418 | IQCB1 | 9657 | SCR |
| D015418 | CEP290 | 80184 | SCR |
| D015418 | ATR | 545 | SCR |
| D015418 | ANTXR1 | 84168 | SCR |
| D015418 | NPHP1 | 4867 | SCR |
| D015418 | TMEM126A | 84233 | SCR |
| D015418 | NXPH1 | 30010 | SCR |
| D015418 | SDCCAG8 | 10806 | Non-MeSH |
| D020788 | BBS4 | 585 | MH |
| D020788 | TRIM32 | 22954 | MH |
| D020788 | BBS12 | 166379 | MH |
| D020788 | ARL6 | 84100 | MH |
| D020788 | TMEM67 | 91147 | MH |
| D020788 | BBS1 | 582 | MH |
| D020788 | CCDC28B | 79140 | MH |
| D020788 | BBS9 | 27241 | MH |
| D020788 | BBS2 | 583 | MH |
| D020788 | TTC8 | 123016 | MH |
| D020788 | MOCOS | 55034 | MH |
| D020788 | LZTFL1 | 54585 | MH |
| D020788 | BBS7 | 55212 | MH |
| D020788 | CEP290 | 80184 | MH |
| D020788 | MKS1 | 54903 | MH |
| D020788 | MKKS | 8195 | MH |
| D020788 | BBS10 | 79738 | MH |
| D020788 | BBS4 | 585 | SCR |
| D020788 | TRIM32 | 22954 | SCR |
| D020788 | ARL6 | 84100 | SCR |
| D020788 | MKKS | 8195 | SCR |
| D020788 | TMEM67 | 91147 | SCR |
| D020788 | BBS12 | 166379 | SCR |
| D020788 | BBS1 | 582 | SCR |
| D020788 | CCDC28B | 79140 | SCR |
| D020788 | MOCOS | 55034 | SCR |
| D020788 | TTC8 | 123016 | SCR |
| D020788 | BBS9 | 27241 | SCR |
| D020788 | BBS2 | 583 | SCR |
| D020788 | BBS7 | 55212 | SCR |
| D020788 | CEP290 | 80184 | SCR |
| D020788 | MKS1 | 54903 | SCR |
| D020788 | BBS10 | 79738 | SCR |
| D020788 | WDPCP | 51057 | Non-MeSH |
| D002972 | SP7 | 121340 | MH |
| D002972 | FAM20C | 56975 | MH |
| D002972 | COL2A1 | 1280 | MH |
| D002972 | STAC3 | 246329 | MH |
| D002972 | IRF6 | 3664 | MH |
| D002972 | TTF2 | 8458 | MH |
| D002972 | RIPK4 | 54101 | MH |
| D002972 | TP63 | 8626 | MH |
| D002972 | UBB | 7314 | MH |
| D002972 | FGFR1 | 2260 | MH |
| D002972 | MSX1 | 4487 | MH |
| D002972 | FOXE1 | 2304 | MH |
| D002972 | MID1 | 4281 | MH |
| D002972 | SUMO1 | 7341 | MH |
| D002972 | BMP4 | 652 | MH |
| D002972 | FAM20C | 56975 | SCR |
| D002972 | SP7 | 121340 | SCR |
| D002972 | COL2A1 | 1280 | SCR |
| D002972 | STAC3 | 246329 | SCR |
| D002972 | IRF6 | 3664 | SCR |
| D002972 | TTF2 | 8458 | SCR |
| D002972 | RIPK4 | 54101 | SCR |
| D002972 | TP63 | 8626 | SCR |
| D002972 | FGFR1 | 2260 | SCR |
| D002972 | MSX1 | 4487 | SCR |
| D002972 | FOXE1 | 2304 | SCR |
| D002972 | SUMO1 | 7341 | SCR |
| D002972 | MID1 | 4281 | SCR |
| D002972 | BMP4 | 652 | SCR |
| D002972 | TP63 | 8626 | Non-MeSH |
| D028361 | NDUFS3 | 4722 | MH |
| D028361 | DARS2 | 55157 | MH |
| D028361 | UQCRB | 7381 | MH |
| D028361 | NDUFB9 | 4715 | MH |
| D028361 | NDUFS1 | 4719 | MH |
| D028361 | COX10 | 1352 | MH |
| D028361 | ACADVL | 37 | MH |
| D028361 | CPS1 | 1373 | MH |
| D028361 | NDUFB3 | 4709 | MH |
| D028361 | NDUFV1 | 4723 | MH |
| D028361 | NDUFAF1 | 51103 | MH |
| D028361 | MRPS22 | 56945 | MH |
| D028361 | NDUFS4 | 4724 | MH |
| D028361 | ELAC2 | 60528 | MH |
| D028361 | NDUFA11 | 126328 | MH |
| D028361 | FASTKD2 | 22868 | MH |
| D028361 | PC | 5091 | MH |
| D028361 | TUFM | 7284 | MH |
| D028361 | COX15 | 1355 | MH |
| D028361 | TACO1 | 51204 | MH |
| D028361 | SDHA | 6389 | MH |
| D028361 | PDSS2 | 57107 | MH |
| D028361 | NDUFA10 | 4705 | MH |
| D028361 | COX14 | 84987 | MH |
| D028361 | COX20 | 116228 | MH |
| D028361 | NDUFV2 | 4729 | MH |
| D028361 | COA5 | 493753 | MH |
| D028361 | NDUFS7 | 374291 | MH |
| D028361 | PDHA1 | 5160 | MH |
| D028361 | NUBPL | 80224 | MH |
| D028361 | CYC1 | 1537 | MH |
| D028361 | NDUFS2 | 4720 | MH |
| D028361 | FXN | 2395 | MH |
| D028361 | BCS1L | 617 | MH |
| D028361 | SURF1 | 6834 | MH |
| D028361 | SLC25A12 | 8604 | MH |
| D028361 | SLC25A3 | 5250 | MH |
| D028361 | NDUFAF5 | 79133 | MH |
| D028361 | CISD2 | 493856 | MH |
| D028361 | COQ2 | 27235 | MH |
| D028361 | COQ6 | 51004 | MH |
| D028361 | OPA1 | 4976 | MH |
| D028361 | SFXN4 | 119559 | MH |
| D028361 | ACAD9 | 28976 | MH |
| D028361 | NDUFAF4 | 29078 | MH |
| D028361 | FOXRED1 | 55572 | MH |
| D028361 | NDUFAF3 | 25915 | MH |
| D028361 | SDHB | 6390 | MH |
| D028361 | COQ9 | 57017 | MH |
| D028361 | TSFM | 10102 | MH |
| D028361 | PDSS1 | 23590 | MH |
| D028361 | NDUFA1 | 4694 | MH |
| D028361 | SDHAF1 | 644096 | MH |
| D028361 | UQCRQ | 27089 | MH |
| D028361 | NDUFAF2 | 91942 | MH |
| D028361 | CPT2 | 1376 | MH |
| D028361 | LMBR1 | 64327 | MH |
| D028361 | NDUFS3 | 4722 | SCR |
| D028361 | DARS2 | 55157 | SCR |
| D028361 | UQCRB | 7381 | SCR |
| D028361 | NDUFB9 | 4715 | SCR |
| D028361 | NDUFS1 | 4719 | SCR |
| D028361 | NDUFB3 | 4709 | SCR |
| D028361 | NDUFV1 | 4723 | SCR |
| D028361 | NDUFAF1 | 51103 | SCR |
| D028361 | MRPS22 | 56945 | SCR |
| D028361 | NDUFS4 | 4724 | SCR |
| D028361 | NDUFA11 | 126328 | SCR |
| D028361 | TUFM | 7284 | SCR |
| D028361 | PDSS1 | 23590 | SCR |
| D028361 | PDSS2 | 57107 | SCR |
| D028361 | NDUFV2 | 4729 | SCR |
| D028361 | POLG | 5428 | SCR |
| D028361 | CYC1 | 1537 | SCR |
| D028361 | NDUFS2 | 4720 | SCR |
| D028361 | BCS1L | 617 | SCR |
| D028361 | SLC25A12 | 8604 | SCR |
| D028361 | SLC25A3 | 5250 | SCR |
| D028361 | CISD2 | 493856 | SCR |
| D028361 | COQ2 | 27235 | SCR |
| D028361 | COQ6 | 51004 | SCR |
| D028361 | NDUFAF3 | 25915 | SCR |
| D028361 | NDUFAF4 | 29078 | SCR |
| D028361 | FOXRED1 | 55572 | SCR |
| D028361 | UQCRQ | 27089 | SCR |
| D028361 | NDUFA1 | 4694 | SCR |
| D028361 | COQ9 | 57017 | SCR |
| D028361 | NDUFAF5 | 79133 | SCR |
| D028361 | NUBPL | 80224 | SCR |
| D028361 | SDHAF1 | 644096 | SCR |
| D028361 | ACAD9 | 28976 | SCR |
| D028361 | NDUFAF2 | 91942 | SCR |
| D028361 | CPT2 | 1376 | SCR |
| D028361 | LMBR1 | 64327 | SCR |
| D028361 | BCS1L | 617 | Non-MeSH |
| D012878 | RSPO1 | 284654 | MH |
| D012878 | PTEN | 5728 | MH |
| D012878 | CYLD | 1540 | MH |
| D012878 | NF2 | 4771 | MH |
| D012878 | BRAF | 673 | MH |
| D012878 | KRT14 | 3861 | MH |
| D012878 | SMO | 6608 | MH |
| D012878 | STK11 | 6794 | MH |
| D012878 | ARHGAP11A | 9824 | MH |
| D012878 | HRAS | 3265 | MH |
| D012878 | RASA1 | 5921 | MH |
| D012878 | PTCH2 | 8643 | MH |
| D012878 | TP53 | 7157 | MH |
| D012878 | CDKN2A | 1029 | MH |
| D012878 | PTCH1 | 5727 | MH |
| D012878 | CDK4 | 1019 | SCR |
| D012878 | CYLD | 1540 | SCR |
| D012878 | NF2 | 4771 | SCR |
| D012878 | TERT | 7015 | SCR |
| D012878 | KRT14 | 3861 | SCR |
| D012878 | XRCC3 | 7517 | SCR |
| D012878 | CDKN2A | 1029 | SCR |
| D012878 | CYLD | 1540 | Non-MeSH |
| D005532 | FAT4 | 79633 | MH |
| D005532 | SNX3 | 8724 | MH |
| D005532 | MIPOL1 | 145282 | MH |
| D005532 | HOXD13 | 3239 | MH |
| D005532 | HOXD10 | 3236 | MH |
| D005532 | TRPS1 | 7227 | MH |
| D005532 | SMARCA2 | 6595 | MH |
| D005532 | GJA1 | 2697 | MH |
| D005532 | CDH19 | 28513 | MH |
| D005532 | WNT7A | 7476 | MH |
| D005532 | FGFR1 | 2260 | MH |
| D005532 | DCHS1 | 8642 | MH |
| D005532 | HOXA13 | 3209 | MH |
| D005532 | NOG | 9241 | MH |
| D005532 | NUBP2 | 10101 | MH |
| D005532 | PANK2 | 80025 | MH |
| D005532 | FGFR2 | 2263 | MH |
| D005532 | PAX6 | 5080 | MH |
| D005532 | CRISP3 | 10321 | MH |
| D005532 | LMBR1 | 64327 | MH |
| D005532 | FAT4 | 79633 | SCR |
| D005532 | SNX3 | 8724 | SCR |
| D005532 | MIPOL1 | 145282 | SCR |
| D005532 | HOXD13 | 3239 | SCR |
| D005532 | TRPS1 | 7227 | SCR |
| D005532 | SMARCA2 | 6595 | SCR |
| D005532 | GJA1 | 2697 | SCR |
| D005532 | CDH19 | 28513 | SCR |
| D005532 | WNT7A | 7476 | SCR |
| D005532 | FGFR1 | 2260 | SCR |
| D005532 | DCHS1 | 8642 | SCR |
| D005532 | HOXA13 | 3209 | SCR |
| D005532 | NOG | 9241 | SCR |
| D005532 | NUBP2 | 10101 | SCR |
| D005532 | PANK2 | 80025 | SCR |
| D005532 | FGFR2 | 2263 | SCR |
| D005532 | PAX6 | 5080 | SCR |
| D005532 | CRISP3 | 10321 | SCR |
| D005532 | LMBR1 | 64327 | SCR |
| D005532 | HOXD10 | 3236 | Non-MeSH |
| D020821 | SLC2A1 | 6513 | MH |
| D020821 | ATP1A3 | 478 | MH |
| D020821 | SGCE | 8910 | MH |
| D020821 | TOR1A | 1861 | MH |
| D020821 | PEA15 | 8682 | MH |
| D020821 | RAX | 30062 | MH |
| D020821 | PRKRA | 8575 | MH |
| D020821 | DRD5 | 1816 | MH |
| D020821 | TAF10 | 6881 | MH |
| D020821 | TAF1 | 6872 | MH |
| D020821 | BCAP31 | 10134 | MH |
| D020821 | NELFE | 7936 | MH |
| D020821 | SLC6A3 | 6531 | MH |
| D020821 | ACTB | 60 | MH |
| D020821 | DRD2 | 1813 | MH |
| D020821 | DRD2 | 1813 | SCR |
| D020821 | SLC2A1 | 6513 | SCR |
| D020821 | ATP1A3 | 478 | SCR |
| D020821 | SGCE | 8910 | SCR |
| D020821 | PEA15 | 8682 | SCR |
| D020821 | RAX | 30062 | SCR |
| D020821 | PRKRA | 8575 | SCR |
| D020821 | TAF1 | 6872 | SCR |
| D020821 | NELFE | 7936 | SCR |
| D020821 | SLC6A3 | 6531 | SCR |
| D020821 | ACTB | 60 | SCR |
| D020821 | TAF10 | 6881 | SCR |
| D008133 | SCN4B | 6330 | MH |
| D008133 | ALG10B | 144245 | MH |
| D008133 | KCNE2 | 9992 | MH |
| D008133 | ALG10 | 84920 | MH |
| D008133 | TYMS | 7298 | MH |
| D008133 | SCN5A | 6331 | MH |
| D008133 | CACNA1C | 775 | MH |
| D008133 | AKAP9 | 10142 | MH |
| D008133 | KCNE1 | 3753 | MH |
| D008133 | SNTA1 | 6640 | MH |
| D008133 | KCNH2 | 3757 | MH |
| D008133 | FRS2 | 10818 | MH |
| D008133 | CAV3 | 859 | MH |
| D008133 | KCNQ1 | 3784 | MH |
| D008133 | ANK2 | 287 | MH |
| D008133 | KCNJ2 | 3759 | MH |
| D008133 | SEC61G | 23480 | MH |
| D008133 | SCN4B | 6330 | SCR |
| D008133 | KCNE2 | 9992 | SCR |
| D008133 | ALG10 | 84920 | SCR |
| D008133 | TYMS | 7298 | SCR |
| D008133 | AKAP9 | 10142 | SCR |
| D008133 | KCNE1 | 3753 | SCR |
| D008133 | SNTA1 | 6640 | SCR |
| D008133 | CACNA1C | 775 | SCR |
| D008133 | ALG10B | 144245 | SCR |
| D008133 | FRS2 | 10818 | SCR |
| D008133 | CAV3 | 859 | SCR |
| D008133 | KCNQ1 | 3784 | SCR |
| D008133 | ANK2 | 287 | SCR |
| D008133 | KCNH2 | 3757 | SCR |
| D008133 | KCNJ5 | 3762 | Non-MeSH |
| D010300 | HTRA2 | 27429 | MH |
| D010300 | PINK1 | 65018 | MH |
| D010300 | ADH1C | 126 | MH |
| D010300 | SNCA | 6622 | MH |
| D010300 | GIGYF2 | 26058 | MH |
| D010300 | SYNJ1 | 8867 | MH |
| D010300 | PLA2G6 | 8398 | MH |
| D010300 | TBP | 6908 | MH |
| D010300 | SLC6A3 | 6531 | MH |
| D010300 | GBA | 2629 | MH |
| D010300 | MAPT | 4137 | MH |
| D010300 | PARK7 | 11315 | MH |
| D010300 | HTRA2 | 27429 | SCR |
| D010300 | PINK1 | 65018 | SCR |
| D010300 | ADH1C | 126 | SCR |
| D010300 | GIGYF2 | 26058 | SCR |
| D010300 | SLC6A3 | 6531 | SCR |
| D010300 | TBP | 6908 | SCR |
| D010300 | GBA | 2629 | SCR |
| D010300 | MAPT | 4137 | SCR |
| D010300 | PARK7 | 11315 | SCR |
| D010300 | EIF4G3 | 8672 | Non-MeSH |
| D010300 | EIF4G1 | 1981 | Non-MeSH |
| D010300 | VPS35 | 55737 | Non-MeSH |
| D008268 | CFH | 3075 | MH |
| D008268 | HMCN1 | 83872 | MH |
| D008268 | ABCA4 | 24 | MH |
| D008268 | HTRA1 | 5654 | MH |
| D008268 | ROM1 | 6094 | MH |
| D008268 | CFHR1 | 3078 | MH |
| D008268 | C3 | 718 | MH |
| D008268 | CDH3 | 1001 | MH |
| D008268 | C2 | 717 | MH |
| D008268 | ELOVL4 | 6785 | MH |
| D008268 | BEST1 | 7439 | MH |
| D008268 | CNGB3 | 54714 | MH |
| D008268 | RAX2 | 84839 | MH |
| D008268 | C9 | 735 | MH |
| D008268 | CST3 | 1471 | MH |
| D008268 | DHDDS | 79947 | MH |
| D008268 | PRPH2 | 5961 | MH |
| D008268 | PRPH | 5630 | MH |
| D008268 | FBLN5 | 10516 | MH |
| D008268 | CFHR3 | 10878 | MH |
| D008268 | ERCC6 | 2074 | MH |
| D008268 | CFI | 3426 | MH |
| D008268 | TLR4 | 7099 | MH |
| D008268 | PROM1 | 8842 | MH |
| D008268 | CST3 | 1471 | SCR |
| D008268 | C3 | 718 | SCR |
| D008268 | CDH3 | 1001 | SCR |
| D008268 | ABCA4 | 24 | SCR |
| D008268 | ELOVL4 | 6785 | SCR |
| D008268 | TLR4 | 7099 | SCR |
| D008268 | CNGB3 | 54714 | SCR |
| D008268 | PROM1 | 8842 | SCR |
| D008268 | CFHR1 | 3078 | Non-MeSH |
| D008268 | C9 | 735 | Non-MeSH |
| D008268 | RP1L1 | 94137 | Non-MeSH |
| D008268 | CDH3 | 1001 | Non-MeSH |
| D008268 | ARMS2 | 387715 | Non-MeSH |
| D008268 | CFHR3 | 10878 | Non-MeSH |
| D008268 | C2 | 717 | Non-MeSH |
| D008268 | ERCC6 | 2074 | Non-MeSH |
| D008268 | DHDDS | 79947 | Non-MeSH |
| D008268 | CFI | 3426 | Non-MeSH |
| D008268 | PRPH2 | 5961 | Non-MeSH |
| D008268 | ELOVL4 | 6785 | Non-MeSH |
| D008268 | HTRA1 | 5654 | Non-MeSH |
| D008268 | ROM1 | 6094 | Non-MeSH |
| D008268 | PRPH | 5630 | Non-MeSH |
| D008268 | PROM1 | 8842 | Non-MeSH |
| D003920 | KCNJ11 | 3767 | MH |
| D003920 | IER3IP1 | 51124 | MH |
| D003920 | SLC30A8 | 169026 | MH |
| D003920 | PON1 | 5444 | MH |
| D003920 | INS | 3630 | MH |
| D003920 | ABCC8 | 6833 | MH |
| D003920 | SUMO4 | 387082 | MH |
| D003920 | OAS1 | 4938 | MH |
| D003920 | ITPA | 3704 | MH |
| D003920 | RFX6 | 222546 | MH |
| D003920 | PCAT1 | 100750225 | MH |
| D003920 | MAPK8IP1 | 9479 | MH |
| D003920 | HNF1A | 6927 | MH |
| D003920 | DCAF17 | 80067 | MH |
| D003920 | IRS1 | 3667 | MH |
| D003920 | GCGR | 2642 | MH |
| D003920 | SOD2 | 6648 | MH |
| D003920 | TCF4 | 6925 | MH |
| D003920 | CTLA4 | 1493 | MH |
| D003920 | IRS2 | 8660 | MH |
| D003920 | GPD2 | 2820 | MH |
| D003920 | HNF1B | 6928 | MH |
| D003920 | FOXP3 | 50943 | MH |
| D003920 | HNF4A | 3172 | MH |
| D003920 | TCF7L2 | 6934 | MH |
| D003920 | SLC2A2 | 6514 | MH |
| D003920 | NEUROD1 | 4760 | MH |
| D003920 | ENPP1 | 5167 | MH |
| D003920 | SLC19A2 | 10560 | MH |
| D003920 | CDKAL1 | 54901 | MH |
| D003920 | PTF1A | 256297 | MH |
| D003920 | PDX1 | 3651 | MH |
| D003920 | ITPR3 | 3710 | MH |
| D003920 | AKT2 | 208 | MH |
| D003920 | MTNR1B | 4544 | MH |
| D003920 | GLIS3 | 169792 | MH |
| D003920 | IL6 | 3569 | MH |
| D003920 | UMOD | 7369 | MH |
| D003920 | HMGA1 | 3159 | MH |
| D003920 | EPO | 2056 | MH |
| D003920 | HGF | 3082 | MH |
| D003920 | INSR | 3643 | MH |
| D003920 | UCP3 | 7352 | MH |
| D003920 | IGF2BP2 | 10644 | MH |
| D003920 | IL2RA | 3559 | MH |
| D003920 | PTPN22 | 26191 | MH |
| D003920 | PAX4 | 5078 | MH |
| D003920 | CAPN10 | 11132 | MH |
| D003920 | SLC2A4 | 6517 | MH |
| D003920 | SH2B3 | 10019 | MH |
| D003920 | RETN | 56729 | MH |
| D003920 | FOXC2 | 2303 | MH |
| D003920 | IL1RN | 3557 | MH |
| D003920 | VEGFA | 7422 | MH |
| D003920 | SPPL3 | 121665 | MH |
| D003920 | HSF5 | 124535 | MH |
| D003920 | GCK | 2645 | MH |
| D003920 | INS | 3630 | SCR |
| D003920 | ABCC8 | 6833 | SCR |
| D003920 | GLIS3 | 169792 | SCR |
| D003920 | ITPA | 3704 | SCR |
| D003920 | RFX6 | 222546 | SCR |
| D003920 | INSR | 3643 | SCR |
| D003920 | SLC19A2 | 10560 | SCR |
| D003920 | DCAF17 | 80067 | SCR |
| D003920 | GCK | 2645 | SCR |
| D003920 | KCNJ11 | 3767 | Non-MeSH |
| D003920 | IER3IP1 | 51124 | Non-MeSH |
| D003920 | SLC30A8 | 169026 | Non-MeSH |
| D003920 | PON1 | 5444 | Non-MeSH |
| D003920 | ABCC8 | 6833 | Non-MeSH |
| D003920 | PCAT1 | 100750225 | Non-MeSH |
| D003920 | MAPK8IP1 | 9479 | Non-MeSH |
| D003920 | PPOX | 5498 | Non-MeSH |
| D003920 | HNF1A | 6927 | Non-MeSH |
| D003920 | IRS1 | 3667 | Non-MeSH |
| D003920 | GCGR | 2642 | Non-MeSH |
| D003920 | SOD2 | 6648 | Non-MeSH |
| D003920 | TCF4 | 6925 | Non-MeSH |
| D003920 | CTLA4 | 1493 | Non-MeSH |
| D003920 | IRS2 | 8660 | Non-MeSH |
| D003920 | GPD2 | 2820 | Non-MeSH |
| D003920 | HNF1B | 6928 | Non-MeSH |
| D003920 | HNF4A | 3172 | Non-MeSH |
| D003920 | TCF7L2 | 6934 | Non-MeSH |
| D003920 | SLC2A2 | 6514 | Non-MeSH |
| D003920 | NEUROD1 | 4760 | Non-MeSH |
| D003920 | ENPP1 | 5167 | Non-MeSH |
| D003920 | AVP | 551 | Non-MeSH |
| D003920 | CDKAL1 | 54901 | Non-MeSH |
| D003920 | PTF1A | 256297 | Non-MeSH |
| D003920 | PDX1 | 3651 | Non-MeSH |
| D003920 | ITPR3 | 3710 | Non-MeSH |
| D003920 | AKT2 | 208 | Non-MeSH |
| D003920 | MTNR1B | 4544 | Non-MeSH |
| D003920 | IL6 | 3569 | Non-MeSH |
| D003920 | UMOD | 7369 | Non-MeSH |
| D003920 | HMGA1 | 3159 | Non-MeSH |
| D003920 | EPO | 2056 | Non-MeSH |
| D003920 | HGF | 3082 | Non-MeSH |
| D003920 | UCP3 | 7352 | Non-MeSH |
| D003920 | IGF2BP2 | 10644 | Non-MeSH |
| D003920 | IL2RA | 3559 | Non-MeSH |
| D003920 | PAX4 | 5078 | Non-MeSH |
| D003920 | CAPN10 | 11132 | Non-MeSH |
| D003920 | SLC2A4 | 6517 | Non-MeSH |
| D003920 | SH2B3 | 10019 | Non-MeSH |
| D003920 | RETN | 56729 | Non-MeSH |
| D003920 | FOXC2 | 2303 | Non-MeSH |
| D003920 | IL1RN | 3557 | Non-MeSH |
| D003920 | VEGFA | 7422 | Non-MeSH |
| D003920 | SPPL3 | 121665 | Non-MeSH |
| D003920 | HSF5 | 124535 | Non-MeSH |
| D003920 | GCK | 2645 | Non-MeSH |
| D002872 | DHODH | 1723 | MH |
| D002872 | KANSL1 | 284058 | MH |
| D002872 | EHMT1 | 79813 | MH |
| D002872 | SHANK3 | 85358 | SCR |
| D002872 | DHODH | 1723 | SCR |
| D002872 | KANSL1 | 284058 | SCR |
| D002872 | EHMT1 | 79813 | SCR |
| D002872 | HDAC4 | 9759 | Non-MeSH |
| D002872 | SHANK3 | 85358 | Non-MeSH |
| D013964 | TRIM33 | 51592 | MH |
| D013964 | SLC25A19 | 60386 | MH |
| D013964 | NCOA4 | 8031 | MH |
| D013964 | TRIM24 | 8805 | MH |
| D013964 | CCDC6 | 8030 | MH |
| D013964 | MINPP1 | 9562 | MH |
| D013964 | NTRK1 | 4914 | MH |
| D013964 | GOLGA5 | 9950 | MH |
| D013964 | RET | 5979 | MH |
| D013964 | HRH4 | 59340 | MH |
| D013964 | PCM1 | 5108 | MH |
| D013964 | NDUFA13 | 51079 | MH |
| D013964 | TRIM33 | 51592 | SCR |
| D013964 | SLC25A19 | 60386 | SCR |
| D013964 | NCOA4 | 8031 | SCR |
| D013964 | TRIM24 | 8805 | SCR |
| D013964 | CCDC6 | 8030 | SCR |
| D013964 | NTRK1 | 4914 | SCR |
| D013964 | GOLGA5 | 9950 | SCR |
| D013964 | HRH4 | 59340 | SCR |
| D013964 | RET | 5979 | SCR |
| D013964 | PCM1 | 5108 | SCR |
| D013964 | NDUFA13 | 51079 | SCR |
| D013964 | TRIM33 | 51592 | Non-MeSH |
| D013964 | SLC25A19 | 60386 | Non-MeSH |
| D013964 | NCOA4 | 8031 | Non-MeSH |
| D013964 | TRIM24 | 8805 | Non-MeSH |
| D013964 | MINPP1 | 9562 | Non-MeSH |
| D013964 | CCDC6 | 8030 | Non-MeSH |
| D013964 | GOLGA5 | 9950 | Non-MeSH |
| D013964 | HRH4 | 59340 | Non-MeSH |
| D013964 | PCM1 | 5108 | Non-MeSH |
| D016511 | GINS1 | 9837 | MH |
| D016511 | TAP2 | 6891 | MH |
| D016511 | CD3E | 916 | MH |
| D016511 | LIG4 | 3981 | MH |
| D016511 | TAPBP | 6892 | MH |
| D016511 | NHEJ1 | 79840 | MH |
| D016511 | IL2RG | 3561 | MH |
| D016511 | RAG2 | 5897 | MH |
| D016511 | RAG1 | 5896 | MH |
| D016511 | GINS2 | 51659 | MH |
| D016511 | ADA | 100 | MH |
| D016511 | DCLRE1C | 64421 | MH |
| D016511 | TAP1 | 6890 | MH |
| D016511 | AK2 | 204 | MH |
| D016511 | LCK | 3932 | MH |
| D016511 | CIITA | 4261 | MH |
| D016511 | GINS1 | 9837 | SCR |
| D016511 | RFXAP | 5994 | SCR |
| D016511 | TAP2 | 6891 | SCR |
| D016511 | TAPBP | 6892 | SCR |
| D016511 | NHEJ1 | 79840 | SCR |
| D016511 | RAG2 | 5897 | SCR |
| D016511 | RAG1 | 5896 | SCR |
| D016511 | GINS2 | 51659 | SCR |
| D016511 | ADA | 100 | SCR |
| D016511 | TAP1 | 6890 | SCR |
| D016511 | RFX5 | 5993 | SCR |
| D016511 | AK2 | 204 | SCR |
| D016511 | CIITA | 4261 | SCR |
| D016511 | LIG4 | 3981 | SCR |
| D016511 | ADA | 100 | Non-MeSH |
| D016511 | DCLRE1C | 64421 | Non-MeSH |
| D020754 | ATM | 472 | MH |
| D020754 | SACS | 26278 | MH |
| D020754 | PLEKHG4 | 25894 | MH |
| D020754 | SLC33A1 | 9197 | MH |
| D020754 | ATXN2 | 6311 | MH |
| D020754 | TTBK2 | 146057 | MH |
| D020754 | MARS2 | 92935 | MH |
| D020754 | SYNE1 | 23345 | MH |
| D020754 | AFG3L2 | 10939 | MH |
| D020754 | ATXN10 | 25814 | MH |
| D020754 | ITPR1 | 3708 | MH |
| D020754 | BEAN1 | 146227 | MH |
| D020754 | POLG | 5428 | MH |
| D020754 | TBP | 6908 | MH |
| D020754 | PPP2R2B | 5521 | MH |
| D020754 | SPTBN2 | 6712 | MH |
| D020754 | ATXN7 | 6314 | MH |
| D020754 | ATP2B3 | 492 | MH |
| D020754 | ATXN1 | 6310 | MH |
| D020754 | CACNA1A | 773 | MH |
| D020754 | TPP1 | 1200 | MH |
| D020754 | ATXN3 | 4287 | MH |
| D020754 | MTPAP | 55149 | MH |
| D020754 | SACS | 26278 | SCR |
| D020754 | TTBK2 | 146057 | SCR |
| D020754 | MARS2 | 92935 | SCR |
| D020754 | SYNE1 | 23345 | SCR |
| D020754 | PRKCG | 5582 | SCR |
| D020754 | AFG3L2 | 10939 | SCR |
| D020754 | SETX | 23064 | SCR |
| D020754 | ATXN10 | 25814 | SCR |
| D020754 | ITPR1 | 3708 | SCR |
| D020754 | BEAN1 | 146227 | SCR |
| D020754 | POLG | 5428 | SCR |
| D020754 | TBP | 6908 | SCR |
| D020754 | ATP2B3 | 492 | SCR |
| D020754 | PPP2R2B | 5521 | SCR |
| D020754 | KCNC3 | 3748 | SCR |
| D020754 | TPP1 | 1200 | SCR |
| D020754 | MTPAP | 55149 | SCR |
| D020754 | NOP56 | 10528 | Non-MeSH |
| D020754 | ANO10 | 55129 | Non-MeSH |
| D052177 | PKHD1 | 5314 | MH |
| D052177 | RPGRIP1L | 23322 | MH |
| D052177 | GLIS2 | 84662 | MH |
| D052177 | INPP5E | 56623 | MH |
| D052177 | HNF1B | 6928 | MH |
| D052177 | NPHP3 | 27031 | MH |
| D052177 | AHI1 | 54806 | MH |
| D052177 | NPHP4 | 261734 | MH |
| D052177 | ANKS6 | 203286 | MH |
| D052177 | CSPP1 | 79848 | MH |
| D052177 | INVS | 27130 | MH |
| D052177 | TMEM216 | 51259 | MH |
| D052177 | UMOD | 7369 | MH |
| D052177 | LMLN | 89782 | MH |
| D052177 | IQCB1 | 9657 | MH |
| D052177 | CEP290 | 80184 | MH |
| D052177 | NPHP1 | 4867 | MH |
| D052177 | PKD1 | 5310 | MH |
| D052177 | NXPH1 | 30010 | MH |
| D052177 | NPHP4 | 261734 | SCR |
| D052177 | LMLN | 89782 | SCR |
| D052177 | INVS | 27130 | SCR |
| D052177 | CSPP1 | 79848 | SCR |
| D052177 | HNF1B | 6928 | SCR |
| D052177 | RPGRIP1L | 23322 | SCR |
| D052177 | IQCB1 | 9657 | SCR |
| D052177 | TMEM216 | 51259 | SCR |
| D052177 | UMOD | 7369 | SCR |
| D052177 | CEP290 | 80184 | SCR |
| D052177 | AHI1 | 54806 | SCR |
| D052177 | NPHP1 | 4867 | SCR |
| D052177 | GLIS2 | 84662 | SCR |
| D052177 | INPP5E | 56623 | SCR |
| D052177 | NXPH1 | 30010 | SCR |
| D052177 | TCTN1 | 79600 | Non-MeSH |
| D052177 | ANKS6 | 203286 | Non-MeSH |
| D052177 | TMEM237 | 65062 | Non-MeSH |
| D052177 | SDCCAG8 | 10806 | Non-MeSH |
| D052177 | NPHP3 | 27031 | Non-MeSH |
| D052177 | CEP41 | 95681 | Non-MeSH |
| D052177 | KIF7 | 374654 | Non-MeSH |
| D052177 | TMEM138 | 51524 | Non-MeSH |
| D010009 | GYPE | 2996 | MH |
| D010009 | COL9A3 | 1299 | MH |
| D010009 | DGCR2 | 9993 | MH |
| D010009 | MATN3 | 4148 | MH |
| D010009 | COL9A1 | 1297 | MH |
| D010009 | SCN8A | 6334 | MH |
| D010009 | SLC26A1 | 10861 | MH |
| D010009 | SOST | 50964 | MH |
| D010009 | HSPG2 | 3339 | MH |
| D010009 | COMP | 1311 | MH |
| D010009 | SLC26A2 | 1836 | MH |
| D010009 | FLNA | 2316 | MH |
| D010009 | CLMP | 79827 | MH |
| D010009 | COL9A2 | 1298 | MH |
| D010009 | OBP2A | 29991 | SCR |
| D010009 | LIFR | 3977 | SCR |
| D010009 | MMP9 | 4318 | SCR |
| D010009 | SHOX | 6473 | SCR |
| D010009 | EIF2AK3 | 9451 | SCR |
| D010009 | COL2A1 | 1280 | SCR |
| D010009 | SLC26A2 | 1836 | SCR |
| D010009 | PAPSS2 | 9060 | SCR |
| D010009 | CLMP | 79827 | SCR |
| D010009 | CDKN1C | 1028 | SCR |
| D010009 | MMP13 | 4322 | SCR |
| D010009 | COL11A2 | 1302 | SCR |
| D010009 | SH3PXD2B | 285590 | SCR |
| D010009 | PNPLA6 | 10908 | SCR |
| D010009 | GPC6 | 10082 | SCR |
| D010009 | KIF22 | 3835 | SCR |
| D010009 | RMRP | 6023 | SCR |
| D010009 | ACP5 | 54 | SCR |
| D010009 | SLC39A13 | 91252 | SCR |
| D010009 | ACAN | 176 | SCR |
| D010009 | TREM2 | 54209 | SCR |
| D010009 | RNU4ATAC | 100151683 | SCR |
| D010009 | SMC2 | 10592 | SCR |
| D010009 | PCNT | 5116 | SCR |
| D010009 | TYROBP | 7305 | SCR |
| D010009 | COL11A1 | 1301 | SCR |
| D010009 | GYPE | 2996 | SCR |
| D010009 | FGFR1 | 2260 | SCR |
| D010009 | GDF5 | 8200 | SCR |
| D010009 | CIB2 | 10518 | SCR |
| D010009 | SCT | 6343 | SCR |
| D010009 | COL10A1 | 1300 | SCR |
| D010009 | SOST | 50964 | SCR |
| D010009 | SLC26A1 | 10861 | SCR |
| D010009 | CXCL17 | 284340 | SCR |
| D010009 | CSPG4 | 1464 | SCR |
| D010009 | DDR2 | 4921 | SCR |
| D010009 | ABCC9 | 10060 | SCR |
| D010009 | TBCE | 6905 | SCR |
| D010009 | RAB33B | 83452 | SCR |
| D010009 | ANKS4B | 257629 | SCR |
| D010009 | TKT | 7086 | SCR |
| D010009 | CENPJ | 55835 | SCR |
| D010009 | SLC25A12 | 8604 | SCR |
| D010009 | NKX3-2 | 579 | SCR |
| D010009 | SMARCAL1 | 50485 | SCR |
| D010009 | CHST3 | 9469 | SCR |
| D010009 | INPPL1 | 3636 | SCR |
| D010009 | H19 | 283120 | SCR |
| D010009 | SLC35D1 | 23169 | SCR |
| D010009 | FLNB | 2317 | SCR |
| D010009 | PRKAR1A | 5573 | SCR |
| D010009 | PTH1R | 5745 | SCR |
| D010009 | FLNA | 2316 | SCR |
| D010009 | DYM | 54808 | SCR |
| D010009 | COL2A1 | 1280 | Non-MeSH |
| D010009 | TRAPPC2 | 6399 | Non-MeSH |
| D010009 | GDF5 | 8200 | Non-MeSH |
| D003398 | RECQL4 | 9401 | MH |
| D003398 | CYP26B1 | 56603 | MH |
| D003398 | GJA1 | 2697 | MH |
| D003398 | MASP1 | 5648 | MH |
| D003398 | TWIST1 | 7291 | MH |
| D003398 | SKI | 6497 | MH |
| D003398 | FGFR1 | 2260 | MH |
| D003398 | IFT43 | 112752 | MH |
| D003398 | ASXL1 | 171023 | MH |
| D003398 | NUBP2 | 10101 | MH |
| D003398 | ERF | 2077 | MH |
| D003398 | IFT122 | 55764 | MH |
| D003398 | PANK2 | 80025 | MH |
| D003398 | FGFR2 | 2263 | MH |
| D003398 | MSX2 | 4488 | MH |
| D003398 | ENOSF1 | 55556 | MH |
| D003398 | FGFR3 | 2261 | MH |
| D003398 | PRDM5 | 11107 | MH |
| D003398 | TCF12 | 6938 | MH |
| D003398 | CRISP3 | 10321 | MH |
| D003398 | ALX4 | 60529 | MH |
| D003398 | FGFR1 | 2260 | SCR |
| D003398 | NUBP2 | 10101 | SCR |
| D003398 | RECQL4 | 9401 | SCR |
| D003398 | IFT122 | 55764 | SCR |
| D003398 | PANK2 | 80025 | SCR |
| D003398 | FGFR2 | 2263 | SCR |
| D003398 | ENOSF1 | 55556 | SCR |
| D003398 | MASP1 | 5648 | SCR |
| D003398 | SKI | 6497 | SCR |
| D003398 | MSX2 | 4488 | SCR |
| D003398 | FGFR3 | 2261 | SCR |
| D003398 | GJA1 | 2697 | SCR |
| D003398 | IFT43 | 112752 | SCR |
| D003398 | CRISP3 | 10321 | SCR |
| D003398 | ASXL1 | 171023 | SCR |
| D003398 | WDR35 | 57539 | Non-MeSH |
| D003398 | FREM1 | 158326 | Non-MeSH |
| D003398 | WDR19 | 57728 | Non-MeSH |
| D004677 | TCTN2 | 79867 | MH |
| D004677 | TMEM67 | 91147 | MH |
| D004677 | MSX2 | 4488 | MH |
| D004677 | RPGRIP1L | 23322 | MH |
| D004677 | TMEM216 | 51259 | MH |
| D004677 | NPHP3 | 27031 | MH |
| D004677 | B9D1 | 27077 | MH |
| D004677 | B9D2 | 80776 | MH |
| D004677 | CEP290 | 80184 | MH |
| D004677 | PRDM5 | 11107 | MH |
| D004677 | TMEM231 | 79583 | MH |
| D004677 | CC2D2A | 57545 | MH |
| D004677 | COL18A1 | 80781 | MH |
| D004677 | MKS1 | 54903 | MH |
| D004677 | ALX4 | 60529 | MH |
| D004677 | MSX2 | 4488 | SCR |
| D004677 | COL18A1 | 80781 | SCR |
| D004677 | PRDM5 | 11107 | SCR |
| D004677 | ALX4 | 60529 | SCR |
| D004677 | TCTN2 | 79867 | Non-MeSH |
| D004677 | TMEM67 | 91147 | Non-MeSH |
| D004677 | RPGRIP1L | 23322 | Non-MeSH |
| D004677 | TMEM216 | 51259 | Non-MeSH |
| D004677 | NPHP3 | 27031 | Non-MeSH |
| D004677 | B9D1 | 27077 | Non-MeSH |
| D004677 | CEP290 | 80184 | Non-MeSH |
| D004677 | TMEM231 | 79583 | Non-MeSH |
| D004677 | CC2D2A | 57545 | Non-MeSH |
| D004677 | B9D2 | 80776 | Non-MeSH |
| D004677 | MKS1 | 54903 | Non-MeSH |
| D002925 | SPAG1 | 6674 | MH |
| D002925 | RSPH1 | 89765 | MH |
| D002925 | RPGRIP1L | 23322 | MH |
| D002925 | DAB1 | 1600 | MH |
| D002925 | HYDIN | 54768 | MH |
| D002925 | ZMYND10 | 51364 | MH |
| D002925 | TCTN2 | 79867 | MH |
| D002925 | TMEM67 | 91147 | MH |
| D002925 | NPHP3 | 27031 | MH |
| D002925 | B9D1 | 27077 | MH |
| D002925 | MKS1 | 54903 | MH |
| D002925 | NME8 | 51314 | MH |
| D002925 | RSPH4A | 345895 | MH |
| D002925 | TMEM216 | 51259 | MH |
| D002925 | CCDC65 | 85478 | MH |
| D002925 | ARMC4 | 55130 | MH |
| D002925 | B9D2 | 80776 | MH |
| D002925 | DNAAF2 | 55172 | MH |
| D002925 | SPAG8 | 26206 | MH |
| D002925 | DNAAF3 | 352909 | MH |
| D002925 | DNAAF1 | 123872 | MH |
| D002925 | CEP290 | 80184 | MH |
| D002925 | TMEM231 | 79583 | MH |
| D002925 | CC2D2A | 57545 | MH |
| D002925 | RSPH9 | 221421 | MH |
| D002925 | DRC1 | 92749 | MH |
| D002925 | DNAAF1 | 123872 | SCR |
| D002925 | RSPH9 | 221421 | SCR |
| D002925 | DNAAF2 | 55172 | SCR |
| D002925 | RSPH4A | 345895 | SCR |
| D002925 | CCDC40 | 55036 | Non-MeSH |
| D002925 | TCTN2 | 79867 | Non-MeSH |
| D002925 | CCDC114 | 93233 | Non-MeSH |
| D002925 | TMEM67 | 91147 | Non-MeSH |
| D002925 | RPGRIP1L | 23322 | Non-MeSH |
| D002925 | TMEM216 | 51259 | Non-MeSH |
| D002925 | NPHP3 | 27031 | Non-MeSH |
| D002925 | B9D1 | 27077 | Non-MeSH |
| D002925 | CEP290 | 80184 | Non-MeSH |
| D002925 | DNAL1 | 83544 | Non-MeSH |
| D002925 | LRRC6 | 23639 | Non-MeSH |
| D002925 | TMEM231 | 79583 | Non-MeSH |
| D002925 | CC2D2A | 57545 | Non-MeSH |
| D002925 | B9D2 | 80776 | Non-MeSH |
| D002925 | MKS1 | 54903 | Non-MeSH |
| D002925 | CCDC103 | 388389 | Non-MeSH |
| D015419 | CYP7B1 | 9420 | MH |
| D015419 | B4GALNT1 | 2583 | MH |
| D015419 | SPG11 | 80208 | MH |
| D015419 | KIF1C | 10749 | MH |
| D015419 | SLC33A1 | 9197 | MH |
| D015419 | SPAST | 6683 | MH |
| D015419 | NIPA1 | 123606 | MH |
| D015419 | ALS2 | 57679 | MH |
| D015419 | PNPLA6 | 10908 | MH |
| D015419 | ZFYVE26 | 23503 | MH |
| D015419 | RTN2 | 6253 | MH |
| D015419 | KIF5C | 3800 | MH |
| D015419 | SPG7 | 6687 | MH |
| D015419 | BSCL2 | 26580 | MH |
| D015419 | SPG21 | 51324 | MH |
| D015419 | ATL1 | 51062 | MH |
| D015419 | ZFYVE27 | 118813 | MH |
| D015419 | AP4M1 | 9179 | MH |
| D015419 | HSPD1 | 3329 | MH |
| D015419 | PLP1 | 5354 | MH |
| D015419 | GJC2 | 57165 | MH |
| D015419 | ERLIN2 | 11160 | MH |
| D015419 | L1CAM | 3897 | MH |
| D015419 | REEP1 | 65055 | MH |
| D015419 | OPA1 | 4976 | MH |
| D015419 | KIF5A | 3798 | MH |
| D015419 | ENOPH1 | 58478 | MH |
| D015419 | CCT5 | 22948 | MH |
| D015419 | CYP7B1 | 9420 | SCR |
| D015419 | B4GALNT1 | 2583 | SCR |
| D015419 | SPG11 | 80208 | SCR |
| D015419 | KIF1C | 10749 | SCR |
| D015419 | SLC33A1 | 9197 | SCR |
| D015419 | SPAST | 6683 | SCR |
| D015419 | NIPA1 | 123606 | SCR |
| D015419 | ALS2 | 57679 | SCR |
| D015419 | PNPLA6 | 10908 | SCR |
| D015419 | ZFYVE26 | 23503 | SCR |
| D015419 | RTN2 | 6253 | SCR |
| D015419 | KIF5C | 3800 | SCR |
| D015419 | SPG7 | 6687 | SCR |
| D015419 | BSCL2 | 26580 | SCR |
| D015419 | SPG21 | 51324 | SCR |
| D015419 | ATL1 | 51062 | SCR |
| D015419 | ZFYVE27 | 118813 | SCR |
| D015419 | AP4M1 | 9179 | SCR |
| D015419 | HSPD1 | 3329 | SCR |
| D015419 | PLP1 | 5354 | SCR |
| D015419 | GJC2 | 57165 | SCR |
| D015419 | ERLIN2 | 11160 | SCR |
| D015419 | L1CAM | 3897 | SCR |
| D015419 | REEP1 | 65055 | SCR |
| D015419 | OPA1 | 4976 | SCR |
| D015419 | KIF5A | 3798 | SCR |
| D015419 | ENOPH1 | 58478 | SCR |
| D015419 | DDHD2 | 23259 | Non-MeSH |
| D015419 | TECPR2 | 9895 | Non-MeSH |
| D015419 | AP4S1 | 11154 | Non-MeSH |
| D015419 | AP5Z1 | 9907 | Non-MeSH |
| D015419 | C12orf65 | 91574 | Non-MeSH |
| D015419 | VPS37A | 137492 | Non-MeSH |
| D015419 | AP4E1 | 23431 | Non-MeSH |
| D015419 | CYP2U1 | 113612 | Non-MeSH |
| D003638 | MYO6 | 4646 | MH |
| D003638 | SDHD | 6392 | MH |
| D003638 | ATRX | 546 | MH |
| D003638 | COL2A1 | 1280 | MH |
| D003638 | PQBP1 | 10084 | MH |
| D003638 | IGF1 | 3479 | MH |
| D003638 | PTGDS | 5730 | MH |
| D003638 | KIF22 | 3835 | MH |
| D003638 | TYR | 7299 | MH |
| D003638 | KLKB1 | 3818 | MH |
| D003638 | GJB3 | 2707 | MH |
| D003638 | GJB6 | 10804 | MH |
| D003638 | SLC26A4 | 5172 | MH |
| D003638 | DNMT1 | 1786 | MH |
| D003638 | MPZL2 | 10205 | MH |
| D003638 | MITF | 4286 | MH |
| D003638 | BCAP31 | 10134 | MH |
| D003638 | GJB2 | 2706 | MH |
| D003638 | ITM2B | 9445 | MH |
| D003638 | TRMU | 55687 | MH |
| D003638 | SLITRK6 | 84189 | MH |
| D003638 | DSPP | 1834 | MH |
| D003638 | CHSY1 | 22856 | MH |
| D003638 | ATP2B2 | 491 | MH |
| D003638 | JAG1 | 182 | MH |
| D003638 | TNC | 3371 | MH |
| D003638 | TRNT1 | 51095 | MH |
| D003638 | ITM2B | 9445 | SCR |
| D003638 | KIF22 | 3835 | SCR |
| D003638 | TYR | 7299 | SCR |
| D003638 | CHSY1 | 22856 | SCR |
| D003638 | ATRX | 546 | SCR |
| D003638 | KLKB1 | 3818 | SCR |
| D003638 | MITF | 4286 | SCR |
| D003638 | PQBP1 | 10084 | SCR |
| D003638 | GJB2 | 2706 | SCR |
| D003638 | DIABLO | 56616 | Non-MeSH |
| D003638 | TJP2 | 9414 | Non-MeSH |
| D002311 | COTL1 | 23406 | MH |
| D002311 | TNNC1 | 7134 | MH |
| D002311 | TTN | 7273 | MH |
| D002311 | TCAP | 8557 | MH |
| D002311 | FKTN | 2218 | MH |
| D002311 | LAMA4 | 3910 | MH |
| D002311 | TNNI3 | 7137 | MH |
| D002311 | TMPO | 7112 | MH |
| D002311 | SEC61G | 23480 | MH |
| D002311 | RBM20 | 282996 | MH |
| D002311 | ROBO3 | 64221 | MH |
| D002311 | SOX9 | 6662 | MH |
| D002311 | PLN | 5350 | MH |
| D002311 | TBC1D8 | 11138 | MH |
| D002311 | NEXN | 91624 | MH |
| D002311 | PLB1 | 151056 | MH |
| D002311 | TNNT2 | 7139 | MH |
| D002311 | PRDM16 | 63976 | MH |
| D002311 | LAMA3 | 3909 | MH |
| D002311 | VCL | 7414 | MH |
| D002311 | DSP | 1832 | MH |
| D002311 | LDB3 | 11155 | MH |
| D002311 | MYPN | 84665 | MH |
| D002311 | LMNA | 4000 | MH |
| D002311 | EYA4 | 2070 | MH |
| D002311 | MYH6 | 4624 | MH |
| D002311 | SGCD | 6444 | MH |
| D002311 | ABCC9 | 10060 | MH |
| D002311 | CSRP3 | 8048 | MH |
| D002311 | SCN5A | 6331 | MH |
| D002311 | MYH7 | 4625 | MH |
| D002311 | DSG2 | 1829 | MH |
| D002311 | PSEN1 | 5663 | MH |
| D002311 | PSEN2 | 5664 | MH |
| D002311 | MYBPC3 | 4607 | MH |
| D002311 | TPM1 | 7168 | MH |
| D002311 | COTL1 | 23406 | SCR |
| D002311 | TCAP | 8557 | SCR |
| D002311 | TTN | 7273 | SCR |
| D002311 | TNNC1 | 7134 | SCR |
| D002311 | FKTN | 2218 | SCR |
| D002311 | TNNI3 | 7137 | SCR |
| D002311 | TMPO | 7112 | SCR |
| D002311 | SEC61G | 23480 | SCR |
| D002311 | RBM20 | 282996 | SCR |
| D002311 | ROBO3 | 64221 | SCR |
| D002311 | PLN | 5350 | SCR |
| D002311 | TBC1D8 | 11138 | SCR |
| D002311 | NEXN | 91624 | SCR |
| D002311 | PLB1 | 151056 | SCR |
| D002311 | TNNT2 | 7139 | SCR |
| D002311 | VCL | 7414 | SCR |
| D002311 | LMNA | 4000 | SCR |
| D002311 | EYA4 | 2070 | SCR |
| D002311 | SGCD | 6444 | SCR |
| D002311 | MYH6 | 4624 | SCR |
| D002311 | ABCC9 | 10060 | SCR |
| D002311 | CSRP3 | 8048 | SCR |
| D002311 | SCN5A | 6331 | SCR |
| D002311 | MYH7 | 4625 | SCR |
| D002311 | DSG2 | 1829 | SCR |
| D002311 | PSEN1 | 5663 | SCR |
| D002311 | PSEN2 | 5664 | SCR |
| D002311 | TPM1 | 7168 | SCR |
| D002311 | SDHA | 6389 | Non-MeSH |
| D002311 | ACTC1 | 70 | Non-MeSH |
| D002311 | SDHB | 6390 | Non-MeSH |
| D002292 | HNF1B | 6928 | MH |
| D002292 | SH2B3 | 10019 | MH |
| D002292 | PRCC | 5546 | MH |
| D002292 | DIRC2 | 84925 | MH |
| D002292 | FH | 2271 | MH |
| D002292 | RNF139 | 11236 | MH |
| D002292 | OGG1 | 4968 | MH |
| D002292 | UMOD | 7369 | MH |
| D002292 | VHL | 7428 | MH |
| D002292 | TFE3 | 7030 | MH |
| D002292 | HNF1A | 6927 | MH |
| D002292 | FLCN | 201163 | MH |
| D002292 | MET | 4233 | SCR |
| D002292 | HNF1B | 6928 | Non-MeSH |
| D002292 | SH2B3 | 10019 | Non-MeSH |
| D002292 | PRCC | 5546 | Non-MeSH |
| D002292 | DIRC2 | 84925 | Non-MeSH |
| D002292 | FH | 2271 | Non-MeSH |
| D002292 | RNF139 | 11236 | Non-MeSH |
| D002292 | OGG1 | 4968 | Non-MeSH |
| D002292 | UMOD | 7369 | Non-MeSH |
| D002292 | VHL | 7428 | Non-MeSH |
| D002292 | TFE3 | 7030 | Non-MeSH |
| D002292 | HNF1A | 6927 | Non-MeSH |
| D002292 | FLCN | 201163 | Non-MeSH |
| D007938 | MSTO1 | 55154 | MH |
| D007938 | ABL1 | 25 | MH |
| D007938 | NUMA1 | 4926 | MH |
| D007938 | RERE | 473 | MH |
| D007938 | GATA2 | 2624 | MH |
| D007938 | NPM1 | 4869 | MH |
| D007938 | BCL3 | 602 | MH |
| D007938 | BCR | 613 | MH |
| D007938 | RARA | 5914 | MH |
| D007938 | TCL1A | 8115 | MH |
| D007938 | ZBTB16 | 7704 | MH |
| D007938 | KDSR | 2531 | MH |
| D007938 | FLT3 | 2322 | MH |
| D007938 | TCL1B | 9623 | MH |
| D007938 | PML | 5371 | MH |
| D007938 | ABL2 | 27 | MH |
| D007938 | NQO1 | 1728 | MH |
| D007938 | STAT5B | 6777 | MH |
| D007938 | PDGFRA | 5156 | SCR |
| D007938 | LPP | 4026 | Non-MeSH |
| D007938 | ACSL5 | 51703 | Non-MeSH |
| D007938 | GNL3 | 26354 | Non-MeSH |
| D007938 | CHIC2 | 26511 | Non-MeSH |
| D007938 | KIT | 3815 | Non-MeSH |
| D007938 | ARNT | 405 | Non-MeSH |
| D007938 | NUP214 | 8021 | Non-MeSH |
| D007938 | IKZF1 | 10320 | Non-MeSH |
| D007938 | HOXD4 | 3233 | Non-MeSH |
| D007938 | NLRP2 | 55655 | Non-MeSH |
| D007938 | TERT | 7015 | Non-MeSH |
| D007938 | NPM1 | 4869 | Non-MeSH |
| D007938 | GMPS | 8833 | Non-MeSH |
| D007938 | PAX5 | 5079 | Non-MeSH |
| D007938 | BCR | 613 | Non-MeSH |
| D007938 | CPEB1 | 64506 | Non-MeSH |
| D007938 | JAK2 | 3717 | Non-MeSH |
| D007938 | MLLT10 | 8028 | Non-MeSH |
| D007938 | CABIN1 | 23523 | Non-MeSH |
| D007938 | SH3GL1 | 6455 | Non-MeSH |
| D007938 | SNAP91 | 9892 | Non-MeSH |
| D007938 | FLT3 | 2322 | Non-MeSH |
| D007938 | MLF1 | 4291 | Non-MeSH |
| D007938 | NME1 | 4830 | Non-MeSH |
| D007938 | KRAS | 3845 | Non-MeSH |
| D007938 | IRF1 | 3659 | Non-MeSH |
| D007938 | TCF3 | 6929 | Non-MeSH |
| D007938 | NSD1 | 64324 | Non-MeSH |
| D007938 | DEK | 7913 | Non-MeSH |
| D007938 | CEBPA | 1050 | Non-MeSH |
| D007938 | ARHGEF12 | 23365 | Non-MeSH |
| D007938 | LYL1 | 4066 | Non-MeSH |
| D007938 | PICALM | 8301 | Non-MeSH |
| D007938 | ACSL6 | 23305 | Non-MeSH |
| D007938 | RUNX1 | 861 | Non-MeSH |
| D003409 | NR1D1 | 9572 | MH |
| D003409 | COL11A1 | 1301 | MH |
| D003409 | NKX2-5 | 1482 | MH |
| D003409 | KAT6B | 23522 | MH |
| D003409 | SLC5A5 | 6528 | MH |
| D003409 | GLIS3 | 169792 | MH |
| D003409 | EZH1 | 2145 | MH |
| D003409 | EZH2 | 2146 | MH |
| D003409 | NFIX | 4784 | MH |
| D003409 | PAX8 | 7849 | MH |
| D003409 | TPO | 7173 | MH |
| D003409 | IYD | 389434 | MH |
| D003409 | THRA | 7067 | MH |
| D003409 | COL11A1 | 1301 | SCR |
| D003409 | EZH1 | 2145 | SCR |
| D003409 | SLC5A5 | 6528 | SCR |
| D003409 | EZH2 | 2146 | SCR |
| D003409 | GLIS3 | 169792 | SCR |
| D003409 | KAT6B | 23522 | SCR |
| D003409 | NFIX | 4784 | SCR |
| D003409 | PAX8 | 7849 | SCR |
| D003409 | TSHR | 7253 | SCR |
| D003409 | TPO | 7173 | SCR |
| D003409 | IYD | 389434 | SCR |
| D003409 | NR1D1 | 9572 | Non-MeSH |
| D003409 | NKX2-5 | 1482 | Non-MeSH |
| D003409 | THRA | 7067 | Non-MeSH |
| D000690 | ERBB4 | 2066 | MH |
| D000690 | FIG4 | 9896 | MH |
| D000690 | TARDBP | 23435 | MH |
| D000690 | ANG | 283 | MH |
| D000690 | C9orf72 | 203228 | MH |
| D000690 | CHMP2B | 25978 | MH |
| D000690 | HNRNPA1 | 3178 | MH |
| D000690 | SETX | 23064 | MH |
| D000690 | DCTN1 | 1639 | MH |
| D000690 | NEFH | 4744 | MH |
| D000690 | VAPB | 9217 | MH |
| D000690 | FUS | 2521 | MH |
| D000690 | TRPM7 | 54822 | MH |
| D000690 | ALS2 | 57679 | MH |
| D000690 | PRPH | 5630 | MH |
| D000690 | SOD1 | 6647 | MH |
| D000690 | FIG4 | 9896 | SCR |
| D000690 | TARDBP | 23435 | SCR |
| D000690 | ANG | 283 | SCR |
| D000690 | CHMP2B | 25978 | SCR |
| D000690 | DCTN1 | 1639 | SCR |
| D000690 | NEFH | 4744 | SCR |
| D000690 | SETX | 23064 | SCR |
| D000690 | VAPB | 9217 | SCR |
| D000690 | TRPM7 | 54822 | SCR |
| D000690 | ALS2 | 57679 | SCR |
| D000690 | PRPH | 5630 | SCR |
| D000690 | SOD1 | 6647 | SCR |
| D000690 | VCP | 7415 | Non-MeSH |
| D000690 | UBQLN2 | 29978 | Non-MeSH |
