## Supplementary table 4 for "pyMeSHSim: an integrative python package for biomedical named entity recognition, normalization and comparison"

**Supplementary table 4.** GWAS phenotypes parsed by Nelson's group and pyMeSHSim, and the semantic similarity between them calculated by pyMeSHSim and meshes.

| **GWAS phenotype** | **pyMeSHSim recognized term** | | | **Nelson's group recognized term** | | | **pyMeSHSim calculated similarity** | | | | | **meshes calculated similarity** | | | | |
| --- | --- | --- | --- | --- | --- | --- | --- | --- | --- | --- | --- | --- | --- | --- | --- | --- |
|  | **Term** | **ID** | **Category** | **Term** | **ID** | **Category** | **Lin** | **Res** | **Jiang** | **Rel** | **Wang** | **Lin** | **Res** | **Jiang** | **Rel** | **Wang** |
| Nephrolithiasis | Kidney Calculi | D007669 | C | Nephrolithiasis | D007669 | C | 1.00 | 1.00 | 1.00 | 1.00 | 1.00 | 1.00 | 1.00 | 1.00 | 1.00 | 1.00 |
| Age-related macular degeneration | Degeneration, Macular | D008268 | C | Macular degeneration | D008268 | C | 1.00 | 1.00 | 1.00 | 1.00 | 1.00 | 1.00 | 1.00 | 1.00 | 1.00 | 1.00 |
| Systemic lupus erythematosus | Lupus Erythematosus, Systemic | D008180 | C | Lupus erythematosus, systemic | D008180 | C | 1.00 | 1.00 | 1.00 | 1.00 | 1.00 | 1.00 | 1.00 | 1.00 | 1.00 | 1.00 |
| Type 1 diabetes autoantibodies | Type 1 Diabetes Mellitus | D003922 | C | Diabetes mellitus, type 1 | D003922 | C | 1.00 | 1.00 | 1.00 | 1.00 | 1.00 | 1.00 | 1.00 | 1.00 | 1.00 | 1.00 |
| Type 1 diabetes | Type 1 Diabetes Mellitus | D003922 | C | Diabetes mellitus, type 1 | D003922 | C | 1.00 | 1.00 | 1.00 | 1.00 | 1.00 | 1.00 | 1.00 | 1.00 | 1.00 | 1.00 |
| Systolic blood pressure | Blood Pressure | D001794 | G,E | Blood pressure | D001794 | G,E | 1.00 | 1.00 | 1.00 | 1.00 | 1.00 | 1.00 | 1.00 | 1.00 | 1.00 | 1.00 |
| Esophageal cancer | Esophageal Neoplasm | D004938 | C | Esophageal neoplasms | D004938 | C | 1.00 | 1.00 | 1.00 | 1.00 | 1.00 | 1.00 | 1.00 | 1.00 | 1.00 | 1.00 |
| Triglycerides | Triglycerides | D014280 | D | Triglycerides | D014280 | D | 1.00 | 1.00 | 1.00 | 1.00 | 1.00 | 1.00 | 1.00 | 1.00 | 1.00 | 1.00 |
| Crohn`s disease | Crohn Disease | D003424 | C | Crohn disease | D003424 | C | 1.00 | 1.00 | 1.00 | 1.00 | 1.00 | 1.00 | 1.00 | 1.00 | 1.00 | 1.00 |
| Breast cancer | Breast Neoplasms | D001943 | C | Breast neoplasms | D001943 | C | 1.00 | 1.00 | 1.00 | 1.00 | 1.00 | 1.00 | 1.00 | 1.00 | 1.00 | 1.00 |
| Corneal structure | Cornea | D003315 | A | Cornea | D003315 | A | 1.00 | 1.00 | 1.00 | 1.00 | 1.00 | 1.00 | 1.00 | 1.00 | 1.00 | 1.00 |
| Leprosy | Leprosies | D007918 | C | Leprosy | D007918 | C | 1.00 | 1.00 | 1.00 | 1.00 | 1.00 | 1.00 | 1.00 | 1.00 | 1.00 | 1.00 |
| Waist-hip ratio | Waist-to-Hip Ratio | D049629 | G,E | Waist-hip ratio | D049629 | G,E | 1.00 | 1.00 | 1.00 | 1.00 | 1.00 | 1.00 | 1.00 | 1.00 | 1.00 | 1.00 |
| Alzheimer`s disease | Alzheimer Disease | D000544 | C,F | Alzheimer disease | D000544 | F,C | 1.00 | 1.00 | 1.00 | 1.00 | 1.00 | 1.00 | 1.00 | 1.00 | 1.00 | 1.00 |
| Chronic Hepatitis C infection | Hepatitis C, Chronic | D019698 | C | Hepatitis c, chronic | D019698 | C | 1.00 | 1.00 | 1.00 | 1.00 | 1.00 | 1.00 | 1.00 | 1.00 | 1.00 | 1.00 |
| Amyotrophic lateral sclerosis | Amyotrophic Lateral Sclerosis | D000690 | C | Amyotrophic lateral sclerosis | D000690 | C | 1.00 | 1.00 | 1.00 | 1.00 | 1.00 | 1.00 | 1.00 | 1.00 | 1.00 | 1.00 |
| Bladder cancer | Urinary Bladder Neoplasms | D001749 | C | Urinary bladder neoplasms | D001749 | C | 1.00 | 1.00 | 1.00 | 1.00 | 1.00 | 1.00 | 1.00 | 1.00 | 1.00 | 1.00 |
| Glaucoma (primary open-angle) | Glaucoma, Open-Angle | D005902 | C | Glaucoma, open-angle | D005902 | C | 1.00 | 1.00 | 1.00 | 1.00 | 1.00 | 1.00 | 1.00 | 1.00 | 1.00 | 1.00 |
| Freckles | Melanoses | D008548 | C | Melanosis | D008548 | C | 1.00 | 1.00 | 1.00 | 1.00 | 1.00 | 1.00 | 1.00 | 1.00 | 1.00 | 1.00 |
| Hematocrit | Hematocrit | D006400 | G,E | Hematocrit | D006400 | G,E | 1.00 | 1.00 | 1.00 | 1.00 | 1.00 | 1.00 | 1.00 | 1.00 | 1.00 | 1.00 |
| Fibrinogen | Fibrinogen | D005340 | D | Fibrinogen | D005340 | D | 1.00 | 1.00 | 1.00 | 1.00 | 1.00 | 1.00 | 1.00 | 1.00 | 1.00 | 1.00 |
| Crohn`s disease and Celiac disease | Celiac Disease | D002446 | C | Celiac disease | D002446 | C | 1.00 | 1.00 | 1.00 | 1.00 | 1.00 | 1.00 | 1.00 | 1.00 | 1.00 | 1.00 |
| Venous thromboembolism | Venous Thromboembolism | D054556 | C | Venous thromboembolism | D054556 | C | 1.00 | 1.00 | 1.00 | 1.00 | 1.00 | 1.00 | 1.00 | 1.00 | 1.00 | 1.00 |
| Hypertension | Hypertension | D006973 | C | Hypertension | D006973 | C | 1.00 | 1.00 | 1.00 | 1.00 | 1.00 | 1.00 | 1.00 | 1.00 | 1.00 | 1.00 |
| Pericardial fat | Pericardium | D010496 | A | Pericardium | D010496 | A | 1.00 | 1.00 | 1.00 | 1.00 | 1.00 | 1.00 | 1.00 | 1.00 | 1.00 | 1.00 |
| Cardiovascular disease risk factors | Cardiovascular Disease | D002318 | C | Cardiovascular disease | D002318 | C | 1.00 | 1.00 | 1.00 | 1.00 | 1.00 | 1.00 | 1.00 | 1.00 | 1.00 | 1.00 |
| Bone mineral density | Bone Densities | D015519 | G | Bone density | D015519 | G | 1.00 | 1.00 | 1.00 | 1.00 | 1.00 | 1.00 | 1.00 | 1.00 | 1.00 | 1.00 |
| Testicular germ cell cancer | Neoplasm, Testicular | D013736 | C | Testicular neoplasms | D013736 | C | 1.00 | 1.00 | 1.00 | 1.00 | 1.00 | 1.00 | 1.00 | 1.00 | 1.00 | 1.00 |
| Hemoglobin | Hemoglobins | D006454 | D | Hemoglobins | D006454 | D | 1.00 | 1.00 | 1.00 | 1.00 | 1.00 | 1.00 | 1.00 | 1.00 | 1.00 | 1.00 |
| Alopecia areata | Alopecia Areata | D000506 | C | Alopecia areata | D000506 | C | 1.00 | 1.00 | 1.00 | 1.00 | 1.00 | 1.00 | 1.00 | 1.00 | 1.00 | 1.00 |
| Graves` disease | Disease, Graves' | D006111 | C | Graves disease | D006111 | C | 1.00 | 1.00 | 1.00 | 1.00 | 1.00 | 1.00 | 1.00 | 1.00 | 1.00 | 1.00 |
| C-reactive protein and white blood cell count | C-Reactive Protein | D002097 | D | C-reactive protein | D002097 | D | 1.00 | 1.00 | 1.00 | 1.00 | 1.00 | 1.00 | 1.00 | 1.00 | 1.00 | 1.00 |
| Sagittal craniosynostosis | Craniosynostoses | D003398 | C | Craniosynostoses | D003398 | C | 1.00 | 1.00 | 1.00 | 1.00 | 1.00 | 1.00 | 1.00 | 1.00 | 1.00 | 1.00 |
| Pancreatitis | Pancreatitides | D010195 | C | Pancreatitis | D010195 | C | 1.00 | 1.00 | 1.00 | 1.00 | 1.00 | 1.00 | 1.00 | 1.00 | 1.00 | 1.00 |
| Myopia (pathological) | Degenerative Myopias | D047728 | C | Myopia, degenerative | D047728 | C | 1.00 | 1.00 | 1.00 | 1.00 | 1.00 | 1.00 | 1.00 | 1.00 | 1.00 | 1.00 |
| Prostate cancer | Neoplasm, Prostatic | D011471 | C | Prostatic neoplasms | D011471 | C | 1.00 | 1.00 | 1.00 | 1.00 | 1.00 | 1.00 | 1.00 | 1.00 | 1.00 | 1.00 |
| Longevity | Longevity | D008136 | G | Longevity | D008136 | G | 1.00 | 1.00 | 1.00 | 1.00 | 1.00 | 1.00 | 1.00 | 1.00 | 1.00 | 1.00 |
| Colorectal cancer | Colorectal Neoplasm | D015179 | C | Colorectal neoplasms | D015179 | C | 1.00 | 1.00 | 1.00 | 1.00 | 1.00 | 1.00 | 1.00 | 1.00 | 1.00 | 1.00 |
| Inflammatory bowel disease | Inflammatory Bowel Diseases | D015212 | C | Inflammatory bowel diseases | D015212 | C | 1.00 | 1.00 | 1.00 | 1.00 | 1.00 | 1.00 | 1.00 | 1.00 | 1.00 | 1.00 |
| Birth weight | Birth Weight | D001724 | G,E,C | Birth weight | D001724 | G,E,C | 1.00 | 1.00 | 1.00 | 1.00 | 1.00 | 1.00 | 1.00 | 1.00 | 1.00 | 1.00 |
| Erythrocyte sedimentation rate | Blood Sedimentation | D001799 | E | Blood sedimentation | D001799 | E | 1.00 | 1.00 | 1.00 | 1.00 | 1.00 | 1.00 | 1.00 | 1.00 | 1.00 | 1.00 |
| Response to statin therapy | Hydroxymethylglutaryl CoA Reductase Inhibitors | D019161 | D | Hydroxymethylglutaryl-coa reductase inhibitors | D019161 | D | 1.00 | 1.00 | 1.00 | 1.00 | 1.00 | 1.00 | 1.00 | 1.00 | 1.00 | 1.00 |
| Lipoprotein-associated phospholipase A2 activity change in response to statin therapy | 1 Alkyl 2 acetylglycerophosphocholine Esterase | D043203 | D | 1-alkyl-2-acetylglycerophosphocholine esterase | D043203 | D | 1.00 | 1.00 | 1.00 | 1.00 | 1.00 | 1.00 | 1.00 | 1.00 | 1.00 | 1.00 |
| Response to statin therapy (LDL-C) | Hydroxymethylglutaryl CoA Reductase Inhibitors | D019161 | D | Hydroxymethylglutaryl-coa reductase inhibitors | D019161 | D | 1.00 | 1.00 | 1.00 | 1.00 | 1.00 | 1.00 | 1.00 | 1.00 | 1.00 | 1.00 |
| Multiple sclerosis | Multiple Sclerosis | D009103 | C | Multiple sclerosis | D009103 | C | 1.00 | 1.00 | 1.00 | 1.00 | 1.00 | 1.00 | 1.00 | 1.00 | 1.00 | 1.00 |
| Glycated hemoglobin levels | Glycated Hemoglobin A | D006442 | D | Hemoglobin a, glycosylated | D006442 | D | 1.00 | 1.00 | 1.00 | 1.00 | 1.00 | 1.00 | 1.00 | 1.00 | 1.00 | 1.00 |
| Homocysteine levels | Homocystine | D006711 | D | Homocysteine | D006711 | D | 1.00 | 1.00 | 1.00 | 1.00 | 1.00 | 1.00 | 1.00 | 1.00 | 1.00 | 1.00 |
| Rheumatoid arthritis | Arthritis, Rheumatoid | D001172 | C | Arthritis, rheumatoid | D001172 | C | 1.00 | 1.00 | 1.00 | 1.00 | 1.00 | 1.00 | 1.00 | 1.00 | 1.00 | 1.00 |
| Primary biliary cirrhosis | Liver Cirrhoses, Biliary | D008105 | C | Liver cirrhosis, biliary | D008105 | C | 1.00 | 1.00 | 1.00 | 1.00 | 1.00 | 1.00 | 1.00 | 1.00 | 1.00 | 1.00 |
| Systemic sclerosis | Scleroderma, Systemic | D012595 | C | Scleroderma, systemic | D012595 | C | 1.00 | 1.00 | 1.00 | 1.00 | 1.00 | 1.00 | 1.00 | 1.00 | 1.00 | 1.00 |
| Schizophrenia | Schizophrenias | D012559 | F | Schizophrenia | D012559 | F | 1.00 | 1.00 | 1.00 | 1.00 | 1.00 | 1.00 | 1.00 | 1.00 | 1.00 | 1.00 |
| HDL cholesterol | Cholesterol, HDL | D008076 | D | Cholesterol, hdl | D008076 | D | 1.00 | 1.00 | 1.00 | 1.00 | 1.00 | 1.00 | 1.00 | 1.00 | 1.00 | 1.00 |
| Age-related macular degeneration (GA) | Degeneration, Macular | D008268 | C | Macular degeneration | D008268 | C | 1.00 | 1.00 | 1.00 | 1.00 | 1.00 | 1.00 | 1.00 | 1.00 | 1.00 | 1.00 |
| Age-related macular degeneration (CNV vs. GA) | Degeneration, Macular | D008268 | C | Macular degeneration | D008268 | C | 1.00 | 1.00 | 1.00 | 1.00 | 1.00 | 1.00 | 1.00 | 1.00 | 1.00 | 1.00 |
| Alcohol consumption | Alcohol Drinking | D000428 | F | Alcohol drinking | D000428 | F | 1.00 | 1.00 | 1.00 | 1.00 | 1.00 | 1.00 | 1.00 | 1.00 | 1.00 | 1.00 |
| Electrocardiographic traits | Electrocardiography | D004562 | E | Electrocardiography | D004562 | E | 1.00 | 1.00 | 1.00 | 1.00 | 1.00 | 1.00 | 1.00 | 1.00 | 1.00 | 1.00 |
| Paget`s disease | Osteitis Deformans | D010001 | C | Osteitis deformans | D010001 | C | 1.00 | 1.00 | 1.00 | 1.00 | 1.00 | 1.00 | 1.00 | 1.00 | 1.00 | 1.00 |
| Proinsulin levels | Proinsulin | D011384 | D | Proinsulin | D011384 | D | 1.00 | 1.00 | 1.00 | 1.00 | 1.00 | 1.00 | 1.00 | 1.00 | 1.00 | 1.00 |
| Primary tooth development (time to first tooth eruption) | Tooth, Deciduous | D014094 | A | Dentition, primary | D014094 | A | 1.00 | 1.00 | 1.00 | 1.00 | 1.00 | 1.00 | 1.00 | 1.00 | 1.00 | 1.00 |
| Acenocoumarol maintenance dosage | Acenocoumarol | D000074 | D | Acenocoumarol | D000074 | D | 1.00 | 1.00 | 1.00 | 1.00 | 1.00 | 1.00 | 1.00 | 1.00 | 1.00 | 1.00 |
| Nicotine dependence | Disorder, Tobacco Use | D014029 | C,F | Tobacco use disorder | D014029 | F,C | 1.00 | 1.00 | 1.00 | 1.00 | 1.00 | 1.00 | 1.00 | 1.00 | 1.00 | 1.00 |
| Thyroid hormone levels | Hormones, Thyroid | D013963 | D | Thyroid hormones | D013963 | D | 1.00 | 1.00 | 1.00 | 1.00 | 1.00 | 1.00 | 1.00 | 1.00 | 1.00 | 1.00 |
| C-reactive protein | C-Reactive Protein | D002097 | D | C-reactive protein | D002097 | D | 1.00 | 1.00 | 1.00 | 1.00 | 1.00 | 1.00 | 1.00 | 1.00 | 1.00 | 1.00 |
| Warfarin maintenance dose | Warfarin | D014859 | D | Warfarin | D014859 | D | 1.00 | 1.00 | 1.00 | 1.00 | 1.00 | 1.00 | 1.00 | 1.00 | 1.00 | 1.00 |
| Lung cancer | Lung Neoplasm | D008175 | C | Lung neoplasms | D008175 | C | 1.00 | 1.00 | 1.00 | 1.00 | 1.00 | 1.00 | 1.00 | 1.00 | 1.00 | 1.00 |
| Coagulation factor levels | Blood Coagulation Factors | D001779 | D | Blood coagulation factors | D001779 | D | 1.00 | 1.00 | 1.00 | 1.00 | 1.00 | 1.00 | 1.00 | 1.00 | 1.00 | 1.00 |
| Insulin-like growth factors | Somatomedins | D013002 | D | Somatomedins | D013002 | D | 1.00 | 1.00 | 1.00 | 1.00 | 1.00 | 1.00 | 1.00 | 1.00 | 1.00 | 1.00 |
| End-stage coagulation | Blood Coagulation | D001777 | G | Blood coagulation | D001777 | G | 1.00 | 1.00 | 1.00 | 1.00 | 1.00 | 1.00 | 1.00 | 1.00 | 1.00 | 1.00 |
| Coronary artery disease | Artery Diseases, Coronary | D003324 | C | Coronary artery disease | D003324 | C | 1.00 | 1.00 | 1.00 | 1.00 | 1.00 | 1.00 | 1.00 | 1.00 | 1.00 | 1.00 |
| Ovarian cancer | Neoplasm, Ovarian | D010051 | C | Ovarian neoplasms | D010051 | C | 1.00 | 1.00 | 1.00 | 1.00 | 1.00 | 1.00 | 1.00 | 1.00 | 1.00 | 1.00 |
| Mean corpuscular volume | Erythrocyte Indices | D004909 | G,E | Erythrocyte indices | D004909 | G,E | 1.00 | 1.00 | 1.00 | 1.00 | 1.00 | 1.00 | 1.00 | 1.00 | 1.00 | 1.00 |
| Mean corpuscular hemoglobin | Erythrocyte Indices | D004909 | G,E | Erythrocyte indices | D004909 | G,E | 1.00 | 1.00 | 1.00 | 1.00 | 1.00 | 1.00 | 1.00 | 1.00 | 1.00 | 1.00 |
| Sudden cardiac arrest | Cardiac Death, Sudden | D016757 | C | Death, sudden, cardiac | D016757 | C | 1.00 | 1.00 | 1.00 | 1.00 | 1.00 | 1.00 | 1.00 | 1.00 | 1.00 | 1.00 |
| Vitiligo | Vitiligo | D014820 | C | Vitiligo | D014820 | C | 1.00 | 1.00 | 1.00 | 1.00 | 1.00 | 1.00 | 1.00 | 1.00 | 1.00 | 1.00 |
| Psoriasis | Psoriases | D011565 | C | Psoriasis | D011565 | C | 1.00 | 1.00 | 1.00 | 1.00 | 1.00 | 1.00 | 1.00 | 1.00 | 1.00 | 1.00 |
| Brain structure | Brain | D001921 | A | Brain | D001921 | A | 1.00 | 1.00 | 1.00 | 1.00 | 1.00 | 1.00 | 1.00 | 1.00 | 1.00 | 1.00 |
| Metabolic syndrome | Metabolic Syndrome | D024821 | C | Metabolic syndrome x | D024821 | C | 1.00 | 1.00 | 1.00 | 1.00 | 1.00 | 1.00 | 1.00 | 1.00 | 1.00 | 1.00 |
| Waist Circumference - Triglycerides (WC-TG) | Waist Circumferences | D055105 | G,E | Waist circumference | D055105 | G,E | 1.00 | 1.00 | 1.00 | 1.00 | 1.00 | 1.00 | 1.00 | 1.00 | 1.00 | 1.00 |
| Ulcerative colitis | Colitis, Ulcerative | D003093 | C | Colitis, ulcerative | D003093 | C | 1.00 | 1.00 | 1.00 | 1.00 | 1.00 | 1.00 | 1.00 | 1.00 | 1.00 | 1.00 |
| Celiac disease | Celiac Disease | D002446 | C | Celiac disease | D002446 | C | 1.00 | 1.00 | 1.00 | 1.00 | 1.00 | 1.00 | 1.00 | 1.00 | 1.00 | 1.00 |
| Type 2 diabetes | Type 2 Diabetes Mellitus | D003924 | C | Diabetes mellitus, type 2 | D003924 | C | 1.00 | 1.00 | 1.00 | 1.00 | 1.00 | 1.00 | 1.00 | 1.00 | 1.00 | 1.00 |
| Polycystic ovary syndrome | Ovary Syndrome, Polycystic | D011085 | C | Polycystic ovary syndrome | D011085 | C | 1.00 | 1.00 | 1.00 | 1.00 | 1.00 | 1.00 | 1.00 | 1.00 | 1.00 | 1.00 |
| Atrial fibrillation | Atrial Fibrillation | D001281 | C | Atrial fibrillation | D001281 | C | 1.00 | 1.00 | 1.00 | 1.00 | 1.00 | 1.00 | 1.00 | 1.00 | 1.00 | 1.00 |
| Acute lymphoblastic leukemia (childhood) | Precursor Cell Lymphoblastic Leukemia-Lymphoma | D054198 | C | Precursor cell lymphoblastic leukemia-lymphoma | D054198 | C | 1.00 | 1.00 | 1.00 | 1.00 | 1.00 | 1.00 | 1.00 | 1.00 | 1.00 | 1.00 |
| Androgen levels | Androgens | D000728 | D | Androgens | D000728 | D | 1.00 | 1.00 | 1.00 | 1.00 | 1.00 | 1.00 | 1.00 | 1.00 | 1.00 | 1.00 |
| Metabolic syndrome (bivariate traits) | Metabolic Syndrome | D024821 | C | Metabolic syndrome x | D024821 | C | 1.00 | 1.00 | 1.00 | 1.00 | 1.00 | 1.00 | 1.00 | 1.00 | 1.00 | 1.00 |
| Waist circumference | Waist Circumferences | D055105 | G,E | Waist circumference | D055105 | G,E | 1.00 | 1.00 | 1.00 | 1.00 | 1.00 | 1.00 | 1.00 | 1.00 | 1.00 | 1.00 |
| Endometriosis | Endometrioses | D004715 | C | Endometriosis | D004715 | C | 1.00 | 1.00 | 1.00 | 1.00 | 1.00 | 1.00 | 1.00 | 1.00 | 1.00 | 1.00 |
| Cholesterol, total | Cholesterol | D002784 | D | Cholesterol | D002784 | D | 1.00 | 1.00 | 1.00 | 1.00 | 1.00 | 1.00 | 1.00 | 1.00 | 1.00 | 1.00 |
| Celiac disease and Rheumatoid arthritis | Arthritis, Rheumatoid | D001172 | C | Arthritis, rheumatoid | D001172 | C | 1.00 | 1.00 | 1.00 | 1.00 | 1.00 | 1.00 | 1.00 | 1.00 | 1.00 | 1.00 |
| Malaria | Malaria | D008288 | C | Malaria | D008288 | C | 1.00 | 1.00 | 1.00 | 1.00 | 1.00 | 1.00 | 1.00 | 1.00 | 1.00 | 1.00 |
| Dilated cardiomyopathy | Cardiomyopathies, Dilated | D002311 | C | Cardiomyopathy, dilated | D002311 | C | 1.00 | 1.00 | 1.00 | 1.00 | 1.00 | 1.00 | 1.00 | 1.00 | 1.00 | 1.00 |
| Neuroblastoma (high-risk) | Neuroblastoma | D009447 | C | Neuroblastoma | D009447 | C | 1.00 | 1.00 | 1.00 | 1.00 | 1.00 | 1.00 | 1.00 | 1.00 | 1.00 | 1.00 |
| Drug-induced liver injury (flucloxacillin) | Floxacillin | D005436 | D | Floxacillin | D005436 | D | 1.00 | 1.00 | 1.00 | 1.00 | 1.00 | 1.00 | 1.00 | 1.00 | 1.00 | 1.00 |
| Intracranial aneurysm | Aneurysms, Intracranial | D002532 | C | Intracranial aneurysm | D002532 | C | 1.00 | 1.00 | 1.00 | 1.00 | 1.00 | 1.00 | 1.00 | 1.00 | 1.00 | 1.00 |
| Myeloproliferative neoplasms | Disorder, Myeloproliferative | D009196 | C | Myeloproliferative disorders | D009196 | C | 1.00 | 1.00 | 1.00 | 1.00 | 1.00 | 1.00 | 1.00 | 1.00 | 1.00 | 1.00 |
| Thyroid cancer | Neoplasm, Thyroid | D013964 | C | Thyroid neoplasms | D013964 | C | 1.00 | 1.00 | 1.00 | 1.00 | 1.00 | 1.00 | 1.00 | 1.00 | 1.00 | 1.00 |
| Bipolar disorder | Bipolar Disorders | D001714 | F | Bipolar disorder | D001714 | F | 1.00 | 1.00 | 1.00 | 1.00 | 1.00 | 1.00 | 1.00 | 1.00 | 1.00 | 1.00 |
| Atopic dermatitis | Atopic Dermatitides | D003876 | C | Dermatitis, atopic | D003876 | C | 1.00 | 1.00 | 1.00 | 1.00 | 1.00 | 1.00 | 1.00 | 1.00 | 1.00 | 1.00 |
| Immune reponse to smallpox (secreted IL-1beta) | Variola virus | D012901 | B | Smallpox | D012901 | B | 1.00 | 1.00 | 1.00 | 1.00 | 1.00 | 1.00 | 1.00 | 1.00 | 1.00 | 1.00 |
| Asthma | Asthma | D001249 | C | Asthma | D001249 | C | 1.00 | 1.00 | 1.00 | 1.00 | 1.00 | 1.00 | 1.00 | 1.00 | 1.00 | 1.00 |
| Neuroblastoma | Neuroblastoma | D009447 | C | Neuroblastoma | D009447 | C | 1.00 | 1.00 | 1.00 | 1.00 | 1.00 | 1.00 | 1.00 | 1.00 | 1.00 | 1.00 |
| Chronic lymphocytic leukemia | Leukemia, Lymphocytic, Chronic, B-Cell | D015451 | C | Leukemia, lymphocytic, chronic, b-cell | D015451 | C | 1.00 | 1.00 | 1.00 | 1.00 | 1.00 | 1.00 | 1.00 | 1.00 | 1.00 | 1.00 |
| Adiponectin levels | Adiponectin | D052242 | D | Adiponectin | D052242 | D | 1.00 | 1.00 | 1.00 | 1.00 | 1.00 | 1.00 | 1.00 | 1.00 | 1.00 | 1.00 |
| Digit length ratio | Finger | D005385 | A | Fingers | D005385 | A | 1.00 | 1.00 | 1.00 | 1.00 | 1.00 | 1.00 | 1.00 | 1.00 | 1.00 | 1.00 |
| Immune reponse to smallpox (secreted IFN-alpha) | Variola virus | D012901 | B | Smallpox | D012901 | B | 1.00 | 1.00 | 1.00 | 1.00 | 1.00 | 1.00 | 1.00 | 1.00 | 1.00 | 1.00 |
| Migraine | Disorder, Migraine | D008881 | C | Migraine disorders | D008881 | C | 1.00 | 1.00 | 1.00 | 1.00 | 1.00 | 1.00 | 1.00 | 1.00 | 1.00 | 1.00 |
| Scoliosis | Scolioses | D012600 | C | Scoliosis | D012600 | C | 1.00 | 1.00 | 1.00 | 1.00 | 1.00 | 1.00 | 1.00 | 1.00 | 1.00 | 1.00 |
| Blood pressure | Blood Pressure | D001794 | G,E | Blood pressure | D001794 | G,E | 1.00 | 1.00 | 1.00 | 1.00 | 1.00 | 1.00 | 1.00 | 1.00 | 1.00 | 1.00 |
| Age-related macular degeneration (wet) | Degeneration, Macular | D008268 | C | Wet macular degeneration | D008268 | C | 1.00 | 1.00 | 1.00 | 1.00 | 1.00 | 1.00 | 1.00 | 1.00 | 1.00 | 1.00 |
| Ankylosing spondylitis | Spondylitis, Ankylosing | D013167 | C | Spondylitis, ankylosing | D013167 | C | 1.00 | 1.00 | 1.00 | 1.00 | 1.00 | 1.00 | 1.00 | 1.00 | 1.00 | 1.00 |
| Response to metformin | Metformin | D008687 | D | Metformin | D008687 | D | 1.00 | 1.00 | 1.00 | 1.00 | 1.00 | 1.00 | 1.00 | 1.00 | 1.00 | 1.00 |
| Immune reponse to smallpox (secreted IL-2) | Variola virus | D012901 | B | Smallpox | D012901 | B | 1.00 | 1.00 | 1.00 | 1.00 | 1.00 | 1.00 | 1.00 | 1.00 | 1.00 | 1.00 |
| Serum matrix metalloproteinase | Matrix Metalloproteinases | D020782 | D | Matrix metalloproteinases | D020782 | D | 1.00 | 1.00 | 1.00 | 1.00 | 1.00 | 1.00 | 1.00 | 1.00 | 1.00 | 1.00 |
| Myocardial infarction (early onset) | Infarctions, Myocardial | D009203 | C | Myocardial infarction | D009203 | C | 1.00 | 1.00 | 1.00 | 1.00 | 1.00 | 1.00 | 1.00 | 1.00 | 1.00 | 1.00 |
| Coronary heart disease | Artery Diseases, Coronary | D003324 | C | Coronary disease | D003324 | C | 1.00 | 1.00 | 1.00 | 1.00 | 1.00 | 1.00 | 1.00 | 1.00 | 1.00 | 1.00 |
| Parkinson`s disease | Parkinson Disease | D010300 | C | Parkinson disease | D010300 | C | 1.00 | 1.00 | 1.00 | 1.00 | 1.00 | 1.00 | 1.00 | 1.00 | 1.00 | 1.00 |
| Ribavirin-induced anemia | Anemia | D000740 | C | Anemia | D000740 | C | 1.00 | 1.00 | 1.00 | 1.00 | 1.00 | 1.00 | 1.00 | 1.00 | 1.00 | 1.00 |
| Cystatin C | Cystatin C | D055316 | D | Cystatin c | D055316 | D | 1.00 | 1.00 | 1.00 | 1.00 | 1.00 | 1.00 | 1.00 | 1.00 | 1.00 | 1.00 |
| LDL cholesterol | Cholesterol, LDL | D008078 | D | Cholesterol, ldl | D008078 | D | 1.00 | 1.00 | 1.00 | 1.00 | 1.00 | 1.00 | 1.00 | 1.00 | 1.00 | 1.00 |
| Tanning | Tanning | D013633 | J | Tanning | D013633 | J | 1.00 | 1.00 | 1.00 | 1.00 | 1.00 | 1.00 | 1.00 | 1.00 | 1.00 | 1.00 |
| Serum butyrylcholinesterase | Cholinesterases | D002802 | D | Cholinesterases | D002802 | D | 1.00 | 1.00 | 1.00 | 1.00 | 1.00 | 1.00 | 1.00 | 1.00 | 1.00 | 1.00 |
| Response to hepatitis C treatment | Hepatitis C | D006526 | C | Hepatitis c | D006526 | C | 1.00 | 1.00 | 1.00 | 1.00 | 1.00 | 1.00 | 1.00 | 1.00 | 1.00 | 1.00 |
| Serum dehydroepiandrosterone sulphate levels | Dehydroepiandrosterone Sulfate | D019314 | D | Dehydroepiandrosterone sulphate | D019314 | D | 1.00 | 1.00 | 1.00 | 1.00 | 1.00 | 1.00 | 1.00 | 1.00 | 1.00 | 1.00 |
| Response to fenofibrate | Fenofibrate | D011345 | D | Fenofibrate | D011345 | D | 1.00 | 1.00 | 1.00 | 1.00 | 1.00 | 1.00 | 1.00 | 1.00 | 1.00 | 1.00 |
| Alzheimer`s disease (late onset) | Alzheimer Disease | D000544 | C,F | Alzheimer disease | D000544 | F,C | 1.00 | 1.00 | 1.00 | 1.00 | 1.00 | 1.00 | 1.00 | 1.00 | 1.00 | 1.00 |
| AIDS progression | Acquired Immunodeficiency Syndrome | D000163 | C | Acquired immunodeficiency syndrome | D000163 | C | 1.00 | 1.00 | 1.00 | 1.00 | 1.00 | 1.00 | 1.00 | 1.00 | 1.00 | 1.00 |
| Fetal hemoglobin levels | Fetal Hemoglobin | D005319 | D | Fetal hemoglobin | D005319 | D | 1.00 | 1.00 | 1.00 | 1.00 | 1.00 | 1.00 | 1.00 | 1.00 | 1.00 | 1.00 |
| Gallstones | Gallstone | D042882 | C | Gallstones | D042882 | C | 1.00 | 1.00 | 1.00 | 1.00 | 1.00 | 1.00 | 1.00 | 1.00 | 1.00 | 1.00 |
| Bilirubin levels | Bilirubin | D001663 | D | Bilirubin | D001663 | D | 1.00 | 1.00 | 1.00 | 1.00 | 1.00 | 1.00 | 1.00 | 1.00 | 1.00 | 1.00 |
| Stroke | Strokes | D020521 | C | Stroke | D020521 | C | 1.00 | 1.00 | 1.00 | 1.00 | 1.00 | 1.00 | 1.00 | 1.00 | 1.00 | 1.00 |
| Bone mineral density (hip) | Bone Densities | D015519 | G | Bone density | D015519 | G | 1.00 | 1.00 | 1.00 | 1.00 | 1.00 | 1.00 | 1.00 | 1.00 | 1.00 | 1.00 |
| Obesity-related traits | Obesity | D009765 | G,E,C | Obesity | D009765 | G,E,C | 1.00 | 1.00 | 1.00 | 1.00 | 1.00 | 1.00 | 1.00 | 1.00 | 1.00 | 1.00 |
| White blood cell count | Count, Leukocyte | D007958 | G,E | Leukocyte count | D007958 | G,E | 1.00 | 1.00 | 1.00 | 1.00 | 1.00 | 1.00 | 1.00 | 1.00 | 1.00 | 1.00 |
| Sex hormone-binding globulin levels | Globulin, Sex Hormone-Binding | D012738 | D | Sex hormone-binding globulin | D012738 | D | 1.00 | 1.00 | 1.00 | 1.00 | 1.00 | 1.00 | 1.00 | 1.00 | 1.00 | 1.00 |
| Testosterone levels | Testosterone | D013739 | D | Testosterone | D013739 | D | 1.00 | 1.00 | 1.00 | 1.00 | 1.00 | 1.00 | 1.00 | 1.00 | 1.00 | 1.00 |
| Progressive supranuclear palsy | Progressive Supranuclear Palsies | D013494 | C | Supranuclear palsy, progressive | D013494 | C | 1.00 | 1.00 | 1.00 | 1.00 | 1.00 | 1.00 | 1.00 | 1.00 | 1.00 | 1.00 |
| Diabetic retinopathy | Diabetic Retinopathies | D003930 | C | Diabetic retinopathy | D003930 | C | 1.00 | 1.00 | 1.00 | 1.00 | 1.00 | 1.00 | 1.00 | 1.00 | 1.00 | 1.00 |
| Squamous cell carcinoma | Carcinoma, Squamous Cell | D002294 | C | Carcinoma, squamous cell | D002294 | C | 1.00 | 1.00 | 1.00 | 1.00 | 1.00 | 1.00 | 1.00 | 1.00 | 1.00 | 1.00 |
| Prostate-specific antigen levels | Prostate-Specific Antigen | D017430 | D | Prostate-specific antigen | D017430 | D | 1.00 | 1.00 | 1.00 | 1.00 | 1.00 | 1.00 | 1.00 | 1.00 | 1.00 | 1.00 |
| Aging | Aging | D000375 | G | Aging | D000375 | G | 1.00 | 1.00 | 1.00 | 1.00 | 1.00 | 1.00 | 1.00 | 1.00 | 1.00 | 1.00 |
| Uterine fibroids | Leiomyoma | D007889 | C | Leiomyoma | D007889 | C | 1.00 | 1.00 | 1.00 | 1.00 | 1.00 | 1.00 | 1.00 | 1.00 | 1.00 | 1.00 |
| Crohn`s disease and psoriasis | Psoriases | D011565 | C | Psoriasis | D011565 | C | 1.00 | 1.00 | 1.00 | 1.00 | 1.00 | 1.00 | 1.00 | 1.00 | 1.00 | 1.00 |
| Inflammatory bowel disease (early onset) | Inflammatory Bowel Diseases | D015212 | C | Inflammatory bowel diseases | D015212 | C | 1.00 | 1.00 | 1.00 | 1.00 | 1.00 | 1.00 | 1.00 | 1.00 | 1.00 | 1.00 |
| IgA nephropathy | Glomerulonephritides, IGA | D005922 | C | Glomerulonephritis, iga | D005922 | C | 1.00 | 1.00 | 1.00 | 1.00 | 1.00 | 1.00 | 1.00 | 1.00 | 1.00 | 1.00 |
| Gamma glutamyl transpeptidase | gamma Glutamyltransferase | D005723 | D | Gamma-glutamyltransferase | D005723 | D | 1.00 | 1.00 | 1.00 | 1.00 | 1.00 | 1.00 | 1.00 | 1.00 | 1.00 | 1.00 |
| Male-pattern baldness | Alopecia | D000505 | C | Alopecia | D000505 | C | 1.00 | 1.00 | 1.00 | 1.00 | 1.00 | 1.00 | 1.00 | 1.00 | 1.00 | 1.00 |
| Restless legs syndrome | Restless Legs Syndrome | D012148 | C,F | Restless legs syndrome | D012148 | F,C | 1.00 | 1.00 | 1.00 | 1.00 | 1.00 | 1.00 | 1.00 | 1.00 | 1.00 | 1.00 |
| Hypertriglyceridemia | Hypertriglyceridemia | D015228 | C | Hypertriglyceridemia | D015228 | C | 1.00 | 1.00 | 1.00 | 1.00 | 1.00 | 1.00 | 1.00 | 1.00 | 1.00 | 1.00 |
| Telomere length | Telomere | D016615 | A,G | Telomere | D016615 | G,A | 1.00 | 1.00 | 1.00 | 1.00 | 1.00 | 1.00 | 1.00 | 1.00 | 1.00 | 1.00 |
| Cystic fibrosis severity | Cystic Fibrosis | D003550 | C | Cystic fibrosis | D003550 | C | 1.00 | 1.00 | 1.00 | 1.00 | 1.00 | 1.00 | 1.00 | 1.00 | 1.00 | 1.00 |
| Renal function and chronic kidney disease | Chronic Renal Insufficiency | D051436 | C | Renal insufficiency, chronic | D051436 | C | 1.00 | 1.00 | 1.00 | 1.00 | 1.00 | 1.00 | 1.00 | 1.00 | 1.00 | 1.00 |
| Psoriatic arthritis | Arthritis, Psoriatic | D015535 | C | Arthritis, psoriatic | D015535 | C | 1.00 | 1.00 | 1.00 | 1.00 | 1.00 | 1.00 | 1.00 | 1.00 | 1.00 | 1.00 |
| Immune reponse to smallpox (secreted IL-10) | Variola virus | D012901 | B | Smallpox | D012901 | B | 1.00 | 1.00 | 1.00 | 1.00 | 1.00 | 1.00 | 1.00 | 1.00 | 1.00 | 1.00 |
| Uric acid levels | Acid, Uric | D014527 | D | Uric acid | D014527 | D | 1.00 | 1.00 | 1.00 | 1.00 | 1.00 | 1.00 | 1.00 | 1.00 | 1.00 | 1.00 |
| Osteoporosis | Osteoporoses | D010024 | C | Osteoporosis | D010024 | C | 1.00 | 1.00 | 1.00 | 1.00 | 1.00 | 1.00 | 1.00 | 1.00 | 1.00 | 1.00 |
| Gastric cancer | Neoplasm, Stomach | D013274 | C | Stomach neoplasms | D013274 | C | 1.00 | 1.00 | 1.00 | 1.00 | 1.00 | 1.00 | 1.00 | 1.00 | 1.00 | 1.00 |
| Melanoma | Melanoma | D008545 | C | Melanoma | D008545 | C | 1.00 | 1.00 | 1.00 | 1.00 | 1.00 | 1.00 | 1.00 | 1.00 | 1.00 | 1.00 |
| Obesity | Obesity | D009765 | G,E,C | Obesity | D009765 | G,E,C | 1.00 | 1.00 | 1.00 | 1.00 | 1.00 | 1.00 | 1.00 | 1.00 | 1.00 | 1.00 |
| Abdominal aortic aneurysm | Aortic Aneurysm, Abdominal | D017544 | C | Aortic aneurysm, abdominal | D017544 | C | 1.00 | 1.00 | 1.00 | 1.00 | 1.00 | 1.00 | 1.00 | 1.00 | 1.00 | 1.00 |
| Gout | Gout | D006073 | C | Gout | D006073 | C | 1.00 | 1.00 | 1.00 | 1.00 | 1.00 | 1.00 | 1.00 | 1.00 | 1.00 | 1.00 |
| Behcet`s disease | Behcet Syndrome | D001528 | C | Behcet syndrome | D001528 | C | 1.00 | 1.00 | 1.00 | 1.00 | 1.00 | 1.00 | 1.00 | 1.00 | 1.00 | 1.00 |
| Keloid | Keloid | D007627 | A,C | Keloid | D007627 | A,C | 1.00 | 1.00 | 1.00 | 1.00 | 1.00 | 1.00 | 1.00 | 1.00 | 1.00 | 1.00 |
| Periodontitis | Periodontitides | D010518 | C | Periodontitis | D010518 | C | 1.00 | 1.00 | 1.00 | 1.00 | 1.00 | 1.00 | 1.00 | 1.00 | 1.00 | 1.00 |
| Pancreatic cancer | Neoplasm, Pancreatic | D010190 | C | Pancreatic neoplasms | D010190 | C | 1.00 | 1.00 | 1.00 | 1.00 | 1.00 | 1.00 | 1.00 | 1.00 | 1.00 | 1.00 |
| Activated partial thromboplastin time | Partial Thromboplastin Time | D010314 | G,E | Partial thromboplastin time | D010314 | G,E | 1.00 | 1.00 | 1.00 | 1.00 | 1.00 | 1.00 | 1.00 | 1.00 | 1.00 | 1.00 |
| Recombination rate (females) | Genetic Recombinations | D011995 | G | Recombination, genetic | D011995 | G | 1.00 | 1.00 | 1.00 | 1.00 | 1.00 | 1.00 | 1.00 | 1.00 | 1.00 | 1.00 |
| Conduct disorder (symptom count) | Conduct Disorder | D019955 | F | Conduct disorder | D019955 | F | 1.00 | 1.00 | 1.00 | 1.00 | 1.00 | 1.00 | 1.00 | 1.00 | 1.00 | 1.00 |
| Bone mineral density (spine) | Bone Densities | D015519 | G | Bone density | D015519 | G | 1.00 | 1.00 | 1.00 | 1.00 | 1.00 | 1.00 | 1.00 | 1.00 | 1.00 | 1.00 |
| Biliary atresia | Biliary Atresia | D001656 | C | Biliary atresia | D001656 | C | 1.00 | 1.00 | 1.00 | 1.00 | 1.00 | 1.00 | 1.00 | 1.00 | 1.00 | 1.00 |
| Aspartate aminotransferase | Aminotransferase, Aspartate | D001219 | D | Aspartate aminotransferases | D001219 | D | 1.00 | 1.00 | 1.00 | 1.00 | 1.00 | 1.00 | 1.00 | 1.00 | 1.00 | 1.00 |
| Major depressive disorder (broad) | Depressive Disorders | D003866 | F | Depressive disorder | D003866 | F | 1.00 | 1.00 | 1.00 | 1.00 | 1.00 | 1.00 | 1.00 | 1.00 | 1.00 | 1.00 |
| Chronic obstructive pulmonary disease-related biomarkers | Pulmonary Disease, Chronic Obstructive | D029424 | C | Pulmonary disease, chronic obstructive | D029424 | C | 1.00 | 1.00 | 1.00 | 1.00 | 1.00 | 1.00 | 1.00 | 1.00 | 1.00 | 1.00 |
| Hepatocellular carcinoma | Carcinoma, Hepatocellular | D006528 | C | Carcinoma, hepatocellular | D006528 | C | 1.00 | 1.00 | 1.00 | 1.00 | 1.00 | 1.00 | 1.00 | 1.00 | 1.00 | 1.00 |
| Resting heart rate | Heart Rate | D006339 | G,E | Heart rate | D006339 | G,E | 1.00 | 1.00 | 1.00 | 1.00 | 1.00 | 1.00 | 1.00 | 1.00 | 1.00 | 1.00 |
| Soluble leptin receptor levels | Receptors, Leptin | D054411 | D | Receptors, leptin | D054411 | D | 1.00 | 1.00 | 1.00 | 1.00 | 1.00 | 1.00 | 1.00 | 1.00 | 1.00 | 1.00 |
| Soluble ICAM-1 | Intercellular Adhesion Molecule 1 | D018799 | D | Intercellular adhesion molecule-1 | D018799 | D | 1.00 | 1.00 | 1.00 | 1.00 | 1.00 | 1.00 | 1.00 | 1.00 | 1.00 | 1.00 |
| Creutzfeldt-Jakob disease | Creutzfeldt Jakob Syndrome | D007562 | C,F | Creutzfeldt-jakob syndrome | D007562 | F,C | 1.00 | 1.00 | 1.00 | 1.00 | 1.00 | 1.00 | 1.00 | 1.00 | 1.00 | 1.00 |
| Kawasaki disease | Mucocutaneous Lymph Node Syndrome | D009080 | C | Mucocutaneous lymph node syndrome | D009080 | C | 1.00 | 1.00 | 1.00 | 1.00 | 1.00 | 1.00 | 1.00 | 1.00 | 1.00 | 1.00 |
| Basal cell carcinoma (cutaneous) | Basal Cell Carcinomas | D002280 | C | Carcinoma, basal cell | D002280 | C | 1.00 | 1.00 | 1.00 | 1.00 | 1.00 | 1.00 | 1.00 | 1.00 | 1.00 | 1.00 |
| Interleukin-18 levels | Interleukin 18 | D020382 | D | Interleukin-18 | D020382 | D | 1.00 | 1.00 | 1.00 | 1.00 | 1.00 | 1.00 | 1.00 | 1.00 | 1.00 | 1.00 |
| Carotid intima media thickness | Carotid Intima-Media Thickness | D059168 | G,E | Carotid intima-media thickness | D059168 | G,E | 1.00 | 1.00 | 1.00 | 1.00 | 1.00 | 1.00 | 1.00 | 1.00 | 1.00 | 1.00 |
| Protein quantitative trait loci | Loci, Quantitative Trait | D040641 | G | Quantitative trait loci | D040641 | G | 1.00 | 1.00 | 1.00 | 1.00 | 1.00 | 1.00 | 1.00 | 1.00 | 1.00 | 1.00 |
| Optic disc parameters | Disk, Optic | D009898 | A | Optic disk | D009898 | A | 1.00 | 1.00 | 1.00 | 1.00 | 1.00 | 1.00 | 1.00 | 1.00 | 1.00 | 1.00 |
| Chronic obstructive pulmonary disease | Pulmonary Disease, Chronic Obstructive | D029424 | C | Pulmonary disease, chronic obstructive | D029424 | C | 1.00 | 1.00 | 1.00 | 1.00 | 1.00 | 1.00 | 1.00 | 1.00 | 1.00 | 1.00 |
| Haptoglobin levels | Haptoglobins | D006242 | D | Haptoglobins | D006242 | D | 1.00 | 1.00 | 1.00 | 1.00 | 1.00 | 1.00 | 1.00 | 1.00 | 1.00 | 1.00 |
| Hodgkin`s lymphoma | Disease, Hodgkin | D006689 | C | Hodgkin disease | D006689 | C | 1.00 | 1.00 | 1.00 | 1.00 | 1.00 | 1.00 | 1.00 | 1.00 | 1.00 | 1.00 |
| Beta thalassemia/hemoglobin E disease | beta-Thalassemia | D017086 | C | Beta-thalassemia | D017086 | C | 1.00 | 1.00 | 1.00 | 1.00 | 1.00 | 1.00 | 1.00 | 1.00 | 1.00 | 1.00 |
| Stroke (ischemic) | Strokes | D020521 | C | Stroke | D020521 | C | 1.00 | 1.00 | 1.00 | 1.00 | 1.00 | 1.00 | 1.00 | 1.00 | 1.00 | 1.00 |
| Kidney stones | Kidney Calculi | D007669 | C | Kidney calculi | D007669 | C | 1.00 | 1.00 | 1.00 | 1.00 | 1.00 | 1.00 | 1.00 | 1.00 | 1.00 | 1.00 |
| Plasminogen activator inhibitor type 1 levels (PAI-1) | Plasminogen Activator Inhibitor 1 | D017395 | D | Plasminogen activator inhibitor 1 | D017395 | D | 1.00 | 1.00 | 1.00 | 1.00 | 1.00 | 1.00 | 1.00 | 1.00 | 1.00 | 1.00 |
| Soluble levels of adhesion molecules | Adhesion Molecules, Cell | D015815 | D | Cell adhesion molecules | D015815 | D | 1.00 | 1.00 | 1.00 | 1.00 | 1.00 | 1.00 | 1.00 | 1.00 | 1.00 | 1.00 |
| Ewing sarcoma | Sarcoma, Ewings | D012512 | C | Sarcoma, ewing | D012512 | C | 1.00 | 1.00 | 1.00 | 1.00 | 1.00 | 1.00 | 1.00 | 1.00 | 1.00 | 1.00 |
| Major mood disorders | Mood Disorders | D019964 | F | Mood disorders | D019964 | F | 1.00 | 1.00 | 1.00 | 1.00 | 1.00 | 1.00 | 1.00 | 1.00 | 1.00 | 1.00 |
| Mean corpuscular hemoglobin concentration | Erythrocyte Indices | D004909 | G,E | Erythrocyte indices | D004909 | G,E | 1.00 | 1.00 | 1.00 | 1.00 | 1.00 | 1.00 | 1.00 | 1.00 | 1.00 | 1.00 |
| Protein biomarker | Biomarkers | D015415 | D | Biological markers | D015415 | D | 1.00 | 1.00 | 1.00 | 1.00 | 1.00 | 1.00 | 1.00 | 1.00 | 1.00 | 1.00 |
| Wilms tumor | Wilm Tumor | D009396 | C | Wilms tumor | D009396 | C | 1.00 | 1.00 | 1.00 | 1.00 | 1.00 | 1.00 | 1.00 | 1.00 | 1.00 | 1.00 |
| Multiple cancers (lung cancer, gastric cancer, and squamous cell carcinoma) | Neoplasms | D009369 | C | Neoplasms | D009369 | C | 1.00 | 1.00 | 1.00 | 1.00 | 1.00 | 1.00 | 1.00 | 1.00 | 1.00 | 1.00 |
| Duodenal ulcer | Duodenal Ulcer | D004381 | C | Duodenal ulcer | D004381 | C | 1.00 | 1.00 | 1.00 | 1.00 | 1.00 | 1.00 | 1.00 | 1.00 | 1.00 | 1.00 |
| Prothrombin time | Prothrombin Time | D011517 | G,E | Prothrombin time | D011517 | G,E | 1.00 | 1.00 | 1.00 | 1.00 | 1.00 | 1.00 | 1.00 | 1.00 | 1.00 | 1.00 |
| Glioma | Gliomas | D005910 | C | Glioma | D005910 | C | 1.00 | 1.00 | 1.00 | 1.00 | 1.00 | 1.00 | 1.00 | 1.00 | 1.00 | 1.00 |
| Nephropathy | Diseases, Kidney | D007674 | C | Kidney diseases | D007674 | C | 1.00 | 1.00 | 1.00 | 1.00 | 1.00 | 1.00 | 1.00 | 1.00 | 1.00 | 1.00 |
| Coffee consumption | Coffee | D003069 | J,G,D | Coffee | D003069 | J,G,D | 1.00 | 1.00 | 1.00 | 1.00 | 1.00 | 1.00 | 1.00 | 1.00 | 1.00 | 1.00 |
| Select biomarker traits | Biomarkers | D015415 | D | Biological markers | D015415 | D | 1.00 | 1.00 | 1.00 | 1.00 | 1.00 | 1.00 | 1.00 | 1.00 | 1.00 | 1.00 |
| Pain | Pain | D010146 | G,C,F | Pain | D010146 | G,F,C | 1.00 | 1.00 | 1.00 | 1.00 | 1.00 | 1.00 | 1.00 | 1.00 | 1.00 | 1.00 |
| Amyloid A Levels | Serum Amyloid Protein A | D000685 | D | Serum amyloid a protein | D000685 | D | 1.00 | 1.00 | 1.00 | 1.00 | 1.00 | 1.00 | 1.00 | 1.00 | 1.00 | 1.00 |
| Epirubicin-induced leukopenia | Leukopenia | D007970 | C | Leukopenia | D007970 | C | 1.00 | 1.00 | 1.00 | 1.00 | 1.00 | 1.00 | 1.00 | 1.00 | 1.00 | 1.00 |
| Primary sclerosing cholangitis | Cholangitides, Sclerosing | D015209 | C | Cholangitis, sclerosing | D015209 | C | 1.00 | 1.00 | 1.00 | 1.00 | 1.00 | 1.00 | 1.00 | 1.00 | 1.00 | 1.00 |
| Glaucoma | Glaucomas | D005901 | C | Glaucoma | D005901 | C | 1.00 | 1.00 | 1.00 | 1.00 | 1.00 | 1.00 | 1.00 | 1.00 | 1.00 | 1.00 |
| Smoking behavior | Smoking | D012907 | F | Smoking | D012907 | F | 1.00 | 1.00 | 1.00 | 1.00 | 1.00 | 1.00 | 1.00 | 1.00 | 1.00 | 1.00 |
| Waist circumference and related phenotypes | Waist Circumferences | D055105 | G,E | Waist circumference | D055105 | G,E | 1.00 | 1.00 | 1.00 | 1.00 | 1.00 | 1.00 | 1.00 | 1.00 | 1.00 | 1.00 |
| Dengue shock syndrome | Severe Dengue | D019595 | C | Dengue hemorrhagic fever | D019595 | C | 1.00 | 1.00 | 1.00 | 1.00 | 1.00 | 1.00 | 1.00 | 1.00 | 1.00 | 1.00 |
| Serum total protein level | Blood Proteins | D001798 | D | Blood proteins | D001798 | D | 1.00 | 1.00 | 1.00 | 1.00 | 1.00 | 1.00 | 1.00 | 1.00 | 1.00 | 1.00 |
| Recombination rate (males) | Genetic Recombinations | D011995 | G | Recombination, genetic | D011995 | G | 1.00 | 1.00 | 1.00 | 1.00 | 1.00 | 1.00 | 1.00 | 1.00 | 1.00 | 1.00 |
| Breast cancer (male) | Breast Neoplasm, Male | D018567 | C | Breast neoplasms, male | D018567 | C | 1.00 | 1.00 | 1.00 | 1.00 | 1.00 | 1.00 | 1.00 | 1.00 | 1.00 | 1.00 |
| Eosinophilic esophagitis (pediatric) | Esophagitides, Eosinophilic | D057765 | C | Eosinophilic esophagitis | D057765 | C | 1.00 | 1.00 | 1.00 | 1.00 | 1.00 | 1.00 | 1.00 | 1.00 | 1.00 | 1.00 |
| Plasma C4b binding protein levels | C4b-Binding Protein, Complement | D050716 | D | Complement c4b-binding protein | D050716 | D | 1.00 | 1.00 | 1.00 | 1.00 | 1.00 | 1.00 | 1.00 | 1.00 | 1.00 | 1.00 |
| Glaucoma (exfoliation) | Exfoliation Syndrome | D017889 | C | Exfoliation syndrome | D017889 | C | 1.00 | 1.00 | 1.00 | 1.00 | 1.00 | 1.00 | 1.00 | 1.00 | 1.00 | 1.00 |
| Resistin levels | Resistin | D052243 | D | Resistin | D052243 | D | 1.00 | 1.00 | 1.00 | 1.00 | 1.00 | 1.00 | 1.00 | 1.00 | 1.00 | 1.00 |
| Dental caries | Dental Caries | D003731 | C | Dental caries | D003731 | C | 1.00 | 1.00 | 1.00 | 1.00 | 1.00 | 1.00 | 1.00 | 1.00 | 1.00 | 1.00 |
| Serum Phytosterol Levels | Phytosterols | D010840 | D | Phytosterols | D010840 | D | 1.00 | 1.00 | 1.00 | 1.00 | 1.00 | 1.00 | 1.00 | 1.00 | 1.00 | 1.00 |
| Methotrexate clearance (acute lymphoblastic leukemia) | Methotrexate | D008727 | D | Methotrexate | D008727 | D | 1.00 | 1.00 | 1.00 | 1.00 | 1.00 | 1.00 | 1.00 | 1.00 | 1.00 | 1.00 |
| Alzheimer`s disease (age of onset) | Alzheimer Disease | D000544 | C,F | Alzheimer disease | D000544 | F,C | 1.00 | 1.00 | 1.00 | 1.00 | 1.00 | 1.00 | 1.00 | 1.00 | 1.00 | 1.00 |
| Endometrial cancer | Endometrial Neoplasm | D016889 | C | Endometrial neoplasms | D016889 | C | 1.00 | 1.00 | 1.00 | 1.00 | 1.00 | 1.00 | 1.00 | 1.00 | 1.00 | 1.00 |
| Testicular cancer | Neoplasm, Testicular | D013736 | C | Testicular neoplasms | D013736 | C | 1.00 | 1.00 | 1.00 | 1.00 | 1.00 | 1.00 | 1.00 | 1.00 | 1.00 | 1.00 |
| Multiple myeloma | Multiple Myeloma | D009101 | C | Multiple myeloma | D009101 | C | 1.00 | 1.00 | 1.00 | 1.00 | 1.00 | 1.00 | 1.00 | 1.00 | 1.00 | 1.00 |
| Nephropathy (idiopathic membranous) | Glomerulonephritides, Membranous | D015433 | C | Glomerulonephritis, membranous | D015433 | C | 1.00 | 1.00 | 1.00 | 1.00 | 1.00 | 1.00 | 1.00 | 1.00 | 1.00 | 1.00 |
| Chronic myeloid leukemia | Leukemia, Myelogenous, Chronic, BCR-ABL Positive | D015464 | C | Leukemia, myelogenous, chronic, bcr-abl positive | D015464 | C | 1.00 | 1.00 | 1.00 | 1.00 | 1.00 | 1.00 | 1.00 | 1.00 | 1.00 | 1.00 |
| Sarcoidosis | Sarcoidoses | D012507 | C | Sarcoidosis | D012507 | C | 1.00 | 1.00 | 1.00 | 1.00 | 1.00 | 1.00 | 1.00 | 1.00 | 1.00 | 1.00 |
| Factor VII | Factor VII | D005167 | D | Factor vii | D005167 | D | 1.00 | 1.00 | 1.00 | 1.00 | 1.00 | 1.00 | 1.00 | 1.00 | 1.00 | 1.00 |
| Coronary disease | Artery Diseases, Coronary | D003324 | C | Coronary disease | D003324 | C | 1.00 | 1.00 | 1.00 | 1.00 | 1.00 | 1.00 | 1.00 | 1.00 | 1.00 | 1.00 |
| Intelligence | Intelligence | D007360 | F | Intelligence | D007360 | F | 1.00 | 1.00 | 1.00 | 1.00 | 1.00 | 1.00 | 1.00 | 1.00 | 1.00 | 1.00 |
| Fuch`s corneal dystrophy | Dystrophy, Fuchs' Endothelial | D005642 | C | Fuchs' endothelial dystrophy | D005642 | C | 1.00 | 1.00 | 1.00 | 1.00 | 1.00 | 1.00 | 1.00 | 1.00 | 1.00 | 1.00 |
| Fuchs`s corneal dystrophy | Dystrophy, Fuchs' Endothelial | D005642 | C | Fuchs' endothelial dystrophy | D005642 | C | 1.00 | 1.00 | 1.00 | 1.00 | 1.00 | 1.00 | 1.00 | 1.00 | 1.00 | 1.00 |
| Plasma E-selectin levels | E-Selectin | D019040 | D | E-selectin | D019040 | D | 1.00 | 1.00 | 1.00 | 1.00 | 1.00 | 1.00 | 1.00 | 1.00 | 1.00 | 1.00 |
| Circulating cell-free DNA | DNA | D004247 | D | Dna | D004247 | D | 1.00 | 1.00 | 1.00 | 1.00 | 1.00 | 1.00 | 1.00 | 1.00 | 1.00 | 1.00 |
| Immune reponse to smallpox (secreted IL-12p40) | Variola virus | D012901 | B | Smallpox | D012901 | B | 1.00 | 1.00 | 1.00 | 1.00 | 1.00 | 1.00 | 1.00 | 1.00 | 1.00 | 1.00 |
| Smoking cessation | Cessation, Smoking | D016540 | F | Smoking cessation | D016540 | F | 1.00 | 1.00 | 1.00 | 1.00 | 1.00 | 1.00 | 1.00 | 1.00 | 1.00 | 1.00 |
| Alzheimer's disease | Alzheimer Disease | D000544 | C,F | Alzheimer disease | D000544 | F,C | 1.00 | 1.00 | 1.00 | 1.00 | 1.00 | 1.00 | 1.00 | 1.00 | 1.00 | 1.00 |
| Renal cell carcinoma | Carcinoma, Renal Cell | D002292 | C | Carcinoma, renal cell | D002292 | C | 1.00 | 1.00 | 1.00 | 1.00 | 1.00 | 1.00 | 1.00 | 1.00 | 1.00 | 1.00 |
| Alcohol dependence | Alcoholism | D000437 | C,F | Alcoholism | D000437 | F,C | 1.00 | 1.00 | 1.00 | 1.00 | 1.00 | 1.00 | 1.00 | 1.00 | 1.00 | 1.00 |
| Corneal astigmatism | Astigmatism | D001251 | C | Astigmatism | D001251 | C | 1.00 | 1.00 | 1.00 | 1.00 | 1.00 | 1.00 | 1.00 | 1.00 | 1.00 | 1.00 |
| Diabetes (gestational) | Diabetes, Gestational | D016640 | C | Diabetes, gestational | D016640 | C | 1.00 | 1.00 | 1.00 | 1.00 | 1.00 | 1.00 | 1.00 | 1.00 | 1.00 | 1.00 |
| Asthma (childhood onset) | Asthma | D001249 | C | Asthma | D001249 | C | 1.00 | 1.00 | 1.00 | 1.00 | 1.00 | 1.00 | 1.00 | 1.00 | 1.00 | 1.00 |
| Urinary bladder cancer | Urinary Bladder Neoplasms | D001749 | C | Urinary bladder neoplasms | D001749 | C | 1.00 | 1.00 | 1.00 | 1.00 | 1.00 | 1.00 | 1.00 | 1.00 | 1.00 | 1.00 |
| Attention deficit hyperactivity disorder | Attention Deficit Disorder with Hyperactivity | D001289 | F | Attention deficit disorder with hyperactivity | D001289 | F | 1.00 | 1.00 | 1.00 | 1.00 | 1.00 | 1.00 | 1.00 | 1.00 | 1.00 | 1.00 |
| Adiposity | Obesity | D009765 | G,E,C | Adiposity | D009765 | G,E,C | 1.00 | 1.00 | 1.00 | 1.00 | 1.00 | 1.00 | 1.00 | 1.00 | 1.00 | 1.00 |
| Plasma coagulation factors | Blood Coagulation Factors | D001779 | D | Blood coagulation factors | D001779 | D | 1.00 | 1.00 | 1.00 | 1.00 | 1.00 | 1.00 | 1.00 | 1.00 | 1.00 | 1.00 |
| Anticoagulant levels | Anticoagulants | D000925 | D | Anticoagulants | D000925 | D | 1.00 | 1.00 | 1.00 | 1.00 | 1.00 | 1.00 | 1.00 | 1.00 | 1.00 | 1.00 |
| Cholelithiasis-related traits in sickle cell anemia | Cell Trait, Sickle | D012805 | C | Sickle cell trait | D012805 | C | 1.00 | 1.00 | 1.00 | 1.00 | 1.00 | 1.00 | 1.00 | 1.00 | 1.00 | 1.00 |
| Esophageal cancer (alcohol interaction) | Esophageal Neoplasm | D004938 | C | Esophageal neoplasms | D004938 | C | 1.00 | 1.00 | 1.00 | 1.00 | 1.00 | 1.00 | 1.00 | 1.00 | 1.00 | 1.00 |
| Natriuretic peptide levels | Natriuretic Peptides | D045265 | D | Natriuretic peptides | D045265 | D | 1.00 | 1.00 | 1.00 | 1.00 | 1.00 | 1.00 | 1.00 | 1.00 | 1.00 | 1.00 |
| Hepatocellular carcinoma (hepatitis B virus related) | Hepatitis B virus | D006515 | B | Hepatitis b virus | D006515 | B | 1.00 | 1.00 | 1.00 | 1.00 | 1.00 | 1.00 | 1.00 | 1.00 | 1.00 | 1.00 |
| Monocyte early outgrowth colony forming units | Monocytes | D009000 | A | Monocytes | D009000 | A | 1.00 | 1.00 | 1.00 | 1.00 | 1.00 | 1.00 | 1.00 | 1.00 | 1.00 | 1.00 |
| Myocardial infarction | Infarctions, Myocardial | D009203 | C | Myocardial infarction | D009203 | C | 1.00 | 1.00 | 1.00 | 1.00 | 1.00 | 1.00 | 1.00 | 1.00 | 1.00 | 1.00 |
| Thoracic aortic aneurysms and dissections | Thoracic Aortic Aneurysm | D017545 | C | Aortic aneurysm, thoracic | D017545 | C | 1.00 | 1.00 | 1.00 | 1.00 | 1.00 | 1.00 | 1.00 | 1.00 | 1.00 | 1.00 |
| Electroencephalographic traits in alcoholism | Electroencephalography | D004569 | E | Electroencephalography | D004569 | E | 1.00 | 1.00 | 1.00 | 1.00 | 1.00 | 1.00 | 1.00 | 1.00 | 1.00 | 1.00 |
| Response to tamoxifen in breast cancer | Breast Neoplasms | D001943 | C | Breast neoplasms | D001943 | C | 1.00 | 1.00 | 1.00 | 1.00 | 1.00 | 1.00 | 1.00 | 1.00 | 1.00 | 1.00 |
| Ankle-brachial index | Brachial Index, Ankle | D055109 | E | Ankle brachial index | D055109 | E | 1.00 | 1.00 | 1.00 | 1.00 | 1.00 | 1.00 | 1.00 | 1.00 | 1.00 | 1.00 |
| Drinking behavior | Behavior, Drinking | D004327 | F | Drinking behavior | D004327 | F | 1.00 | 1.00 | 1.00 | 1.00 | 1.00 | 1.00 | 1.00 | 1.00 | 1.00 | 1.00 |
| Creutzfeldt-Jakob disease (variant) | Creutzfeldt Jakob Syndrome | D007562 | C,F | Creutzfeldt-jakob syndrome | D007562 | F,C | 1.00 | 1.00 | 1.00 | 1.00 | 1.00 | 1.00 | 1.00 | 1.00 | 1.00 | 1.00 |
| Parkinson`s disease (familial) | Parkinson Disease | D010300 | C | Parkinson disease | D010300 | C | 1.00 | 1.00 | 1.00 | 1.00 | 1.00 | 1.00 | 1.00 | 1.00 | 1.00 | 1.00 |
| Obesity (extreme) | Obesity | D009765 | G,E,C | Obesity | D009765 | G,E,C | 1.00 | 1.00 | 1.00 | 1.00 | 1.00 | 1.00 | 1.00 | 1.00 | 1.00 | 1.00 |
| Cleft lip | Cleft Lip | D002971 | C | Cleft lip | D002971 | C | 1.00 | 1.00 | 1.00 | 1.00 | 1.00 | 1.00 | 1.00 | 1.00 | 1.00 | 1.00 |
| Hypothyroidism | Hypothyroidism | D007037 | C | Hypothyroidism | D007037 | C | 1.00 | 1.00 | 1.00 | 1.00 | 1.00 | 1.00 | 1.00 | 1.00 | 1.00 | 1.00 |
| Prion diseases | Prion Diseases | D017096 | C | Prion diseases | D017096 | C | 1.00 | 1.00 | 1.00 | 1.00 | 1.00 | 1.00 | 1.00 | 1.00 | 1.00 | 1.00 |
| Osteoarthritis | Osteoarthritides | D010003 | C | Osteoarthritis | D010003 | C | 1.00 | 1.00 | 1.00 | 1.00 | 1.00 | 1.00 | 1.00 | 1.00 | 1.00 | 1.00 |
| Hirschsprung`s disease | Disease, Hirschsprung | D006627 | C | Hirschsprung disease | D006627 | C | 1.00 | 1.00 | 1.00 | 1.00 | 1.00 | 1.00 | 1.00 | 1.00 | 1.00 | 1.00 |
| Non-small cell lung cancer | Carcinoma, Non Small Cell Lung | D002289 | C | Carcinoma, non-small-cell lung | D002289 | C | 1.00 | 1.00 | 1.00 | 1.00 | 1.00 | 1.00 | 1.00 | 1.00 | 1.00 | 1.00 |
| Panic disorder | Disorders, Panic | D016584 | F | Panic disorder | D016584 | F | 1.00 | 1.00 | 1.00 | 1.00 | 1.00 | 1.00 | 1.00 | 1.00 | 1.00 | 1.00 |
| Hepatitis B | Hepatitis B | D006509 | C | Hepatitis b | D006509 | C | 1.00 | 1.00 | 1.00 | 1.00 | 1.00 | 1.00 | 1.00 | 1.00 | 1.00 | 1.00 |
| Schizophrenia (treatment refractory) | Schizophrenias | D012559 | F | Schizophrenia | D012559 | F | 1.00 | 1.00 | 1.00 | 1.00 | 1.00 | 1.00 | 1.00 | 1.00 | 1.00 | 1.00 |
| Autism | Autistic Disorder | D001321 | F | Autistic disorder | D001321 | F | 1.00 | 1.00 | 1.00 | 1.00 | 1.00 | 1.00 | 1.00 | 1.00 | 1.00 | 1.00 |
| Major depressive disorder | Depressive Disorders | D003866 | F | Depressive disorder | D003866 | F | 1.00 | 1.00 | 1.00 | 1.00 | 1.00 | 1.00 | 1.00 | 1.00 | 1.00 | 1.00 |
| Cystic fibrosis (meconium ileus) | Cystic Fibrosis | D003550 | C | Cystic fibrosis | D003550 | C | 1.00 | 1.00 | 1.00 | 1.00 | 1.00 | 1.00 | 1.00 | 1.00 | 1.00 | 1.00 |
| Diabetic nephropathy | Diabetic Nephropathies | D003928 | C | Diabetic nephropathies | D003928 | C | 1.00 | 1.00 | 1.00 | 1.00 | 1.00 | 1.00 | 1.00 | 1.00 | 1.00 | 1.00 |
| Lung adenocarcinoma | Lung adenocarcinoma | C538231 | C | Lung neoplasms | D008175 | C | 1.00 | 1.00 | 1.00 | 1.00 | 1.00 | 0.00 | 0.00 | 0.00 | 0.00 | 0.00 |
| Infantile hypertrophic pyloric stenosis | Pyloric Stenosis, Infantile Hypertrophic 1 | C566730 | C | Pyloric stenosis, hypertrophic | D046248 | C | 1.00 | 1.00 | 1.00 | 1.00 | 1.00 | 0.00 | 0.00 | 0.00 | 0.00 | 0.00 |
| D-dimer levels | fibrin fragment D | C036309 | D | Fibrin fibrinogen degradation products | D005338 | D | 1.00 | 1.00 | 1.00 | 1.00 | 1.00 | 0.00 | 0.00 | 0.00 | 0.00 | 0.00 |
| Nasopharyngeal carcinoma | Nasopharyngeal carcinoma | C538339 | C | Nasopharyngeal neoplasms | D009303 | C | 1.00 | 1.00 | 1.00 | 1.00 | 1.00 | 0.00 | 0.00 | 0.00 | 0.00 | 0.00 |
| Esophageal cancer (squamous cell) | Esophageal Squamous Cell Carcinoma | C562729 | C | Esophageal neoplasms | D004938 | C | 1.00 | 1.00 | 1.00 | 1.00 | 1.00 | 0.00 | 0.00 | 0.00 | 0.00 | 0.00 |
| Testicular germ cell tumor | Testicular Germ Cell Tumor | C563236 | C | Neoplasms, germ cell and embryonal | D009373 | C | 1.00 | 1.00 | 1.00 | 1.00 | 1.00 | 0.00 | 0.00 | 0.00 | 0.00 | 0.00 |
| Response to temozolomide | CCRG-81045 | C047246 | D | Dacarbazine | D003606 | D | 1.00 | 1.00 | 1.00 | 1.00 | 1.00 | 0.00 | 0.00 | 0.00 | 0.00 | 0.00 |
| Beta-trace protein levels | prostaglandin endoperoxide D-isomerase | C022466 | D | Intramolecular oxidoreductases | D019746 | D | 1.00 | 1.00 | 1.00 | 1.00 | 1.00 | 0.00 | 0.00 | 0.00 | 0.00 | 0.00 |
| LDL (oxidized) | electronegative low-density lipoprotein | C099252 | D | Lipoproteins, ldl | D008077 | D | 1.00 | 1.00 | 1.00 | 1.00 | 1.00 | 0.00 | 0.00 | 0.00 | 0.00 | 0.00 |
| HbA2 levels | hemoglobin A2' | C049102 | D | Hemoglobin a2 | D006443 | D | 1.00 | 1.00 | 1.00 | 1.00 | 1.00 | 0.00 | 0.00 | 0.00 | 0.00 | 0.00 |
| Non-obstructive azoospermia | Azoospermia, Nonobstructive | C564665 | C | Azoospermia | D053713 | C | 1.00 | 1.00 | 1.00 | 1.00 | 1.00 | 0.00 | 0.00 | 0.00 | 0.00 | 0.00 |
| Lapatinib-induced hepatotoxicity | lapatinib ditosylate | C490728 | D | Quinazolines | D011799 | D | 1.00 | 1.00 | 1.00 | 1.00 | 1.00 | 0.00 | 0.00 | 0.00 | 0.00 | 0.00 |
| Response to clopidogrel therapy | Iscover | C055162 | D | Ticlopidine | D013988 | D | 1.00 | 1.00 | 1.00 | 1.00 | 1.00 | 0.00 | 0.00 | 0.00 | 0.00 | 0.00 |
| Monocyte chemoattractant protein-1 | CCL2, Chemokines | D018932 | D | Monocyte chemoattractant proteins | D018945 | D | 0.96 | 0.49 | 0.96 | 0.96 | 0.87 | 0.98 | 0.53 | 0.98 | 0.98 | 0.30 |
| Meningococcal disease | Meningococcal Infections | D008589 | C | Meningitis, meningococcal | D008585 | C | 0.95 | 0.49 | 0.95 | 0.95 | 0.58 | 0.93 | 0.00 | 0.90 | 0.93 | 0.58 |
| Chronic kidney disease | Chronic Renal Insufficiency | D051436 | C | Kidney failure, chronic | D007676 | C | 0.95 | 0.36 | 0.96 | 0.95 | 0.84 | 0.99 | 0.50 | 0.99 | 0.99 | 0.91 |
| Non-albumin protein levels | Albumins | D000418 | D | Serum albumin | D012709 | D | 0.93 | 0.35 | 0.94 | 0.92 | 0.69 | 0.93 | 0.45 | 0.93 | 0.93 | 0.73 |
| Nonalcoholic fatty liver disease | Non-alcoholic Fatty Liver Disease | D065626 | C | Fatty liver | D005234 | C | 0.92 | 0.44 | 0.92 | 0.92 | 0.79 | 0.00 | 0.00 | 0.00 | 0.00 | 0.82 |
| Hepatitis C induced liver fibrosis | Hepatitis C | D006526 | C | Hepatitis c, chronic | D019698 | C | 0.92 | 0.40 | 0.93 | 0.92 | 0.81 | 0.94 | 0.48 | 0.94 | 0.94 | 0.78 |
| Thiazide-induced adverse metabolic effects in hypertensive patients | Thiazides | D049971 | D | Hydrochlorothiazide | D006852 | D | 0.90 | 0.45 | 0.90 | 0.90 | 0.48 | 0.95 | 0.00 | 0.92 | 0.95 | 0.49 |
| Burning and freckling | Melanoses | D008548 | C | Pigmentation disorders | D010859 | C | 0.90 | 0.40 | 0.91 | 0.90 | 0.75 | 0.88 | 0.49 | 0.86 | 0.88 | 0.73 |
| Freckling | Melanoses | D008548 | C | Pigmentation disorders | D010859 | C | 0.90 | 0.40 | 0.91 | 0.90 | 0.75 | 0.88 | 0.49 | 0.86 | 0.88 | 0.73 |
| Complement C3 and C4 levels | C3 Complement | D003176 | D | Complement system proteins | D003165 | D | 0.89 | 0.40 | 0.90 | 0.89 | 0.73 | 0.88 | 0.00 | 0.88 | 0.88 | 0.81 |
| Weight | Body Weight | D001835 | G,E,C | Obesity | D009765 | G,E,C | 0.89 | 0.29 | 0.93 | 0.88 | 0.78 | 0.92 | 0.43 | 0.93 | 0.92 | 0.14 |
| Epilepsy (generalized) | Epilepsies, Generalized | D004829 | C | Epilepsy | D004827 | C | 0.88 | 0.36 | 0.91 | 0.88 | 0.81 | 0.84 | 0.41 | 0.84 | 0.83 | 0.86 |
| Glioma (high-grade) | Gliomas | D005910 | C | Glioblastoma | D005909 | C | 0.88 | 0.38 | 0.90 | 0.88 | 0.77 | 0.87 | 0.36 | 0.89 | 0.86 | 0.87 |
| Peripartum cardiomyopathy | Cardiomyopathy | D009202 | C | Cardiomyopathy, dilated | D002311 | C | 0.86 | 0.37 | 0.88 | 0.86 | 0.57 | 0.89 | 0.44 | 0.90 | 0.89 | 0.59 |
| Retinopathy in non-diabetics | Diabetic Retinopathies | D003930 | C | Retinal diseases | D012164 | C | 0.83 | 0.34 | 0.86 | 0.83 | 0.41 | 0.81 | 0.39 | 0.81 | 0.80 | 0.32 |
| Chronic kidney disease and serum creatinine concentration | Chronic Renal Insufficiency | D051436 | C | Acute kidney injury | D058186 | C | 0.81 | 0.32 | 0.85 | 0.81 | 0.69 | 0.00 | 0.53 | 0.00 | 0.00 | 0.81 |
| Apolipoprotein Levels | Apolipoproteins | D001053 | D | Apolipoprotein a-ii | D016633 | D | 0.78 | 0.41 | 0.77 | 0.78 | 0.77 | 0.79 | 0.05 | 0.76 | 0.79 | 0.84 |
| Other erythrocyte phenotypes | Erythrocyte | D004912 | A | Erythrocytes, abnormal | D004913 | A | 0.78 | 0.33 | 0.81 | 0.78 | 0.88 | 0.71 | 0.36 | 0.71 | 0.71 | 0.91 |
| End-stage renal disease (non-diabetic) | Kidney Failure, Chronic | D007676 | C | Kidney diseases | D007674 | C | 0.78 | 0.26 | 0.85 | 0.77 | 0.65 | 0.55 | 0.30 | 0.79 | 0.73 | 0.75 |
| Esophageal cancer and gastric cancer | Esophageal Neoplasm | D004938 | C | Gastrointestinal neoplasms | D005770 | C | 0.77 | 0.27 | 0.84 | 0.77 | 0.73 | 0.77 | 0.29 | 0.82 | 0.75 | 0.78 |
| Mean platelet volume | Volumes, Mean Platelet | D063847 | G,E | Platelet count | D010976 | G,E | 0.77 | 0.44 | 0.74 | 0.77 | 0.54 | 0.00 | 0.00 | 0.00 | 0.00 | 0.20 |
| Cutaneous nevi | Nevus, Intradermal | D018330 | C | Nevus | D009508 | C | 0.73 | 0.45 | 0.66 | 0.73 | 0.64 | 0.76 | 0.00 | 0.66 | 0.76 | 0.74 |
| Type 2 diabetes and other traits | Type 2 Diabetes Mellitus | D003924 | C | Hyperinsulinism | D006946 | C | 0.67 | 0.25 | 0.75 | 0.66 | 0.45 | 0.69 | 0.31 | 0.73 | 0.68 | 0.55 |
| Type 1 diabetes nephropathy | Type 1 Diabetes Mellitus | D003922 | C | Diabetic nephropathies | D003928 | C | 0.64 | 0.28 | 0.68 | 0.63 | 0.39 | 0.66 | 0.33 | 0.66 | 0.65 | 0.46 |
| Serum albumin level | Albumin, Serum | D012709 | D | Blood proteins | D001798 | D | 0.63 | 0.19 | 0.78 | 0.61 | 0.69 | 0.62 | 0.24 | 0.71 | 0.60 | 0.73 |
| Dietary macronutrient intake | Diets | D004032 | G | Dietary supplements | D019587 | J,G | 0.52 | 0.19 | 0.66 | 0.51 | 0.33 | 0.00 | 0.00 | 0.00 | 0.00 | 0.45 |
| Circulating vasoactive peptide levels | Peptides | D010455 | D | Vasoactive intestinal peptide | D014660 | D | 0.47 | 0.16 | 0.63 | 0.45 | 0.31 | 0.35 | 0.13 | 0.51 | 0.30 | 0.30 |
| Vascular endothelial growth factor levels | Endothelial Growth Factors | D016228 | D | Vascular endothelial growth factor a | D042461 | D | 0.47 | 0.23 | 0.47 | 0.46 | 0.54 | 0.41 | 0.05 | 0.38 | 0.39 | 0.67 |
| Obesity and blood pressure | Obesity | D009765 | G,E,C | Metabolic syndrome x | D024821 | C | 0.43 | 0.18 | 0.53 | 0.42 | 0.07 | 0.00 | 0.00 | 0.02 | 0.00 | 0.13 |
| Sclerosing cholangitis and ulcerative colitis (combined) | Colitis, Ulcerative | D003093 | C | Bile duct diseases | D001649 | C | 0.38 | 0.16 | 0.49 | 0.36 | 0.10 | 0.00 | 0.00 | 0.03 | 0.00 | 0.17 |
| Cognitive function | Cognition | D003071 | F | Executive function | D056344 | F | 0.35 | 0.13 | 0.50 | 0.31 | 0.54 | 0.00 | 0.00 | 0.00 | 0.00 | 0.58 |
| N-glycan levels | Polysaccharides | D011134 | D | Membrane proteins | D008565 | D | 0.00 | 0.00 | 0.56 | 0.00 | 0.00 | 0.00 | 0.00 | 0.50 | 0.00 | 0.00 |
| Cognitive test performance | Cognition | D003071 | F | Neuropsychological tests | D009483 | F | 0.00 | 0.00 | 0.42 | 0.00 | 0.00 | 0.00 | 0.00 | 0.00 | 0.00 | 0.00 |
| YKL-40 levels | CHI3L1 protein, human | C084533 | D | Biological markers | D015415 | D | 0.00 | 0.00 | 0.37 | 0.00 | 0.00 | 0.00 | 0.00 | 0.00 | 0.00 | 0.00 |
| Plasma carotenoid and tocopherol levels | Carotenoids | D002338 | D | Antioxidants | D000975 | D | 0.00 | 0.00 | 0.23 | 0.00 | 0.00 | 0.00 | 0.00 | 0.06 | 0.00 | 0.00 |
| Cognitive decline | Cognition | D003071 | F | Mild cognitive impairment | D060825 | F | 0.00 | 0.00 | 0.23 | 0.00 | 0.00 | 0.00 | 0.00 | 0.00 | 0.00 | 0.00 |
| Personality dimensions | Personality Disorders | D010554 | F | Personality assessment | D010552 | F | 0.00 | 0.00 | 0.19 | 0.00 | 0.00 | 0.00 | 0.00 | 0.00 | 0.00 | 0.00 |
| Angiotensin-converting enzyme activity | Peptidyl Dipeptidase A | D007703 | D | Angiotensins | D000809 | D | 0.00 | 0.00 | 0.06 | 0.00 | 0.00 | 0.00 | 0.00 | 0.00 | 0.00 | 0.07 |
| F-cell distribution | Fluorides | D005459 | D | Fetal hemoglobin | D005319 | D | 0.00 | 0.00 | 0.00 | 0.00 | 0.00 | 0.00 | 0.00 | 0.00 | 0.00 | 0.00 |
| Serum uric acid | Acid, Uric | D014527 | D | Hyperuricemia | D033461 | C | 0.00 | 0.00 | 0.00 | 0.00 | 0.00 | 0.00 | 0.00 | 0.00 | 0.00 | 0.00 |
| Sphingolipid concentrations | Sphingolipids | D013107 | D | Sphingolipidoses | D013106 | C | 0.00 | 0.00 | 0.00 | 0.00 | 0.00 | 0.00 | 0.00 | 0.00 | 0.00 | 0.00 |
| Eye color traits | Color, Eye | D005127 | G | Color vision defects | D003117 | C | 0.00 | 0.00 | 0.00 | 0.00 | 0.00 | 0.00 | 0.00 | 0.00 | 0.00 | 0.00 |
| Red blood cell traits | Erythrocyte | D004912 | A | Sickle cell trait | D012805 | C | 0.00 | 0.00 | 0.00 | 0.00 | 0.00 | 0.00 | 0.00 | 0.00 | 0.00 | 0.00 |
| Immune response to smallpox vaccine | Smallpox Vaccine | D012900 | D | Smallpox | D012901 | B | 0.00 | 0.00 | 0.00 | 0.00 | 0.00 | 0.00 | 0.00 | 0.00 | 0.00 | 0.00 |
| White blood cell types | Count, Leukocyte | D007958 | G,E | Leukocytes | D007962 | A | 0.00 | 0.00 | 0.00 | 0.00 | 0.00 | 0.00 | 0.00 | 0.00 | 0.00 | 0.00 |
| Red vs. non-red hair color | Color, Hair | D006200 | G | Hair diseases | D006201 | C | 0.00 | 0.00 | 0.00 | 0.00 | 0.00 | 0.00 | 0.00 | 0.00 | 0.00 | 0.00 |
| Skin sensitivity to sun | Skin | D012867 | A | Photophobia | D020795 | C | 0.00 | 0.00 | 0.00 | 0.00 | 0.00 | 0.00 | 0.00 | 0.00 | 0.00 | 0.00 |
| Menopause (age at onset) | Menopause | D008593 | G | Menstruation disturbances | D008599 | C | 0.00 | 0.00 | 0.00 | 0.00 | 0.00 | 0.00 | 0.00 | 0.00 | 0.00 | 0.00 |
| Serum creatinine | Creatinine | D003404 | D | Kidney diseases | D007674 | C | 0.00 | 0.00 | 0.00 | 0.00 | 0.00 | 0.00 | 0.00 | 0.00 | 0.00 | 0.00 |
| Phospholipid levels (plasma) | Plasma | D010949 | A | Personality assessment | D010552 | F | 0.00 | 0.00 | 0.00 | 0.00 | 0.00 | 0.00 | 0.00 | 0.00 | 0.00 | 0.00 |
| Two-hour glucose challenge | Glucose | D005947 | D | Glucose tolerance test | D005951 | E | 0.00 | 0.00 | 0.00 | 0.00 | 0.00 | 0.00 | 0.00 | 0.00 | 0.00 | 0.00 |
| Head circumference (infant) | Infant | D007223 | M | Microcephaly | D008831 | C | 0.00 | 0.00 | 0.00 | 0.00 | 0.00 | 0.00 | 0.00 | 0.00 | 0.00 | 0.00 |
| Vitamin B12 levels | B 12, Vitamin | D014805 | D | Vitamin b 12 deficiency | D014806 | C | 0.00 | 0.00 | 0.00 | 0.00 | 0.00 | 0.00 | 0.00 | 0.00 | 0.00 | 0.00 |
| Tumor biomarkers | Biomarkers, Tumor | D014408 | D | Neoplasms | D009369 | C | 0.00 | 0.00 | 0.00 | 0.00 | 0.00 | 0.00 | 0.00 | 0.00 | 0.00 | 0.00 |
| Urate levels | Acid, Uric | D014527 | D | Hyperuricemia | D033461 | C | 0.00 | 0.00 | 0.00 | 0.00 | 0.00 | 0.00 | 0.00 | 0.00 | 0.00 | 0.00 |
| Age-related macular degeneration (CNV) | DNA Copy Number Variations | D056915 | G | Choroidal neovascularization | D020256 | C | 0.00 | 0.00 | 0.00 | 0.00 | 0.00 | 0.00 | 0.00 | 0.00 | 0.00 | 0.00 |
| Platelet counts | Count, Platelet | D010976 | G,E | Blood platelet disorders | D001791 | C | 0.00 | 0.00 | 0.00 | 0.00 | 0.00 | 0.00 | 0.00 | 0.00 | 0.00 | 0.00 |
| Liver enzyme levels (gamma-glutamyl transferase) | gamma Glutamyltransferase | D005723 | D | Drug-induced liver injury | D056486 | C | 0.00 | 0.00 | 0.00 | 0.00 | 0.00 | 0.00 | 0.00 | 0.00 | 0.00 | 0.00 |
| White matter hypersensitivity burden | White Matters | D066127 | A | Leukoencephalopathies | D056784 | C | 0.00 | 0.00 | 0.00 | 0.00 | 0.00 | 0.00 | 0.00 | 0.00 | 0.00 | 0.00 |
| Vitamin D insufficiency | Ergocalciferols | D004872 | D | Vitamin d deficiency | D014808 | C | 0.00 | 0.00 | 0.00 | 0.00 | 0.00 | 0.00 | 0.00 | 0.00 | 0.00 | 0.00 |
| Plasma levels of liver enzymes | Tests, Clinical Enzyme | D004796 | E | Liver failure | D017093 | C | 0.00 | 0.00 | 0.00 | 0.00 | 0.00 | 0.00 | 0.00 | 0.00 | 0.00 | 0.00 |
| Retinal vascular caliber | Retinal Vessel | D012171 | A | Retinal vasculitis | D031300 | C | 0.00 | 0.00 | 0.00 | 0.00 | 0.00 | 0.00 | 0.00 | 0.00 | 0.00 | 0.00 |
| HDL Cholesterol - Triglycerides (HDLC-TG) | Cholesterol, HDL | D008076 | D | Embolism, cholesterol | D017700 | C | 0.00 | 0.00 | 0.00 | 0.00 | 0.00 | 0.00 | 0.00 | 0.00 | 0.00 | 0.00 |
| Fasting glucose-related traits | Fasting | D005215 | G,F | Glucose metabolism disorders | D044882 | C | 0.00 | 0.00 | 0.00 | 0.00 | 0.00 | 0.00 | 0.00 | 0.00 | 0.00 | 0.00 |
| Fasting plasma glucose | Fasting | D005215 | G,F | Glucose tolerance test | D005951 | E | 0.00 | 0.00 | 0.00 | 0.00 | 0.00 | 0.00 | 0.00 | 0.00 | 0.00 | 0.00 |
| Aortic root size | Root, Tooth | D014092 | A | Aortic aneurysm | D001014 | C | 0.00 | 0.00 | 0.00 | 0.00 | 0.00 | 0.00 | 0.00 | 0.00 | 0.00 | 0.00 |
| Retinol levels | Vitamin A | D014801 | D | Vitamin a deficiency | D014802 | C | 0.00 | 0.00 | 0.00 | 0.00 | 0.00 | 0.00 | 0.00 | 0.00 | 0.00 | 0.00 |
| Liver enzyme levels (alanie transaminase) | Tests, Clinical Enzyme | D004796 | E | Drug-induced liver injury | D056486 | C | 0.00 | 0.00 | 0.00 | 0.00 | 0.00 | 0.00 | 0.00 | 0.00 | 0.00 | 0.00 |
| Menarche (age at onset) | Menarche | D008572 | G | Premenstrual syndrome | D011293 | C | 0.00 | 0.00 | 0.00 | 0.00 | 0.00 | 0.00 | 0.00 | 0.00 | 0.00 | 0.00 |
| Thyroid volume | Gland, Thyroid | D013961 | A | Thyroid diseases | D013959 | C | 0.00 | 0.00 | 0.00 | 0.00 | 0.00 | 0.00 | 0.00 | 0.00 | 0.00 | 0.00 |
| Diastolic blood pressure | Blood Pressure | D001794 | G,E | Hypertension | D006973 | C | 0.00 | 0.00 | 0.00 | 0.00 | 0.00 | 0.00 | 0.00 | 0.00 | 0.00 | 0.00 |
| Vitamin E levels | Vitamin E | D014810 | D | Vitamin e deficiency | D014811 | C | 0.00 | 0.00 | 0.00 | 0.00 | 0.00 | 0.00 | 0.00 | 0.00 | 0.00 | 0.00 |
| Lipoprotein-associated phospholipase A2 activity and mass | 1 Alkyl 2 acetylglycerophosphocholine Esterase | D043203 | D | Hypercholesterolemia | D006937 | C | 0.00 | 0.00 | 0.00 | 0.00 | 0.00 | 0.00 | 0.00 | 0.00 | 0.00 | 0.00 |
| Hematological parameters | Hematology | D006405 | H | Hematologic agents | D006401 | D | 0.00 | 0.00 | 0.00 | 0.00 | 0.00 | 0.00 | 0.00 | 0.00 | 0.00 | 0.00 |
| Serum prostate-specific antigen levels | Prostate-Specific Antigen | D017430 | D | Prostatic neoplasms | D011471 | C | 0.00 | 0.00 | 0.00 | 0.00 | 0.00 | 0.00 | 0.00 | 0.00 | 0.00 | 0.00 |
| Lipid metabolism phenotypes | Lipid Metabolism | D050356 | G | Dyslipidemias | D050171 | C | 0.00 | 0.00 | 0.00 | 0.00 | 0.00 | 0.00 | 0.00 | 0.00 | 0.00 | 0.00 |
| Serum magnesium levels | Magnesium | D008274 | D | Magnesium deficiency | D008275 | C | 0.00 | 0.00 | 0.00 | 0.00 | 0.00 | 0.00 | 0.00 | 0.00 | 0.00 | 0.00 |
| Ventricular conduction | Ventricle, Heart | D006352 | A | Atrioventricular block | D054537 | C | 0.00 | 0.00 | 0.00 | 0.00 | 0.00 | 0.00 | 0.00 | 0.00 | 0.00 | 0.00 |
| Serum metabolites | Serum | D044967 | A | Blood proteins | D001798 | D | 0.00 | 0.00 | 0.00 | 0.00 | 0.00 | 0.00 | 0.00 | 0.00 | 0.00 | 0.00 |
| Vitamin D levels | Ergocalciferols | D004872 | D | Vitamin d deficiency | D014808 | C | 0.00 | 0.00 | 0.00 | 0.00 | 0.00 | 0.00 | 0.00 | 0.00 | 0.00 | 0.00 |
| Hematological and biochemical traits | Hematology | D006405 | H | Hematologic diseases | D006402 | C | 0.00 | 0.00 | 0.00 | 0.00 | 0.00 | 0.00 | 0.00 | 0.00 | 0.00 | 0.00 |
| Serum urate | Acid, Uric | D014527 | D | Urinary calculi | D014545 | C | 0.00 | 0.00 | 0.00 | 0.00 | 0.00 | 0.00 | 0.00 | 0.00 | 0.00 | 0.00 |
| Gamma gluatamyl transferase levels | Gamma Ray | D005720 | G | Gamma-glutamyltransferase | D005723 | D | 0.00 | 0.00 | 0.00 | 0.00 | 0.00 | 0.00 | 0.00 | 0.00 | 0.00 | 0.00 |
| Fasting glucose-related traits (interaction with BMI) | Fasting | D005215 | G,F | Glucose metabolism disorders | D044882 | C | 0.00 | 0.00 | 0.00 | 0.00 | 0.00 | 0.00 | 0.00 | 0.00 | 0.00 | 0.00 |
| Menarche and menopause (age at onset) | Menarche | D008572 | G | Menstruation disturbances | D008599 | C | 0.00 | 0.00 | 0.00 | 0.00 | 0.00 | 0.00 | 0.00 | 0.00 | 0.00 | 0.00 |
| Hair morphology | Hair | D006197 | A | Hair diseases | D006201 | C | 0.00 | 0.00 | 0.00 | 0.00 | 0.00 | 0.00 | 0.00 | 0.00 | 0.00 | 0.00 |
| Triglycerides-Blood Pressure (TG-BP) | Blood Pressure | D001794 | G,E | Metabolic syndrome x | D024821 | C | 0.00 | 0.00 | 0.00 | 0.00 | 0.00 | 0.00 | 0.00 | 0.00 | 0.00 | 0.00 |
| Inflammatory biomarkers | Biomarkers | D015415 | D | Inflammation | D007249 | C | 0.00 | 0.00 | 0.00 | 0.00 | 0.00 | 0.00 | 0.00 | 0.00 | 0.00 | 0.00 |
| Neuranatomic and neurocognitive phenotypes | Phenotype | D010641 | G | Costello syndrome | D056685 | C | 0.00 | 0.00 | 0.00 | 0.00 | 0.00 | 0.00 | 0.00 | 0.00 | 0.00 | 0.00 |
| Sphingolipid levels | Sphingolipids | D013107 | D | Sphingolipidoses | D013106 | C | 0.00 | 0.00 | 0.00 | 0.00 | 0.00 | 0.00 | 0.00 | 0.00 | 0.00 | 0.00 |
| Hematology traits | Hematology | D006405 | H | Hematologic diseases | D006402 | C | 0.00 | 0.00 | 0.00 | 0.00 | 0.00 | 0.00 | 0.00 | 0.00 | 0.00 | 0.00 |
| Hair color | Color, Hair | D006200 | G | Hair diseases | D006201 | C | 0.00 | 0.00 | 0.00 | 0.00 | 0.00 | 0.00 | 0.00 | 0.00 | 0.00 | 0.00 |
| Black vs. blond hair color | Americans, African | D001741 | M | Hair diseases | D006201 | C | 0.00 | 0.00 | 0.00 | 0.00 | 0.00 | 0.00 | 0.00 | 0.00 | 0.00 | 0.00 |
| Eye color | Color, Eye | D005127 | G | Color vision defects | D003117 | C | 0.00 | 0.00 | 0.00 | 0.00 | 0.00 | 0.00 | 0.00 | 0.00 | 0.00 | 0.00 |
| Upper aerodigestive tract cancers | Neoplasms | D009369 | C | Head and neck neoplasms | D006258 | C | 0.00 | 0.00 | 0.00 | 0.00 | 0.00 | 0.00 | 0.00 | 0.00 | 0.00 | 0.50 |
| Breast size | Breast | D001940 | A | Breast diseases | D001941 | C | 0.00 | 0.00 | 0.00 | 0.00 | 0.00 | 0.00 | 0.00 | 0.00 | 0.00 | 0.00 |
| Central corneal thickness | Cornea | D003315 | A | Glaucoma | D005901 | C | 0.00 | 0.00 | 0.00 | 0.00 | 0.00 | 0.00 | 0.00 | 0.00 | 0.00 | 0.00 |
| Platelet aggregation | Aggregation, Platelet | D010974 | G | Thrombosis | D013927 | C | 0.00 | 0.00 | 0.00 | 0.00 | 0.00 | 0.00 | 0.00 | 0.00 | 0.00 | 0.00 |
| Optic disc size (disc) | Disk, Optic | D009898 | A | Glaucoma | D005901 | C | 0.00 | 0.00 | 0.00 | 0.00 | 0.00 | 0.00 | 0.00 | 0.00 | 0.00 | 0.00 |
| Plasma eosinophil count | Eosinophil | D004804 | A | Eosinophilia | D004802 | C | 0.00 | 0.00 | 0.00 | 0.00 | 0.00 | 0.00 | 0.00 | 0.00 | 0.00 | 0.00 |
| Blue vs. green eyes | Eye | D005123 | A | Color vision defects | D003117 | C | 0.00 | 0.00 | 0.00 | 0.00 | 0.00 | 0.00 | 0.00 | 0.00 | 0.00 | 0.00 |
| Blond vs. brown hair color | Color, Hair | D006200 | G | Hair diseases | D006201 | C | 0.00 | 0.00 | 0.00 | 0.00 | 0.00 | 0.00 | 0.00 | 0.00 | 0.00 | 0.00 |
| Black vs. red hair color | Americans, African | D001741 | M | Hair diseases | D006201 | C | 0.00 | 0.00 | 0.00 | 0.00 | 0.00 | 0.00 | 0.00 | 0.00 | 0.00 | 0.00 |
| Serum markers of iron status | Iron, Dietary | D019266 | D | Iron deficiency anemia | D018798 | C | 0.00 | 0.00 | 0.00 | 0.00 | 0.00 | 0.00 | 0.00 | 0.00 | 0.00 | 0.00 |
| Left ventricular mass | Macrophage Activation Syndrome | D055501 | C | Hypertrophy, left ventricular | D017379 | C | 0.00 | 0.00 | 0.00 | 0.00 | 0.00 | 0.00 | 0.00 | 0.00 | 0.00 | 0.00 |
| Response to antipsychotic treatment | Agents, Antipsychotic | D014150 | D | Affective disorders, psychotic | D000341 | F | 0.00 | 0.00 | 0.00 | 0.00 | 0.00 | 0.00 | 0.00 | 0.00 | 0.00 | 0.00 |
| Blue vs. brown eyes | Eye | D005123 | A | Color vision defects | D003117 | C | 0.00 | 0.00 | 0.00 | 0.00 | 0.00 | 0.00 | 0.00 | 0.00 | 0.00 | 0.00 |
| Liver enzyme levels (alkaline phosphatase) | Alkaline Phosphatase | D000469 | D | Drug-induced liver injury | D056486 | C | 0.00 | 0.00 | 0.00 | 0.00 | 0.00 | 0.00 | 0.00 | 0.00 | 0.00 | 0.00 |
| Serum phosphorus concentrations | Phosphorus | D010758 | D | Phosphorus metabolism disorders | D010760 | C | 0.00 | 0.00 | 0.00 | 0.00 | 0.00 | 0.00 | 0.00 | 0.00 | 0.00 | 0.00 |
| Corneal curvature | Cornea | D003315 | A | Corneal wavefront aberration | D057108 | C | 0.00 | 0.00 | 0.00 | 0.00 | 0.00 | 0.00 | 0.00 | 0.00 | 0.00 | 0.00 |
| Serum calcium | Calcium | D002118 | D | Hypocalcemia | D006996 | C | 0.00 | 0.00 | 0.00 | 0.00 | 0.00 | 0.00 | 0.00 | 0.00 | 0.00 | 0.00 |
| Urinary metabolites | Tract, Urinary | D014551 | A | Renal tubular transport, inborn errors | D015499 | C | 0.00 | 0.00 | 0.00 | 0.00 | 0.00 | 0.00 | 0.00 | 0.00 | 0.00 | 0.00 |
| Hemostatic factors and hematological phenotypes | Bothrops asper antihemorrhagic factor | C003941 | D | Hemostatic disorders | D020141 | C | 0.00 | 0.00 | 0.00 | 0.00 | 0.00 | 0.00 | 0.00 | 0.00 | 0.00 | 0.00 |
| Serum iron concentration | Attention | D001288 | F | Anemia | D000740 | C | 0.00 | 0.00 | 0.00 | 0.00 | 0.00 | 0.00 | 0.00 | 0.00 | 0.00 | 0.00 |
| Serum hepcidin | Hepcidins | D064451 | D | Anemia, iron-deficiency | D018798 | C | 0.00 | 0.00 | 0.00 | 0.00 | 0.00 | 0.00 | 0.00 | 0.00 | 0.00 | 0.00 |
| Plasma homocysteine | Homocystine | D006711 | D | Homocystinuria | D006712 | C | 0.00 | 0.00 | 0.00 | 0.00 | 0.00 | 0.00 | 0.00 | 0.00 | 0.00 | 0.00 |
| Urinary albumin excretion | Albumins | D000418 | D | Albuminuria | D000419 | C | 0.00 | 0.00 | 0.00 | 0.00 | 0.00 | 0.00 | 0.00 | 0.00 | 0.00 | 0.00 |
| Red vs non-red hair color | Color, Hair | D006200 | G | Hair diseases | D006201 | C | 0.00 | 0.00 | 0.00 | 0.00 | 0.00 | 0.00 | 0.00 | 0.00 | 0.00 | 0.00 |
| Immunoglobulin A | Immunoglobulin A | D007070 | D | Iga deficiency | D017098 | C | 0.00 | 0.00 | 0.00 | 0.00 | 0.00 | 0.00 | 0.00 | 0.00 | 0.00 | 0.00 |
| Neutrophil count | Neutrophil | D009504 | A | Neutropenia | D009503 | C | 0.00 | 0.00 | 0.00 | 0.00 | 0.00 | 0.00 | 0.00 | 0.00 | 0.00 | 0.00 |
| IgE grass sensitization | Poaceae | D006109 | B | Hypersensitivity, immediate | D006969 | C | 0.00 | 0.00 | 0.00 | 0.00 | 0.00 | 0.00 | 0.00 | 0.00 | 0.00 | 0.00 |
| Serum IgE levels | Serum | D044967 | A | Hypersensitivity, immediate | D006969 | C | 0.00 | 0.00 | 0.00 | 0.00 | 0.00 | 0.00 | 0.00 | 0.00 | 0.00 | 0.00 |
| Cortical structure | Cerebral Cortex | D002540 | A | Malformations of cortical development | D054220 | C | 0.00 | 0.00 | 0.00 | 0.00 | 0.00 | 0.00 | 0.00 | 0.00 | 0.00 | 0.00 |
| Caffeine consumption | Caffeine | D002110 | D | Psychoses, substance-induced | D011605 | F,C | 0.00 | 0.00 | 0.00 | 0.00 | 0.00 | 0.00 | 0.00 | 0.00 | 0.00 | 0.00 |
| Renal function-related traits (urea) | Urea | D014508 | D | Acute kidney injury | D058186 | C | 0.00 | 0.00 | 0.00 | 0.00 | 0.00 | 0.00 | 0.00 | 0.00 | 0.00 | 0.00 |
| Serum bilirubin levels | Bilirubin | D001663 | D | Hyperbilirubinemia | D006932 | C | 0.00 | 0.00 | 0.00 | 0.00 | 0.00 | 0.00 | 0.00 | 0.00 | 0.00 | 0.00 |
| Aging (facial) | Aging | D000375 | G | Skin aging | D015595 | G | 0.00 | 0.00 | 0.00 | 0.00 | 0.00 | 0.00 | 0.00 | 0.00 | 0.00 | 0.00 |
| Fasting insulin-related traits | Insulin | D007328 | D | Metabolism, inborn errors | D008661 | C | 0.00 | 0.00 | 0.00 | 0.00 | 0.00 | 0.00 | 0.00 | 0.00 | 0.00 | 0.00 |
| Folate pathway vitamin levels | Vitamins | D014815 | J,G,D | Folic acid deficiency | D005494 | C | 0.00 | 0.00 | 0.00 | 0.00 | 0.00 | 0.00 | 0.00 | 0.00 | 0.00 | 0.00 |
| Plasma level of vitamin B12 | B 12, Vitamin | D014805 | D | Vitamin b12 deficiency | D014806 | C | 0.00 | 0.00 | 0.00 | 0.00 | 0.00 | 0.00 | 0.00 | 0.00 | 0.00 | 0.00 |
| Serum soluble E-selectin | E-Selectin | D019040 | D | Inflammation | D007249 | C | 0.00 | 0.00 | 0.00 | 0.00 | 0.00 | 0.00 | 0.00 | 0.00 | 0.00 | 0.00 |
| Plasma levels of Protein C | Blood Proteins | D001798 | D | Protein c deficiency | D020151 | C | 0.00 | 0.00 | 0.00 | 0.00 | 0.00 | 0.00 | 0.00 | 0.00 | 0.00 | 0.00 |
| Bitter taste response | Taste | D013649 | G,F | Taste disorders | D013651 | C | 0.00 | 0.00 | 0.00 | 0.00 | 0.00 | 0.00 | 0.00 | 0.00 | 0.00 | 0.00 |
| Cardiac repolarization | Heart | D006321 | A | Arrhythmias, cardiac | D001145 | C | 0.00 | 0.00 | 0.00 | 0.00 | 0.00 | 0.00 | 0.00 | 0.00 | 0.00 | 0.00 |
| Facial morphology | Face | D005145 | A | Facial asymmetry | D005146 | C | 0.00 | 0.00 | 0.00 | 0.00 | 0.00 | 0.00 | 0.00 | 0.00 | 0.00 | 0.00 |
| Plasma Lp (a) levels | Lipoprotein(a) | D017270 | D | Hyperlipoproteinemias | D006951 | C | 0.00 | 0.00 | 0.00 | 0.00 | 0.00 | 0.00 | 0.00 | 0.00 | 0.00 | 0.00 |
| Body mass (lean) | Bodies, Human | D018594 | K,I | Thinness | D013851 | G,E,C | 0.00 | 0.00 | 0.00 | 0.00 | 0.00 | 0.00 | 0.00 | 0.00 | 0.00 | 0.00 |
| Immune response to smallpox vaccine (IL-6) | Interleukin 6 | D015850 | D | Smallpox | D012901 | B | 0.00 | 0.00 | 0.00 | 0.00 | 0.00 | 0.00 | 0.00 | 0.00 | 0.00 | 0.00 |
| Coronary artery calcification | Calcification, Physiologic | D002113 | G | Vascular calcification | D061205 | C | 0.00 | 0.00 | 0.00 | 0.00 | 0.00 | 0.00 | 0.00 | 0.00 | 0.00 | 0.00 |
| Cortical thickness | Cerebral Cortex | D002540 | A | Cobblestone lissencephaly | D054222 | C | 0.00 | 0.00 | 0.00 | 0.00 | 0.00 | 0.00 | 0.00 | 0.00 | 0.00 | 0.00 |
| IFN-related cytopenia | Families | D005190 | I,F | Thrombocytopenia | D013921 | C | 0.00 | 0.00 | 0.00 | 0.00 | 0.00 | 0.00 | 0.00 | 0.00 | 0.00 | 0.00 |
| Lymphocyte counts | Lymphocyte Count | D018655 | G,E | Lymphocytosis | D008218 | C | 0.00 | 0.00 | 0.00 | 0.00 | 0.00 | 0.00 | 0.00 | 0.00 | 0.00 | 0.00 |
| Gamma gluatamyl transferase levels (interaction with age) | Gamma Ray | D005720 | G | Gamma-glutamyltransferase | D005723 | D | 0.00 | 0.00 | 0.00 | 0.00 | 0.00 | 0.00 | 0.00 | 0.00 | 0.00 | 0.00 |
| Intraocular pressure | Ocular Tonometry | D014065 | E | Ocular hypertension | D009798 | C | 0.00 | 0.00 | 0.00 | 0.00 | 0.00 | 0.00 | 0.00 | 0.00 | 0.00 | 0.00 |
| Pulmonary function | Lungs | D008168 | A | Lung diseases | D008171 | C | 0.00 | 0.00 | 0.00 | 0.00 | 0.00 | 0.00 | 0.00 | 0.00 | 0.00 | 0.00 |
| Renal function-related traits (BUN) | Kidney | D007668 | A | Acute kidney injury | D058186 | C | 0.00 | 0.00 | 0.00 | 0.00 | 0.00 | 0.00 | 0.00 | 0.00 | 0.00 | 0.00 |
| Renal function-related traits (eGRFcrea) | Kidney | D007668 | A | Acute kidney injury | D058186 | C | 0.00 | 0.00 | 0.00 | 0.00 | 0.00 | 0.00 | 0.00 | 0.00 | 0.00 | 0.00 |
| Renal function-related traits (sCR) | Kidney | D007668 | A | Acute kidney injury | D058186 | C | 0.00 | 0.00 | 0.00 | 0.00 | 0.00 | 0.00 | 0.00 | 0.00 | 0.00 | 0.00 |
| Pulmonary function (interaction) | Lungs | D008168 | A | Respiratory function tests | D012129 | E | 0.00 | 0.00 | 0.00 | 0.00 | 0.00 | 0.00 | 0.00 | 0.00 | 0.00 | 0.00 |
| Pulmonary function measures | Lungs | D008168 | A | Lung diseases | D008171 | C | 0.00 | 0.00 | 0.00 | 0.00 | 0.00 | 0.00 | 0.00 | 0.00 | 0.00 | 0.00 |
| Thyroid function | Gland, Thyroid | D013961 | A | Thyroid diseases | D013959 | C | 0.00 | 0.00 | 0.00 | 0.00 | 0.00 | 0.00 | 0.00 | 0.00 | 0.00 | 0.00 |
| Blood lipid traits | Blood | D001769 | A | Hyperlipidemias | D006949 | C | 0.00 | 0.00 | 0.00 | 0.00 | 0.00 | 0.00 | 0.00 | 0.00 | 0.00 | 0.00 |
| Cardiac structure and function | Heart | D006321 | A | Heart diseases | D006331 | C | 0.00 | 0.00 | 0.00 | 0.00 | 0.00 | 0.00 | 0.00 | 0.00 | 0.00 | 0.00 |
| Response to Vitamin E supplementation | Vitamin E | D014810 | D | Vitamin e deficiency | D014811 | C | 0.00 | 0.00 | 0.00 | 0.00 | 0.00 | 0.00 | 0.00 | 0.00 | 0.00 | 0.00 |
